# *α*-Aminooxyacetic acid derivatives acting as pro-drugs against *Mycobacterium tuberculosis*

**DOI:** 10.1101/2024.04.04.587628

**Authors:** Kristin Vill, Lasse van Geelen, Oliver Michel, Anna-Lene Kiffe-Delf, Alexander Berger, David Podlesainski, Katharina Stenzel, Filip Kovacic, Beate Lungerich, Björn Burkhardt, Taylor A. Crooks, Michael D. Howe, Lev Ostrer, Ziyi Jia, Thomas R. Ioerger, Farnusch Kaschani, Markus Kaiser, Anthony D. Baughn, Thomas Kurz, Rainer Kalscheuer

## Abstract

Tuberculosis (TB), a significant cause of mortality globally, continues to claim 1.5 million lives each year. Despite recent advances in TB management, the emergence of multidrug-resistant strains of TB is exacerbating the treatment of TB. Therefore, there is an immediate necessity to uncover new anti-TB compounds with unprecedented targets. This study introduces novel antimycobacterial molecules that are based on α-aminooxyacetic acid core structures. The lead compounds KSK-104 and KSK-106 displayed potent sub-micromolar antibacterial activity against *Mycobacterium tuberculosis* H37Rv and XDR clinical isolates, while exhibiting virtually no cytotoxicity against various human cells. Complementation experiments following whole genome sequencing of spontaneously resistant mutants generated against these bactericidal compounds suggested that they are pro-drugs that are intracellularly hydrolyzed by one or both of two specific amidohydrolases, Rv0552 and AmiC. Furthermore, proteomic and transcriptomic analyses of stressed cells and genetic interaction mapping employing transposon insertion sequencing suggest a “dirty drug” mechanism that involves the simultaneous attack of the various drug cleavage products on multiple intracellular targets. Our results suggest a primary role of the pyridoxal 5’-phosphate (PLP) synthesis and salvage pathway and/or PLP-dependent enzymes, the oxidative stress network, and the largely uncharacterized *Rv3092c-Rv3095* gene cluster in the mode of action.

## INTRODUCTION

Tuberculosis (TB) is among the oldest illnesses affecting humanity. Caused by the pathogen *Mycobacterium tuberculosis*, it continues to pose a global health concern, accounting for 1.6 million deaths in 2021.^1^ The drugs currently utilized as front-line chemotherapy (isoniazid, rifampicin, pyrazinamide, and ethambutol) were developed over half a century ago with clinical trials determining their optimal combination and duration being mainly conducted in the 1970s.^2^ Although the standard treatment is safe and well-tolerated, treatment of drug-sensitive tuberculosis requires a combination therapy for six months, which frequently leads to patients’ non-adherence and the emergence of drug resistance. The current situation appears grim due to the spread of multidrug (MDR) and extensively drug-resistant (XDR) strains, the fatal combination of TB with HIV/AIDS infection, and the recent setback caused by the coronavirus disease (COVID-19) pandemic, which has led to inadequate TB diagnoses and treatments.^1,3^ Recent advances in the development of drugs against drug-resistant TB led to the regulatory approval of bedaquiline, delamanid, and pretomanid, as well as the entry into clinical trials of several new compounds.^4^ Furthermore, the development of regimens that shorten treatment is a promising and active research area.^5^ While this offers hope for patients, there remains an urgent need to develop novel promising treatments addressing the drug-resistant TB crisis in the future, considering that drug development is a lengthy and risky process. Yet, the development of faster-acting TB drugs with a low risk of resistance remains a challenge, mainly due to the complex biology of the pathogen and the extraordinary pathology of the disease itself. To this end, it is essential to gain a deep understanding of the mode of action of novel compounds and the strategies employed by *M. tuberculosis* to resist their antibacterial effect.

In recent years, high-throughput small molecule library screening campaigns employing *in vitro* cultured *M. tuberculosis* has led to the discovery of several novel growth inhibitors from previously unexplored chemical classes, highlighting the untapped potential of molecules that were previously not envisioned to exhibit activity against the pathogen. Following the same approach by screening of an in-house compound library of the research group Kurz, we identified α-aminooxyacetic acid derivatives as promising novel anti-TB lead compounds. The front-runner compounds, KSK-104 and KSK-106, are *para*-substituted benzoylated derivatives of 2-aminoxy-*N*-(benzyloxy)acetamide. A 1971 patent by Kisfaludy *et al.* first described α-aminooxyhydroxamic acid derivatives as compounds with antimycobacterial activity.^6^ However, to the best of our knowledge, no data on their mechanism of action or molecular target have been published, and the development was discontinued for unknown reasons after two follow-up studies in the 1970s. *para*-Substituted benzoylated derivatives of 2-aminoxy-*N*-(benzyloxy)acetamide such as KSK-104 and KSK-106, which differ from the earlier described compounds, have not been previously reported, nor have their antitubercular activities been evaluated in detail.

Here, we report on the synthesis and anti-tubercular properties of these novel alkoxyamide lead structures. We furthermore provide evidence suggesting that they represent pro-drugs that are activated following intracellular hydrolysis by the putative amidohydrolases AmiC and Rv0552, releasing different active metabolites that act as “dirty drugs” pleiotropically targeting different intracellular pathways, including pyridoxal 5’-phosphate (PLP) synthesis and salvage and/or PLP-dependent pathways, the oxidative stress network as well as the yet uncharacterized *Rv3092c-Rv3095* operon.

## RESULTS

### Novel α-aminooxyacetic acid molecules as potent anti-TB lead structures

The potential of α-aminooxyacetic acid molecules to act as antimycobacterials, has been described back in 1971 by Kisfaludy *et al.*.^6^ However, although some follow-up studies indicated promising *in vivo* and *in vivo* activity, further reports on the characterization of the molecules for the development of new anti-tuberculosis agents have been missing.^6^ It therefore remained unclear how α-aminooxyacetic acid derivatives impact *M. tuberculosis* physiology and growth. During the screening of an in-house α-aminooxyacetic acid compound library, we found two novel compounds, KSK-104 and KSK-106 (subsequently collectively referred to as KSKs) exhibiting potent antibacterial *in-vitro* growth inhibitory activity against cells of the laboratory strain *M. tuberculosis* H37Rv. The structure of both molecules consists of three distinct regions: region B with an aminooxyacetyl backbone, region C with a benzyloxyamine group, and variable region A, occupied by a *para*-phenyl substituted benzoyl group in KSK-104, or a *para*-pentoxy substituted benzoyl group in case of KSK-106 (Figure 1A).

**Figure 1.**
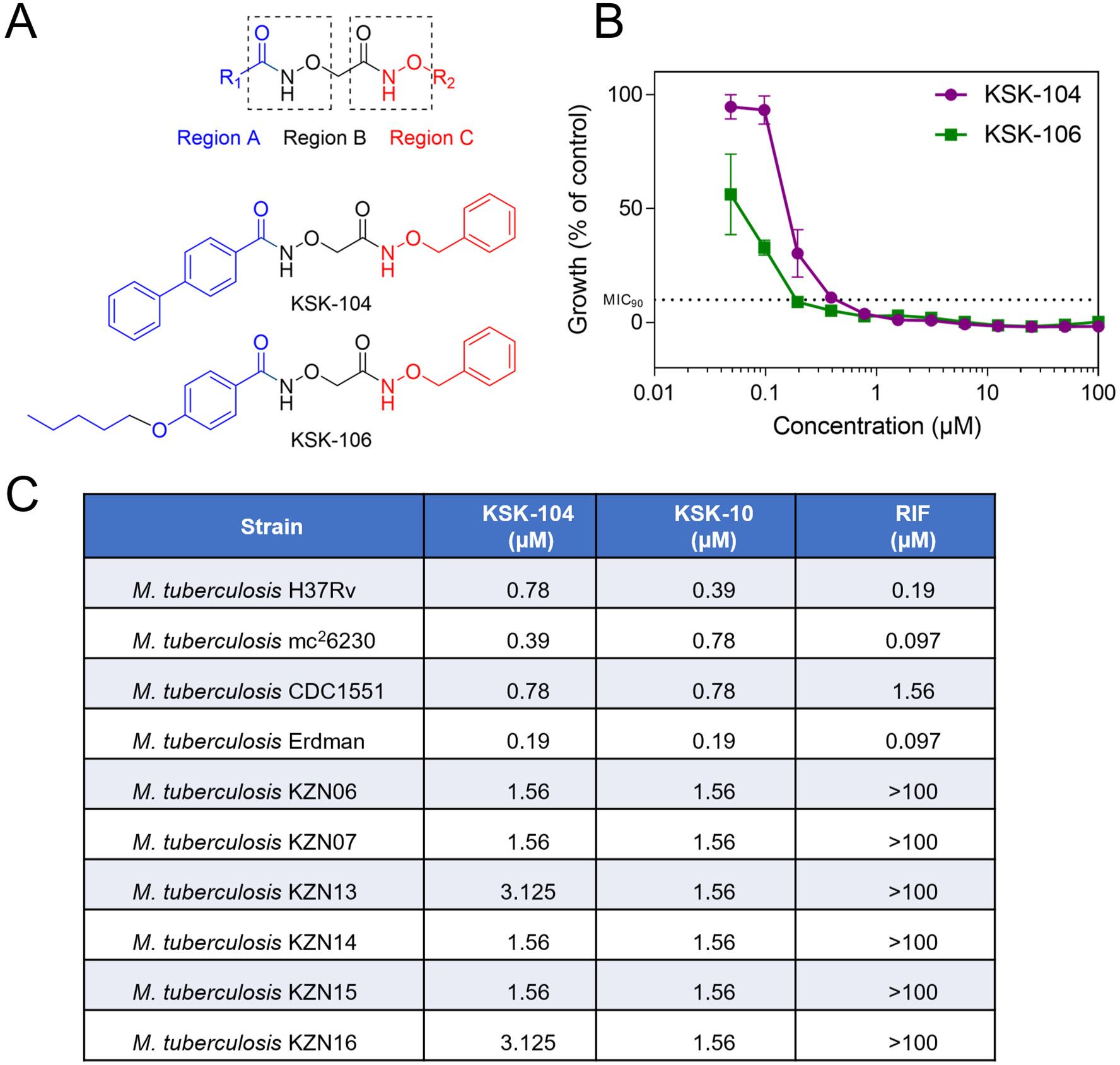
Structures of KSK molecules and antibacterial activity against *M. tuberculosis*. **A)** Chemical structures of the α-aminooxyacetic derivatives KSK-104 and KSK-106 indicating three regions. The alkoxyamide and benzyloxamide moieties/structures are boxed. **B)** Dose-response curves for KSK-104 and KSK-106 demonstrating a concentration-dependent inhibition of *M. tuberculosis* H37Rv growth. Data are shown as means of triplicates with SD. **C)** MIC_90_ values of rifampicin (RIF), KSK-104 and KSK-106 for various *M. tuberculosis* laboratory strains and for clinical XDR isolates (KZN) originating from the KwaZulu-Natal region, South Africa. Growth was quantified by employing the resazurin reduction assay. Data represent a single experiment with *n* = 3 with no variations in MIC_90_ values observed between individual samples.

We found that both KSKs show sub-micromolar minimum inhibitory concentrations for inhibiting at least 90% of growth compared to respective solvent controls (MIC_90_) with KSK-104 having an MIC_90_ of 0.78 μM and KSK-106 of 0.39 μM (Figure 1B). Importantly, extensively drug-resistant (XDR) clinical isolates of *M. tuberculosis* originating from the KwaZulu-Natal region in South Africa were also susceptible to the KSKs with MIC_90_ values ranging from 0.78 to 3.125 μM (Figure 1C). This indicates that the KSKs do not share a similar mechanism of action compared to the clinically used drugs, to which the tested *M. tuberculosis* XDR strains are resistant.

Next, we established a flexible straightforward synthesis route for KSK-104 and KSK-106 to obtain sufficient amounts of these novel anti-TB lead structures for pharmacological tests and to enable their optimization through chemical derivatization (Scheme 1).

**Scheme 1.**
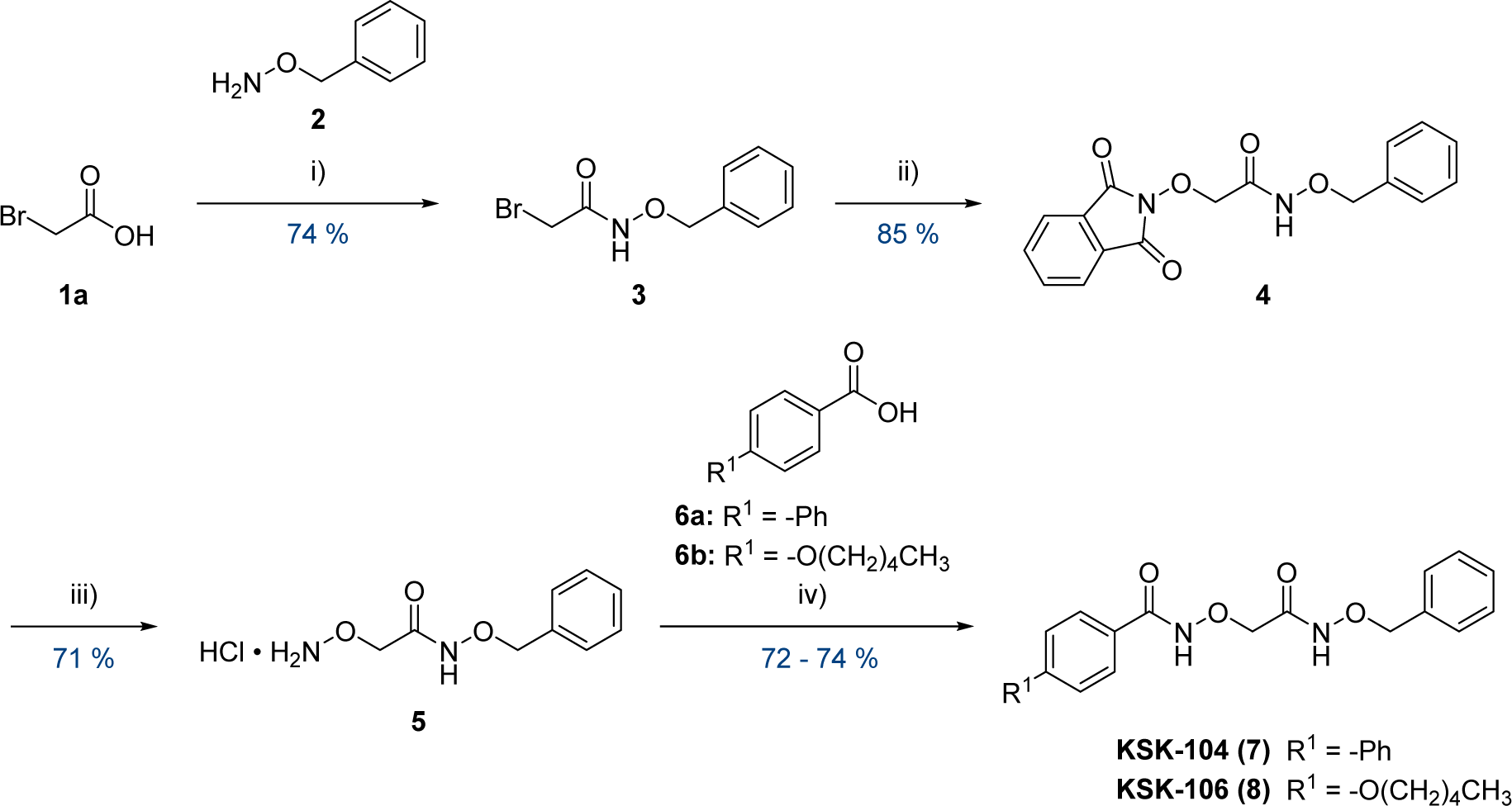
Synthesis of novel anti-TB lead structures KSK-104 and KSK-106. i) 1.00 eq. IBCF, 1.00 eq. NMM, THF, −20 °C to rt, 16 h; ii) 1.15 eq. NHPI, 1.15 eq. NEt_3_, MeCN, reflux, 4 h; iii) 4.00 eq. methylhydrazine, 4.00 eq. HCL in dioxane (4 M), CH_2_Cl_2_, −10 °C, 16 h; iv) 1.20 eq. EDC·HCl, 1.20 eq. NEt_3_, 0.10 eq. DMAP, CH_2_Cl_2_, rt, 16 h.

The protected intermediate **3** was synthesized via a coupling reaction mediated by isobutylchloroformiate (IBCF) and *N*-methylmorpholine (NMM) between bromoacetic acid (**1a**) and *O*-benzylhydroxylamine (**2**), using a mixed anhydride as the acylating intermediate. The subsequent alkylation of *N*-hydroxyphthalimide (NHPI) with **3** yielded the phthaloyl-protected hydroxylamine **4**. Deprotection of **4** was achieved through methylhydrazinolysis, and the resulting hydroxylamine **5** was isolated as hydrochloride. Finally, EDC-mediated coupling reactions were performed for the acylation of **5**, using either 4-phenylbenzoic acid (**6a**) or 4-pentyloxybenzoic acid (**6b**), to produce KSK-104 (**7**) with a yield of 72% or KSK-106 (**8**) with a yield of 74%, respectively.

The stability of both lead structures, KSK-104 and KSK-106, in aqueous media or human EDTA-plasma was tested at 37 °C to evaluate their suitability for further drug development. In aqueous media, both lead structures exhibited a degradation of approximately 3% over 48 hours at pH 2, whereas after 48 hours at pH 7.4 only approximately 1.5% of compounds were degraded (Figure S1, Supporting Information). In human EDTA plasma, KSK-106 showed high stability with no measurable degradation after 6 hours, and only 20% degradation after 24 hours, giving a calculated half-life of 77.9 hours ex vivo. No degradation of KSK-104 in EDTA-plasma could be detected after 24 h (Figure S2). These results indicate their suitable chemical stability in conditions mimicking physiologically relevant conditions, thereby emphasizing their suitability for further development.

### KSK-104 and KSK-106 are bactericidal and selectively active against tuberculous mycobacteria

We continued our investigations of the KSKs by characterizing their anti-tubercular and cytotoxicity profiles. We found that KSK-104 and KSK-106 were specifically active against *M. tuberculosis* and *M. bovis* BCG Pasteur, while no growth inhibition of other tested fast-growing mycobacteria such as *Mycobacterium marinum*, *Mycobacterium abscessus* and *Mycobacterium smegmatis* was observed. These results indicate that the compounds are only effective against tuberculous mycobacteria (Figure S3A). Additionally, the compounds have been tested against several nosocomial bacteria and fungi, such as *Staphylococcus aureus* Mu50, *Acinetobacter baumannii* ATCC BAA-1605, *Pseudomonas aeruginosa* ATCC 27853, and *Candida albicans* ATCC 24433 (Figure S3A). Growth of none of these pathogens was affected by the treatment with KSKs suggesting a specific tuberculous mechanism and/or target of the novel compounds. Of note, KSK-104, and KSK-106 showed no cytotoxic effects on various human cell lines originating from different tissues at concentrations up to 100 μM (Figure S3B).

We then monitored the anti-TB effects of both molecules over 35 days performing a time-killing kinetic to investigate how KSK-104 and KSK-106 exert their growth inhibition on *M. tuberculosis*. Both KSK-104 and KSK-106 exhibited a bactericidal effect within the first 9 days of incubation, resulting in a 2 - 3 −log_10_ reduction in viable cell counts as determined by quantifying colony forming units (CFU) (Figure 2A). No further reduction in viability occurred after day 9; in contrast to the tested clinical drugs, however, CFU counts remained constant and no resumption of growth was observed in monotreatment (Figure S4). Since the standard treatment of TB requires a combination therapy employing four different drugs,^7,8^ we tested KSK-104 and KSK-106 in combination with the first-line antibiotics isoniazid, rifampicin, and ethambutol as well as with delamanid and bedaquiline (Figure S4). Both KSK compounds showed an additive effect in combination with isoniazid, rifampicin, ethambutol, and delamanid, indicated by a reduction of viable cell counts to the detection limit of 10^2^ CFU/mL. Furthermore, particularly the combination with KSK-106 efficiently suppresseded the resurgence of surviving bacteria even after 35 days of incubation. In contrast, bedaquiline substantially dampened the bactericidal effect of KSK compounds, indicating antagonistic drug-drug interference.

**Figure 2.**
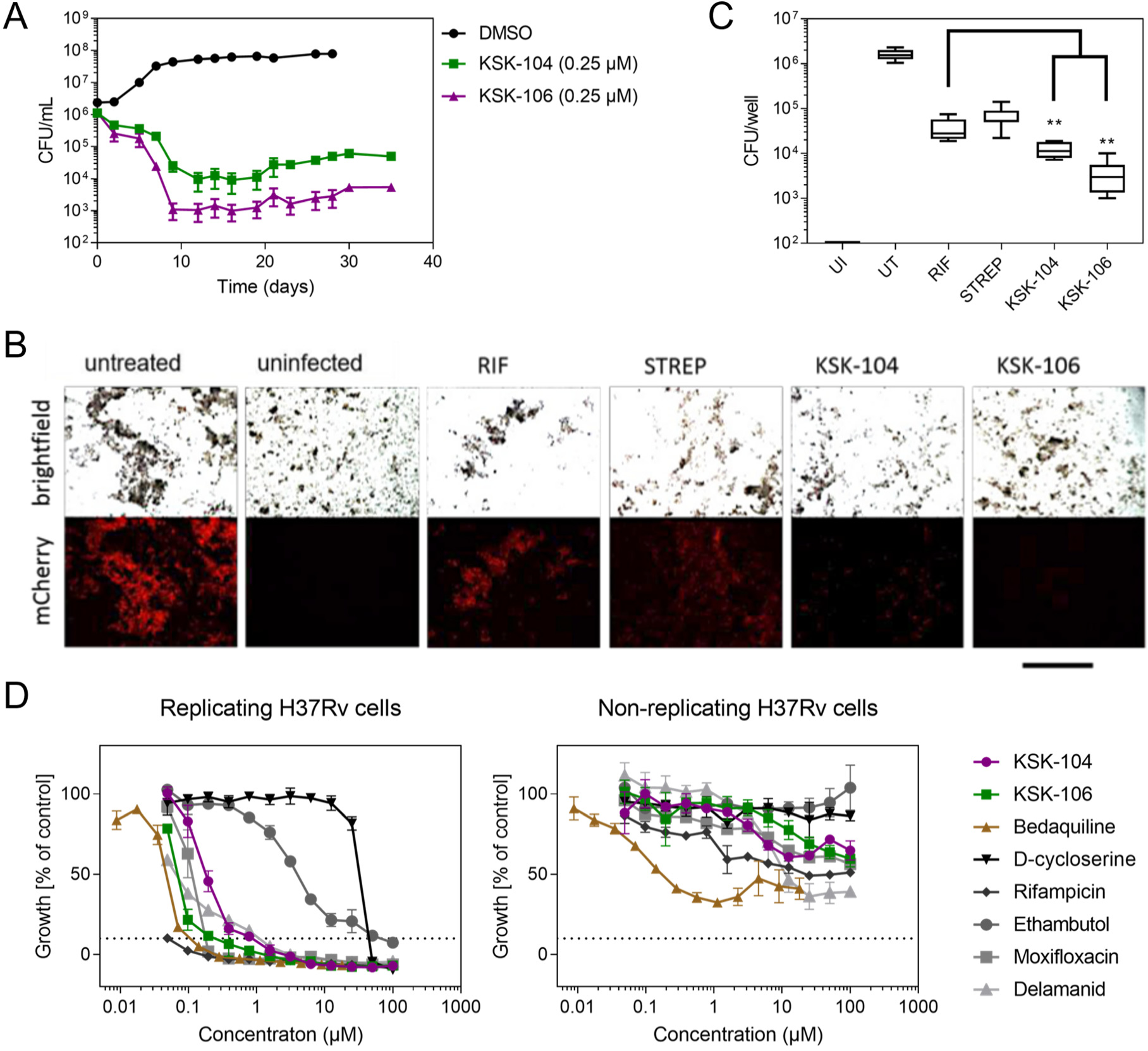
KSKs exhibit a sub-micromolar bactericidal anti-TB activity in vitro and in vivo using a THP1 infection model. **A)** Killing kinetic study by quantifying viable cell count (CFU) following different treatment intervals. *M. tuberculosis* H37Rv cultures treated with KSK-104 or KSK-106. KSK-104 and KSK-106 showed a bactericidal effect. Experiments have been performed in triplicates. Data are shown as mean ± S.D.. **B)** Effects of KSKs on intracellular growth of a constitutively mCherry-expressing *M. tuberculosis* H37Rv reporter strain in a human macrophage THP-1 infection model. THP-1-derived macrophages were treated three hours after infection with either 20 µM DMSO, 0.5 μM KSK-104 or KSK-106, or with the clinical anti-TB drugs rifampicin (RIF, 3 μM) or streptomycin (STREP, 20 μM). After 5 days, fluorescence microscopy was used to visualize internalized bacteria. Representative brightfield pictures of macrophages, and the corresponding fluorescence micrographs are shown. The bar represents 10 µm. **C)** Box-plot of CFU counting after infection of THP-1 macrophages with *M. tuberculosis* H37Rv. Cells were treated with antibiotic and KSK concentrations as described above. UI = uninfected, UT = untreated (DMSO), RIF = rifampicin, STREP = streptomycin. Boxes show upper and lower quartiles with median. Statistical significance (** *p* < 0.01) was calculated using Kolmogorov-Smirnov-test and student’s t-test. All experiments have been performed in triplicates and have been repeated once with similar results. **D)** The activity of KSK-104 and KSK-106 was investigated against replicating and non-replicating *M. tuberculosis* H37Rv cells compared to the several first- and second-line drugs. To generate starvation-induced non-replicating bacilli, cultures of *M. tuberculosis* H37Rv cells were washed, suspended in PBS, and incubated at 37°C for three weeks. Starved cultures were then incubated for 7 days with the drugs. Moderate efficacy against starvation-induced non-replicating cells was observed only for bedaquiline. Growth was quantified by employing the resazurin reduction assay. KSKs exhibited a low activity, comparable to rifampicin, moxifloxacin, and delamanid, while ethambutol and D-cycloserine did not show any activity. Data shown as means of triplicates with SD. The dashed lines indicate 10% residual growth.

*M. tuberculosis* is an intracellular pathogen, mainly residing and replicating in phagolysosomes of macrophages thereby escaping the immune response mechanisms of the infected host.^9,10^ To evaluate whether the KSKs also interfere with intracellular growth, we employed a THP-1 human macrophage infection model that relies on quantifying cell growth of a mCherry-expressing fluorescent *M. tuberculosis* reporter strain. While macrophages treated with the solvent control (DMSO) exhibited a high intracellular bacterial burden, both KSK compounds substantially inhibited intracellular proliferation of *M. tuberculosis* and resulted in a healthy morphology of the treated macrophages (Figure 2B). Remarkably, treatment of infected THP-1 cells with either KSK-104 or KSK-106 at 0.5 µM resulted in bacterial fluorescence lower than the rifampicin (3 µM) and streptomycin (20 µM) controls. These results indicate a stronger potency of both KSKs in killing intracellular *M. tuberculosis* residing in human macrophages (Figure 2B). These results were confirmed by counting CFU after plating bacteria isolated from THP-1 cells infected with the non-fluorescent parental *M. tuberculosis* H37Rv strain (Figure 2C). Conclusively, KSKs can enter relevant human target immune cells and inhibit *M. tuberculosis* growth with higher efficiency than the tested first-line drugs.

The duration of TB therapy is lengthy, in part because current anti-TB drugs are mainly effective against actively growing, replicating mycobacteria, but considerably less active against non-replicating bacteria.^11^ Therefore, drugs that inhibit these non-replicating persisting *M. tuberculosis* cells that are known to be highly tolerant against a multitude of clinical drugsare desired.^12^ We thus evaluated the activity of the KSKs against both replicating and starvation-induced non-replicating cells of *M. tuberculosis* H37Rv in comparison to the first- and second-line drugs bedaquiline, D-cycloserine, rifampicin, ethambutol, moxifloxacin and delamanid (Figure 2D). While replicating cells were sensitive to all tested compounds including the KSKs, the starvation-induced non-replicating cells were highly tolerant to all tested antimycobacterials, with none of them completely inhibiting growth even at the highest tested concentration. While bedaquiline showed highest potency, KSK-104 and KSK-106 demonstrated at least some activity comparable to that of rifampicin, moxifloxacin, and delamanid. In contrast, D-cycloserine and ethambutol showed no activity against non-replicating cells (Figure 2D).

### The putative amidohydrolases AmiC and Rv0552 mediate resistance and susceptibility towards KSK-104 and KSK-106

To elucidate the mechanism of action and resistance, we isolated spontaneous single-step resistant mutants of *M. tuberculosis* H37Rv (Figure S5). All resistant mutants isolated against KSK-104 showed high-level resistance as indicated by 32-fold shift in MIC_90_, and occurred at a frequency of approximately 1 × 10^-8^. For KSK-106, all isolated resistant mutants showed a MIC_90_ shift of approximately 1,024-fold and occurred at a rate < 1 × 10^-10^. Results of whole-genome sequencing of five KSK-104-resistant mutants revealed single nucleotide polymorphisms (SNPs) in the gene *Rv0552,* which encodes a non-essential, conserved hypothetical protein of unknown function (Figure 3A). It is predicted to have amidohydrolase activity and to act on carbon-nitrogen bonds but not on peptide bonds.^13^ The mutations included two independent non-synonymous SNPs resulting in amino acid substitutions, H67R and A229D. The KSK-106-resistant mutants all harbored mutations in the *amiC* gene (Rv2888c), which also codes for a non-essential amidohydrolase (Figure 3A). In contrast to the KSK-104 resistant mutants, we found not only SNPs leading to an amino acid exchange (P185T) but also a SNP resulting in a premature stop codon (E129*) as well as insertions causing a frameshift that most likely led to a non-functional protein (insertion +t at position encoding amino acid 100 of 473, insertion +ccgg at position encoding amino acid 347 of 473). The diversity of identified multiple distinct mutations indicated selection and suggested that resistance to KSKs is associated with loss of function of the identified non-essential amidohydrolase genes. In the majority of the analyzed clones, mutations in *Rv0552* and *amiC* were accompanied by additional mutations. These mutations, however, occurred in diverse genes, which in most cases only had one distinct mutation each. Identical mutations occurring in different mutants strongly suggests that these mutations were copies of a parent clone preexisting in the culture and did not substantially contribute to resistance. In particular, second-site mutations in genes impairing phthiocerol dimycocerosate (PDIM) biosynthesis, such as mutations in *ppsA* and *ppsE*, are known to occur very frequently during *in vitro* culturing of *M. tuberculosis* strains.^14^

**Figure 3.**
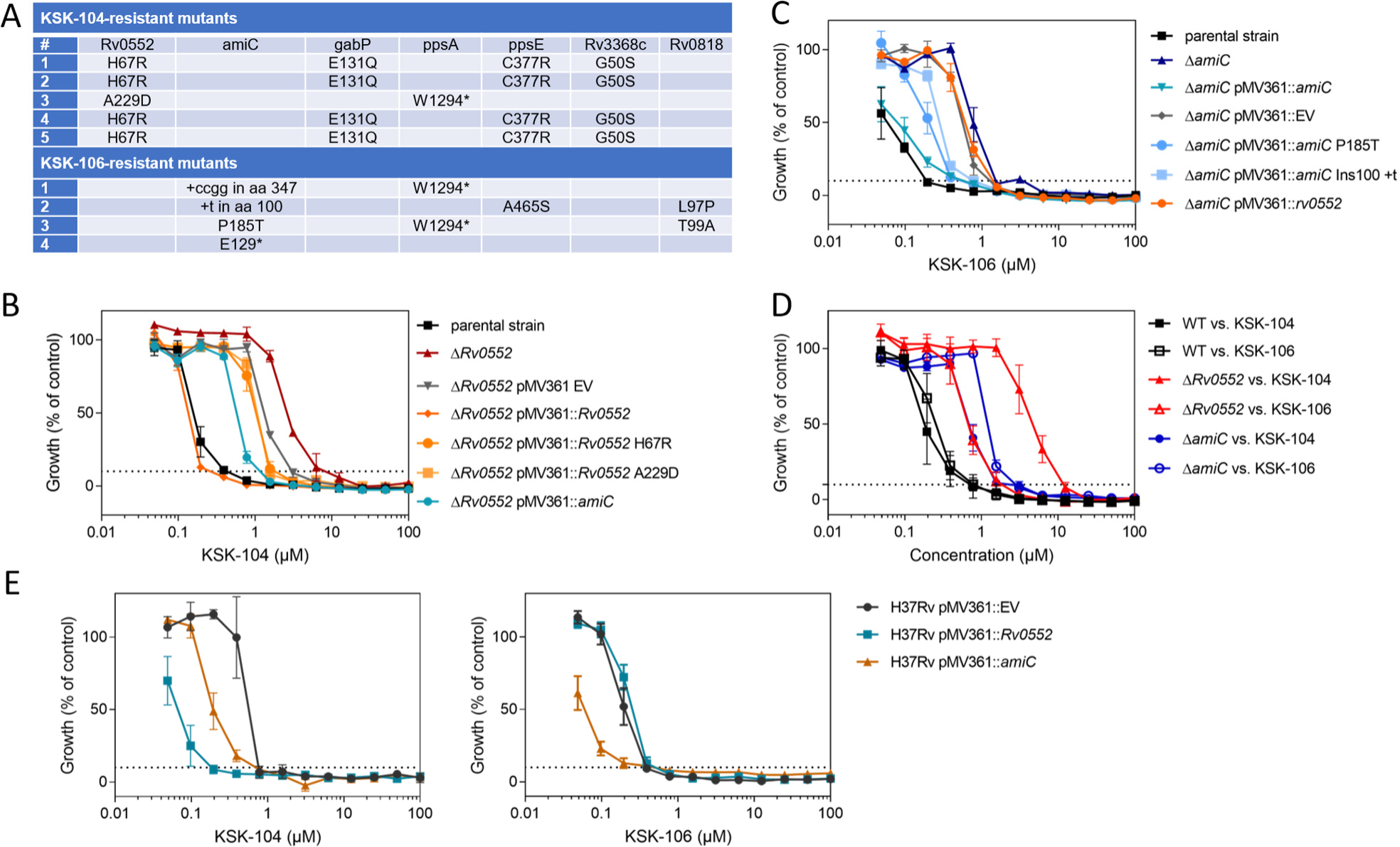
The putative amidohydrolases AmiC and Rv0552 are mediating resistance and sensitivity of *M. tuberculosis* towards KSK-104 and KSK-106. **A)** Overview of mutations identified by whole-genome sequencing in spontaneous resistant *M. tuberculosis* mutants raised against KSK-104 and KSK-106. Dose-response curve of Δ*Rv0552* (**B)** and Δ*amiC* (**C)** knock-out strains against KSK-104 and KSK-106 compared to the parental strain. Deletion of *Rv0552* and *amiC* conferred resistance towards KSK-104 and KSK-106, respectively. Complementation of the gene deletion mutants with integrating plasmids carrying wild-type genes, pMV361::*Rv0552* and pMV361::*amiC*, restored sensitivity in contrast to the ones carrying mutated genes, pMV361::*Rv0552* H67R, pMV361::*Rv0552* A229D, pMV361::*amiC* P185T, pMV361::*amiC* Ins100 +t. Mutants carrying the empty vector control plasmid pMV361::EV served as negative controls. **D)** Cross-resistance of Δ*Rv0552* and Δ*amiC* knock-out strains against KSK-104 and KSK-106. Dose-response curves compared to the parental strain show low-level resistance of the Δ*Rv0552* mutant against KSK-106 and of the Δ*amiC* mutant against KSK-104, respectively. **E)** Overexpression of *amiC* or *Rv0552* leads to increased activity of *M. tuberculosis* H37Rv against KSK-104 and KSK-106. Dose-response curves for KSK-104 (top) and KSK-106 (bottom) showing a concentration-dependent growth inhibition of recombinant strains harboring the empty vector control pMV361::EV or the overexpression constructs pMV361::*amiC* or pMV361::*Rv0552*, respectively. Results in **B**, **C** and **E** are mean ± S.D. from three independent biological replicates. Results in **D** are mean ± S.D. from four to six biological replicates, resulting from two independent experiments each measured at least in duplicate. Growth was quantified by employing the resazurin reduction assay. The dashed lines indicate 10% residual growth.

To confirm the relevance of *Rv0552* and *amiC* in conferring resistance to KSK-104 and KSK-106, we generated individual site-specific *Rv0552* and *amiC* gene deletion mutants in *M. tuberculosis* H37Rv, using specialized transduction (Figure S6). The resulting independent clones of the *M. tuberculosis* Δ*Rv0552* gene deletion mutant were highly resistant (8-fold increase in MIC_90_) against KSK-104 with an MIC_90_ of 12.5 μM (Figure 3B). Genetic complementation of the Δ*Rv0552* deletion mutant with the wild-type *Rv0552* gene constitutively expressed from a single copy integrative plasmid (pMV361::*Rv0552*) fully restored the sensitivity to KSK-104. Site-directed mutagenesis was used to generate plasmids containing the specific mutations found in the spontaneously resistant mutants. Complementation with these mutated versions of the *Rv0552* gene by using plasmids pMV361::*Rv0552* H67R or pMV361::*Rv0552* A229D only marginally restored antitubercular activity (Figure 3B). This confirmed that the resistance phenotype depends only on the loss of function of *Rv0552*, ruling out potential polar effects or relevance of secondary mutations. Likewise, loss of the *amiC* gene in the *M. tuberculosis* Δ*amiC* gene deletion mutant conferred high resistance to KSK-106 with a MIC_90_ of 6.25 μM, corresponding to an 8-fold increase in MIC_90_. The complementation of the Δ*amiC* gene deletion mutant with the wild-type *amiC* gene using plasmid pMV361::*amiC* restored the antitubercular activity of the KSKs, while this effect was not observed when complemented with the mutated versions of the *amiC* gene (Figure 3C). These results demonstrated that the observed resistance to KSK-106 in *M. tuberculosis* is unambiguously linked to loss of function of *amiC*. Interestingly, the Δ*Rv0552* mutant also showed moderate resistance towards KSK-106 with an MIC_90_ of 1.56 μM, while the Δ*amiC* mutant exhibited moderate resistance against KSK-104 with a MIC_90_ of 3.12 μM (each corresponding to a 2-fold increase in MIC_90_), demonstrating some degree of reciprocal cross-resistances (Figure 3D).

To further analyze the role of the putative amidohydrolases, we generated merodiploid strains of *M. tuberculosis* H37Rv, *M. bovis* BCG Pasteur, and *M. smegmatis* using the same integrative plasmids as above (pMV361::*Rv0552* and pMV361::*amiC*) to overexpress *Rv0552* and *amiC*, respectively. In *M. tuberculosis* H37Rv and *M. bovis* BCG Pasteur overexpression led to increased susceptibility towards the KSKs compared to the empty vector control (Figures 3E and S7). In contrast, overexpression of the two putative amidohydrolases in the non-tuberculous species *M. smegmatis* did not result in any activity, indicating that active amidohydrolases are required, but alone not sufficient, for mediating antimycobacterial activity of the KSKs. Of note, while mutations in *amiC* were identified to mediate resistance to KSK-106 in spontaneous resistant mutants, overexpression of *amiC* also increased susceptibility of the recombinant *M. tuberculosis* H37Rv and *M. bovis* BCG Pasteur strains towards KSK-104 (Figure 3E), while overexpression of *Rv0552*, which was identified to mediate resistance to KSK-104 in spontaneous resistant mutants, cross-sensitized recombinant *M. bovis* BCG Pasteur towards KSK-106 (Figure S7).

### KSKs are pro-drugs that are hydrolyzed intracellularly by the amidohydrolases Rv0552 and AmiC

The genetic studies demonstrated that loss of function of the non-essential proteins Rv0552 or AmiC mediates resistance towards the antibacterial KSK compounds, while their overexpression causes hypersensitivity. This phenotype has previously been reported to be associated with the activation of pro-drugs.^15,16^ In fact, AmiC was previously shown to be involved in activation of amide-containing drugs. Indole-4-carboxamides are hydrolyzed by AmiC to yield 4-aminoindole, which acts as an antimetabolite of tryptophan biosynthesis at the stage of tryptophan synthase (TrpAB).^17^ Furthermore, MMV687254, a pyridine carboxamide derivative, also requires AmiC-dependent hydrolysis for antibacterial activity against *M. tuberculosis*, although the antimycobacterial mechanism following hydrolysis remained unknown.^18^ Therefore, we hypothesized that the KSKs are pro-drugs that are hydrolyzed by intracellular amidohydrolases, Rv0552 or AmiC, to bioactive metabolites.

To evaluate the predicted role of AmiC and Rv0552 in the activation of, and resistance to, α-aminooxyacetic acid molecules, further biochemical and structural studies were performed. Structure-homology modeling of Rv0552 using Phyre2^19^ and PyMOL employing an uncharacterized, metal-dependent hydrolase from *Pyrococcus horikoshii* ot3 (PDB ID 3IGH) as the template provided indications that the H67R mutation observed in resistant mutants might interfere with complexation of a Zn^2+^ ion, thereby potentially impairing hydrolytic activity (Figure S8A). Also, the A229D mutation is predicted to be located in close proximity to the Zn^2+^ ion thereby possibly disturbing hydrolytic activity. Structural modeling of AmiC using a fatty acid amide hydrolase from *Arabidopsis thaliana* as the template (PDB ID 6DII) revealed that proline 185 is located within the hydrophobic core. Therefore, the P185T mutation could result in protein destabilization and unfolding (Figure S8B). These observations support that antimycobacterial activity of the KSKs probably relies on hydrolytic activity of Rv0552 and AmiC. To further substantiate this, we performed a multiple sequence alignment of AmiC and found that the protein probably belongs to the amidase superfamily (Figure S8C), which comprises a distinctive catalytic Ser-*cis*-Ser-Lys triad conserved among several known hydrolytic enzyme.^20,21^ We propose that the triad Ser^181^–*cis*-Ser^157^–Lys^82^ represents the catalytic center of AmiC (Figure 4A) as it shares similar positions with that of the 6-aminohexanoate cyclic dimer hydrolase (PDB ID 3A2Q) (Ser^174^–*cis*-Ser^150^–Lys^72^) and the aryl acylamidase (PDB ID 4YJ6) (Ser^187^–*cis*-Ser^163^–Lys^84^).^22,23^ In agreement with this, complementation of the *M. tuberculosis* Δ*amiC* gene deletion mutant with a plasmid expressing a mutated AmiC version, with all three catalytically relevant residues being replaced by alanine (pMV361::*amiC* K82A_S157A_S181A), was unable to restore susceptibility of the recombinant strain to both KSKs (Figure 4B).

**Figure 4.**
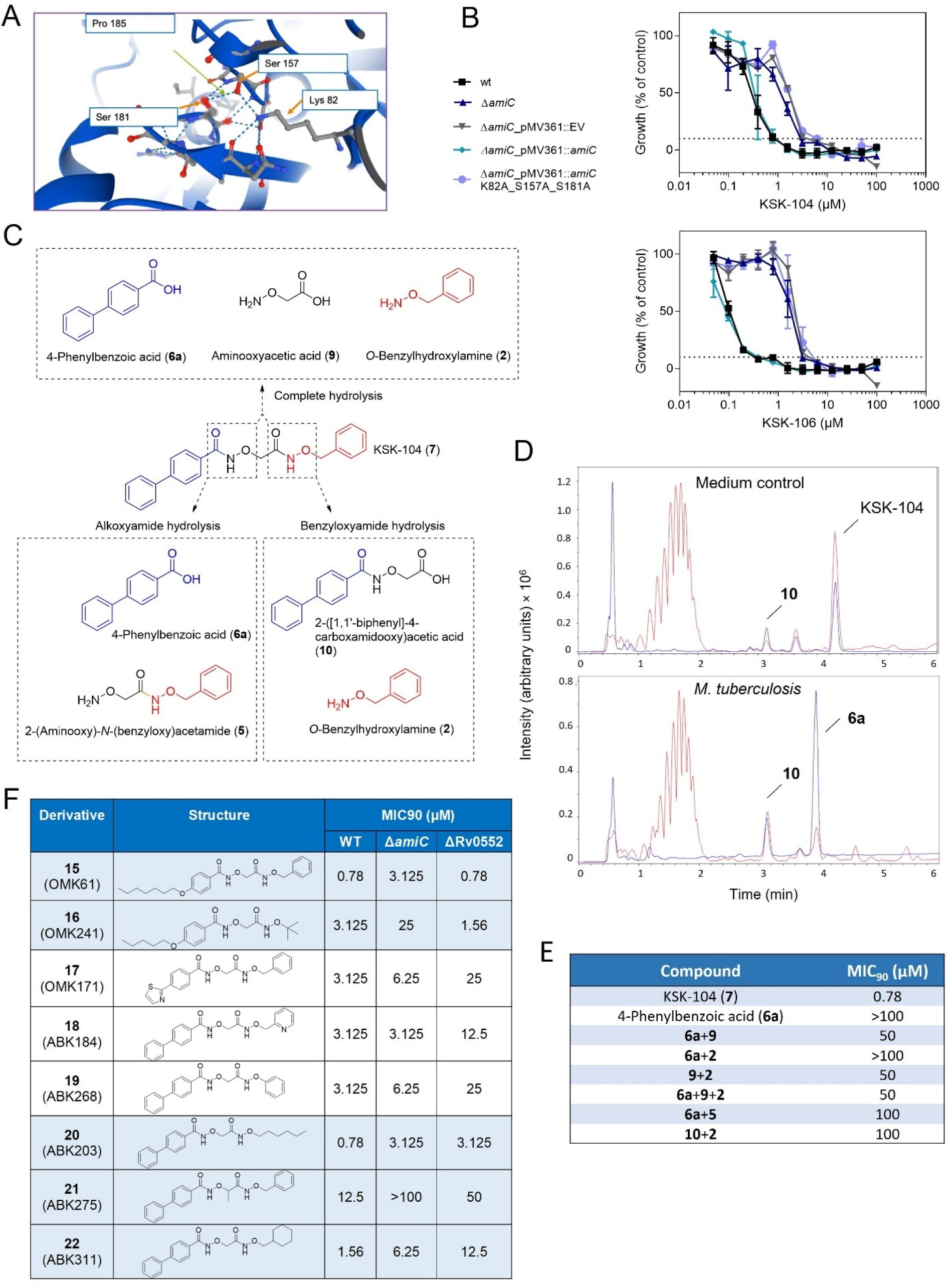
Catalytic activity of Rv0552 and AmiC is required for antimycobacterial effects of KSK-104 and KSK-106. **A)** AlphaFold prediction of the structure of the active center of AmiC, showing the orientation of the catalytic triad Ser^181^–*cis*-Ser^157^–Lys^82^ (https://www.alphafold.ebi.ac.uk/entry/O06418).^25,26^ **B)** Dose-response curve for KSK-104 and KSK-106 of the *M. tuberculosis* Δ*amiC* gene deletion mutant complemented with a plasmid expressing a catalytic site mutant version of AmiC (pMV361::*amiC* K82A_S157A_S181A). The mutant strain containing the empty vector control plasmid pMV361::EV served as the negative control, while the mutant strain expressing wild-type AmiC from plasmid pMV361::*amiC* served as the positive control. Data are means ± SD from three independent biological replicates. Growth was quantified by employing the resazurin reduction assay. The dashed lines indicate 10% residual growth. **C)** Structure of potential hydrolysis products released from KSK-104 by amidohydrolases AmiC and Rv0552. **D)** ESI-LC-MS analysis of methanol extracts obtained after 48 h incubation of 100 µM KSK-104 in sterile 7H9 medium (top) or in 7H9 medium inoculated with *M. tuberculosis* H37Rv cells (bottom). The scan was from 50 to 1500 *m/z* in positive mode. The base peak chromatogram is shown in red, the UV chromatogram at 254 nm is shown in blue. Identified peaks: KSK-104 [*m/z* + H]^+^ = 377.14, **6a** [*m/z* + H]^+^ = 199.07, **10** [*m/z* + H]^+^ = 272.09. **E)** MIC_90_ values of potential KSK-104 hydrolysis products against *M. tuberculosis* H37Rv. Compounds were tested individually and in various combinations. For combination treatments, equimolar mixtures were used containing each compound at the indicated concentration. **F**) Comparison of MIC_90_ values of KSK-104 and KSK-106 derivatives tested against *M. tuberculosis* H37Rv wild type and the Δ*Rv0552* and Δ*amiC* gene deletion mutants.

The hydrolytic activation could affect either the alkoxyamide and/or benzyloxyamide moiety of the lead structures KSK-104 and KSK-106 resulting in different cleavage products (Figure 4C and Figure S9A). To assess whether metabolic cleavage of KSK-104 occurs, the compound was incubated with viable *M. tuberculosis* H37Rv cells. LC-MS analysis of methanol extracts of the cell suspension obtained after 48 h of incubation showed that the parental compound was completely consumed, while two cleavage products accumulated (**6a**, **10**), indicating hydrolysis at both potential cleavage sites (Figure 4D). Incubating KSK-104 in a sterile medium resulted in the formation of small amounts of cleavage product **10**. This suggests that benzyloxyamide hydrolysis of KSK-104 may also occur spontaneously to some extent under the tested conditions, whereas alkoxyamide hydrolysis of KSK-104 is exclusively cell-mediated. KSK-104 hydrolysis to products **6a** and **10** should additionally yield cleavage products **2**, **5** and **9**, which were not detected in the cellular extract. This suggests that further metabolism may occur following the first step of KSK-104 hydrolysis. We confirmed cell-mediated alkoxyamide hydrolysis of KSK-106 by detecting cleavage product **11** (as an analog of **10**) after incubation of *M. tuberculosis* H37Rv cells with KSK-106, while no other predicted cleavage product was detected (Figure S9A+B).

Next, to evaluate whether the antimycobacterial activity of the KSKs arises from a specific hydrolytic product, we tested the susceptibility of *M. tuberculosis* H37Rv cells against all potential KSK-104 (Figure 4E) or KSK-106 (Figure S9C) cleavage products. However, with MIC_90_ values ranging from 25 to >100 µM, the individual compounds were only weakly active or inactive. Furthermore, even different combinations of products that result from alkoxyamide and/or benzyloxyamide hydrolysis were unable to reproduce the effect of the parental compounds (Figure 4E and Figure S9C). One possible explanation is that the tested metabolites, which bear polar carboxyl and aminoxy groups, are polar and potentially charged under the assay conditions and therefore likely exhibit low permeability through the lipophilic mycobacterial cell wall.^24^ From these findings, we conclude that the antibacterial activity of the KSKs relies on efficient diffusion or uptake of the parental compounds by the cells, followed by intracellular hydrolyzis by the amidohydrolases AmiC and/or Rv0552. The antibacterial mechanism(s) might then arise from one or more of the emerging intracellular hydrolysis products which are possibly further metabolized.

To further characterize the contribution of the amidohydrolases AmiC and Rv0552 to susceptibility to KSK compounds, we made use of our flexible synthesis capabilities to synthesize various analogs of KSK-104 and KSK-106 using two different routes (Scheme 2). For the synthesis of the *O-*substituted hydroxamic acids (benzyloxyamides) **15** and **17** via route A, hydroxylamine **5** was acylated with carboxylic acids **6c**-**d** in EDC-mediated amide coupling reactions. For Route B, the hydroxamic acids **12a**-**b** were *O*-alkylated with bromoacetic acid (**1a**) or 2-bromopropanoic acid (**1b**) in alkaline medium to furnish the corresponding carboxylic acids **10**, **11** and **13** as intermediates in good yields. The required regioselectivity of these reactions was investigated by ^1^H-^15^N-HSQC-NMR-spectroscopy. Finally, the carboxylic acids **10**, **11** and **13** were reacted with either *O-*benzylhydroxylamine (**2**) or differently *O-*substituted hydroxylamines **14a**-**e** in EDC-mediated amide coupling reactions to afford the KSK-analogs **16** and **18**-**22**.

The derivatives **15-22** were tested for antibacterial activity against cells of *M. tuberculosis* H37Rv wild type and the Δ*Rv0552* and Δ*amiC* mutants (Figure 4F) (an extensive structure-activity relationship study comprising more than 200 derivatives will be reported elsewhere). We identified that derivatives **15** and **16,** which differ in region C compared to KSK-106, show reduced relative activity only against the Δ*amiC* gene deletion mutant. In contrast, KSK-104 structural variants **17-19** exhibited substantially lower relative activity only with the Δ*Rv0552* mutant, while compounds **20**-**22** differing in region C compared to KSK-104 demonstrated a similar shift in MIC_90_ against both deletion mutants (Figure 4F). As indicated by the reduced antimycobacterial activity, the methyl group in region B of **21** strongly impeded the hydrolysis of the benzyloxyamide group not only in the gene deletion mutants but partly already in the wild type of *M. tuberculosis* H37Rv. In agreement with the increased susceptibility to KSK-104 and KSK-106 observed in the overexpression strains (Figure 3E and Figure S7) and the cross-resistance of the Δ*amiC* and Δ*Rv0552* gene deletion mutant against both KSK-104 and KSK-106 (Figures 3D and 4B), these results collectively suggest that both amidohydrolases can principally be partially redundant pro-drug activators of α-aminooxyacetic acid derivatives, with the preference for AmiC or Rv0552 being influenced by the specific substitution pattern of the compounds.

**Scheme 2.**
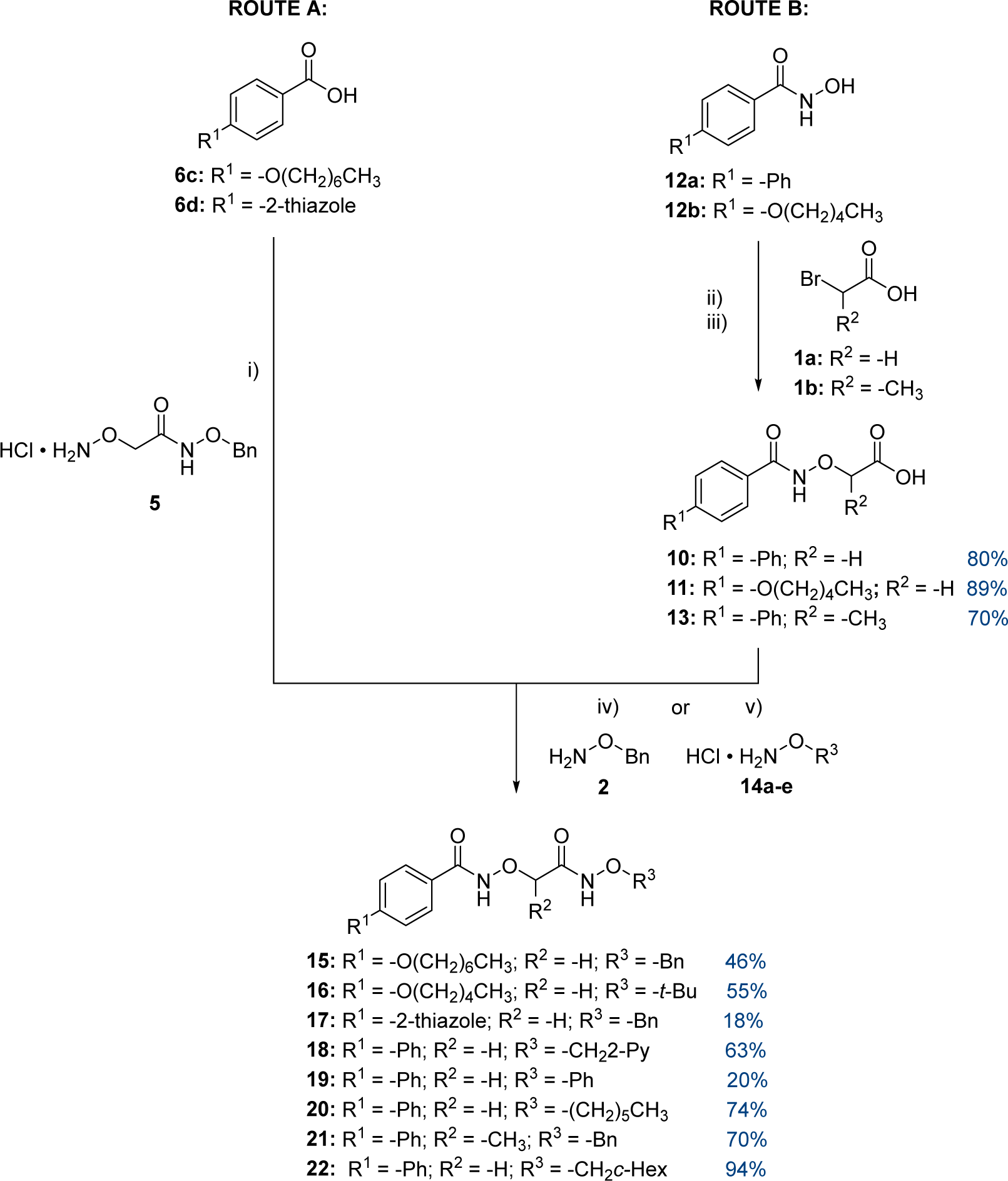
Synthesis of KSK-analogs 15-22. i) 1.25 eq. of hydroxylamine **5**, 0.10 eq. 4-DMAP, 1.25 eq. EDC•HCl, CH_2_Cl_2_, rt; ii) 1.00 eq. bromoacetic acid (**1a**), 2.00 eq. NaOH, EtOH, reflux; iii) 1.10 eq. NaH, THF, −10 °C then 1.00 eq. 2-bromopropanoic acid (**1b**), THF, reflux; iv) 1.25 eq. *O-*benzylhydroxylamine (**2**), 0.10 eq. 4-DMAP, 1.25 eq. EDC•HCl, CH_2_Cl_2_, rt; v) 1.25 eq. H_2_NOR^3^ (**14a**-**e**), 0.10 eq. 4-DMAP, 1.25 eq. EDC•HCl, 1.30 eq. NEt_3,_ CH_2_Cl_2_, rt.

### *α*-Aminooxyacetic acid derivatives might act as “dirty drugs” affecting multiple intracellular targets

To reveal molecular insights into pathways that contribute to the antitubercular effects of the studied compounds, we applied complementary approaches including genetic interaction mapping as well as transcriptomic and proteomic stress response profiling.

For an unbiased identification of pathways or targets that are associated with KSK susceptibility or resistance, we performed a genome-wide quantitative analysis of a saturated transposon mutant pool established in *M. tuberculosis* strain H37Ra, employing transposon-insertion sequencing (TnSeq). The mutant pool was subjected to either DMSO solvent control or to a sublethal concentration of 0.18 μM KSK-106 that was found to decrease growth rate by ca. 50% over five generations (Figure S10). A mean saturation of 67.7% (range 65-69%) was observed between samples, where ‘saturation’ refers to percent of TA dinucleotide sites with one or more insertion. We identified 74 genes with *P_adj_* < 0.05 resulting in apparent fitness changes of transposon mutants using TRANSIT^27^ resampling. Of these genes of interest, 23 met a Log_2_-fold change threshold ≤ −0.55 or ≥ 0.55 and seven a Log_2_-fold change threshold ≤ - 1 or ≥ 1, respectively (Figure 5A). By comparing the composition of the mutant library in the absence and presence of KSK-106, we were able to define 41 genomic regions harboring transposon insertions with a significant increase of mutant abundance during KSK-106 treatment, suggesting that inactivation of the respective genes provides an advantage under the test conditions and contributes to resistance. We also found 33 genomic regions, for which insertions resulted in a significant decrease in mutant abundance over experimental selection, suggesting that these genes are important for fitness under the test conditions as their inactivation led to an increased sensitivity of the cells (Table S1).

**Figure 5.**
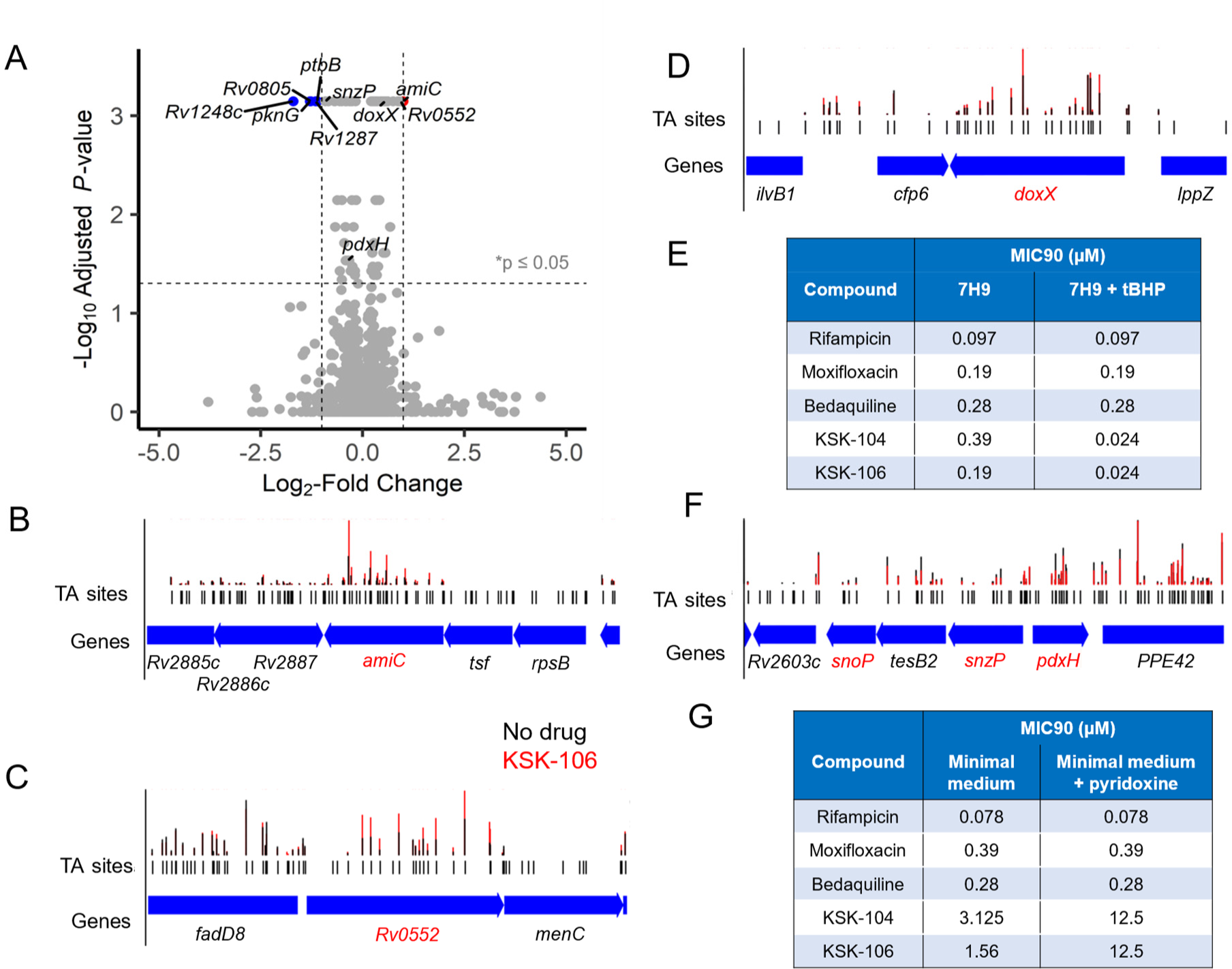
Genes specifically mediating differential susceptibility towards KSK-104 and KSK-106 treatment in *M. tuberculosis* as revealed by Tn-seq analysis. **A)** Volcano plot highlighting transposon mutants selected for or against under selective pressure of subinhibitory concentrations of KSK-106. Transposon insertions in genes with gains in fitness denoted in red. Transposon insertions in genes with loss in fitness denoted in blue. Black dotted line denotes threshold for significance of Log2-Fold change of < −1 or > 1 and adjusted p-value of <0.05. **B - D, F)** Abundance of reads at individual TA dinucleotide insertion sites at selected genomic regions in analyzed mutant pools of *M. tuberculosis* strain H37Ra subjected to sublethal treatment with KSK-106 (red bars) or DMSO solvent control (black bars). Relevant genes are highlighted in red. Enrichment of transposon insertions in the amidohydrolase genes *amiC* (**B)** and *Rv0552* (C) in the KSK-106-treated *M. tuberculosis* H37Ra mutant pool, indicating decreased sensitivity of the corresponding transposon mutants. **D)** Enrichment of transposon insertions in the *doxX* gene in the KSK-106-treated *M. tuberculosis* H37Ra mutant pool, indicating decreased sensitivity of the corresponding transposon mutants. **E)** Amending of 7H9 medium with 0.001 µM *tert*-butyl hydroperoxide (tBHP), a thiol-specific oxidative stressor, resulted in a significantly increased sensitivity of cells of *M. tuberculosis* H37Rv towards KSK104 and KSK-106, as indicated by lowered MIC_90_ values. Cells treated with rifampicin, moxifloxacin and bedaquiline served as negative controls to demonstrate specificity. Growth was quantified by employing the resazurin reduction assay. Measurements were done in triplicates revealing identical MIC_90_ values between samples. **F)** Depletion of transposon insertions in genes involved in pyridoxal 5’-phosphate synthesis (*snzP*) and salvage pathway (*pdxH*) in the KSK-106-treated *M. tuberculosis* H37Ra mutant pool, indicating increased sensitivity of the corresponding transposon mutants. Mutants carrying transposon insertions in the pyridoxal 5’-phosphate synthesis gene *snoP* were also depleted, but did not reach the statistical treshhold. **E**) Supplementation of minimal medium with 100 µg/mL pyridoxine led to resistance of cells of *M. tuberculosis* H37Rv towards KSK104 and KSK-106, as indicated by higher MIC_90_ values. Cells treated with rifampicin, moxifloxacin and bedaquiline served as negative controls to demonstrate specificity. Growth was quantified by employing the resazurin reduction assay. Measurements were done in triplicates revealing identical MIC_90_ values between samples.

Confirming our previous results, mutants harboring inactivating transposon insertions in *amiC* or *Rv0552* led to the strongest enrichment of mutants in the pool upon KSK-106 treatment, consistent with their suggested role in pro-drug activation (Figure 5A-C). These finding are further supported by gain of fitness observed from insertions throughout the open reading frames of these genes (Figure 5B+C). Several genes found to alter mycobacterial fitness in drug-treated cells were already known to affect efficacy against other drugs (Table S2). These are genetic features that likely contribute to general mycobacterial resistance or susceptibility mechanisms that are not specifically linked to the antitubercular effects of KSK-106. These include genes encoding putative drug-efflux pumps (ABC transporter Rv1272c-Rv1273c, the daunorubicin-phthiocerol dimycocerosate-transport ABC transporter DrrA and DrrB, and the resistance-nodulation-cell division (RND) superfamily member MmpL7),^28–31^ where transposon insertions led to higher susceptibility to KSK-106 treatment, suggesting that these transporters mediate efflux of the parental compound or bioactive hydrolysis products. Furthermore, insertions in the universal stress protein family gene *Rv3134c*, reported to be part of a complex involved in the bacterial response to a wide range of stresses, ^32^ resulted in decreased fitness of the corresponding mutants. Additionally, mutants harboring transposon insertions in metabolic genes were also highly overrepresented in the pool following treatment. These included the glycogen metabolism genes *glgC* and *glgP*, and the glycerol kinase gene *glpK*, where mutations have frequently been reported to produce a general drug-tolerant phenotype. ^33–36^ In contrast, transposon insertions in genes implicated in the cell wall structure, such as those encoding PE/PPE family members and enzymes involved in the synthesis of the major cell envelope lipid phtiocerol dimycocerosate (*ppsA*, *ppsB*, *ppsC*, *ppsE*), caused a reduced sensitivity towards KSK-106, which might be linked to a reduced cell permeability of the compound (Table S2).

In addition to these general mechanisms, we found several genes that appeared to specifically alter susceptibility to KSK-106 treatment, providing further insights into the KSK-induced mechanism of action (Table S3). Mutants carrying insertions in the *doxX* gene were enriched in the KSK-106 treated group, conferring a fitness advantage (Figure 5D). DoxX, together with the superoxide detoxifying enzyme SodA and the predicted thiol-oxidoreductase SseA, has been described to form a membrane-associated oxidoreductase complex (MRC) that is responsible for coordinating detoxification of reactive oxygen species and thiol homeostasis during *M. tuberculosis* infection. SseA and DoxX are mediating oxidative recycling of thiyl radicals that are generated by the free radical scavenging activity of mycothiol, while oxidized mycothiol is then recycled by mycothione reductase activity. Superoxide anions might be generated during these enzymatic activities, which are detoxified by the superoxide dismutase SodA. It has been reported that loss of DoxX leads to defective recycling of mycothiol and results in higher sensitivity towards oxidative stressors that react with cytosolic thiols such as *tert*-butyl hydroperoxide (tBHP).^37^ The increased fitness of *doxX* mutants led us to hypothesize that KSK-106 might affect the oxidative stress network of *M. tuberculosis* in a thiol-specific manner. To test this, we investigated synergism between KSK-106 and the thiol-specific oxidative stressor tBHP. Addition of 0.001 µM tBHP resulted in a significantly higher susceptibility of *M. tuberculosis* H37Rv cells with a more than 16-fold decrease in MIC_90_ against both KSK-104 and KSK-106, while sensitivity to the other tested drugs that do not interfere with the cytosolic thiol pool was unot altered (Figure 5E). These results suggest a mechanism by which the treatment with KSK compounds might lead to the production of free radicals that are detoxified in a mycothiol-dependent manner. Inactivation of DoxX by transposon insertions decreases sensitivity towards the compounds by preventing the generation of toxic superoxide anions under these conditions, while the overload of this detoxification system by exogenous tBHP could potentially promote superoxide anion formation.

We also identified a distinct set of genes that are involved in the pyridoxal-5’-phosphate (PLP) synthesis and salvage pathway. *De novo* biosynthesis of PLP in *M. tuberculosis* is mediated by PLP synthase, a complex consisting of the PLP biosynthesis protein SnzP, encoded by the gene *Rv2606c*, and the putative glutamine amidotransferase SnoP,^38^ while the pyridoxamine 5’-phosphate oxidase PdxH, encoded by the neighboring gene *Rv2607c*, is involved in PLP salvage.^39^ Transposon insertions in *snzP* and *pdxH* caused a loss of fitness and rendered the cells more sensitive toward KSK-106 treatment (Figure 5F), suggesting that KSK-106 treatment might trigger a higher PLP demand within the cell. To test this, we supplemented a PLP-free minimal medium with 100 µg/mL pyridoxine and observed a 4-8-fold increase in MIC_90_ for KSK-104 and KSK-106 treated cells compared to non-supplemented medium, whereas no change was observed for the control antibiotics rifampicin, moxifloxacin, and bedaquiline (Figure 5G).

Next, since interrogation of fitness changes by Tn-Seq is largely limited to non-essential genes, we conducted differential transcriptional and proteomic profiling of KSK-106-treated and untreated cells as alternative approaches to asses which genes or proteins are linked to the mode of action of the α-aminooxyacetic acid derivatives. For transcriptomic analysis of *M. tuberculosis* mc^2^6030, exponential phase cells were exposed to an inhibitory dose of 1.9 µM KSK-106 (5 × MIC_90_) for 24 h and compared to DMSO-treated controls. Under this condition, KSK-106 treatment caused a well-defined and very narrow effect on the transcriptome of *M. tuberculosis*. Four genes belonging to the gene cluster *Rv3092c*-*Rv3095* were highly upregulated in treated cells (Figure 6A). In addition, *Rv3096*, which is adjacent to this gene cluster, was also upregulated more than 2-fold (p< 0.05). Although little is known about this gene cluster, two recent studies have explored some of its functions.^40,41^ The gene *Rv3095* (*mxyR*) encodes the mycobacterial xylan regulator (MxyR), which is a member of the family of multiple antibiotic resistance (MarR) transcriptional regulators and may play a role in the metabolic regulation of carbohydrates, including xylan, L-arabinose and galactose.^41^ The HTH-type transcriptional regulator gene *Rv3095* is divergently oriented to genes encoding a hydrolase (*Rv3094c*), an oxidoreductase (*Rv3093c*), and an ABC transporter (*Rv3092c*) and convergently oriented to the putative xylanase gene *Rv3096*. It was shown that Rv3094c is likely a flavin-dependent monooxygenase with an FAD-binding site and acyl-CoA dehydrogenase activity that is involved in ethionamide activation by sulfoxidation,^41^ while the main route for multistage ethionamide pro-drug activation occurs through conversion to active radicals by the Baeyer-Villiger monooxygenase EthA, followed by further conversion to a toxic adduct with NADH.^42^

**Figure 6.**
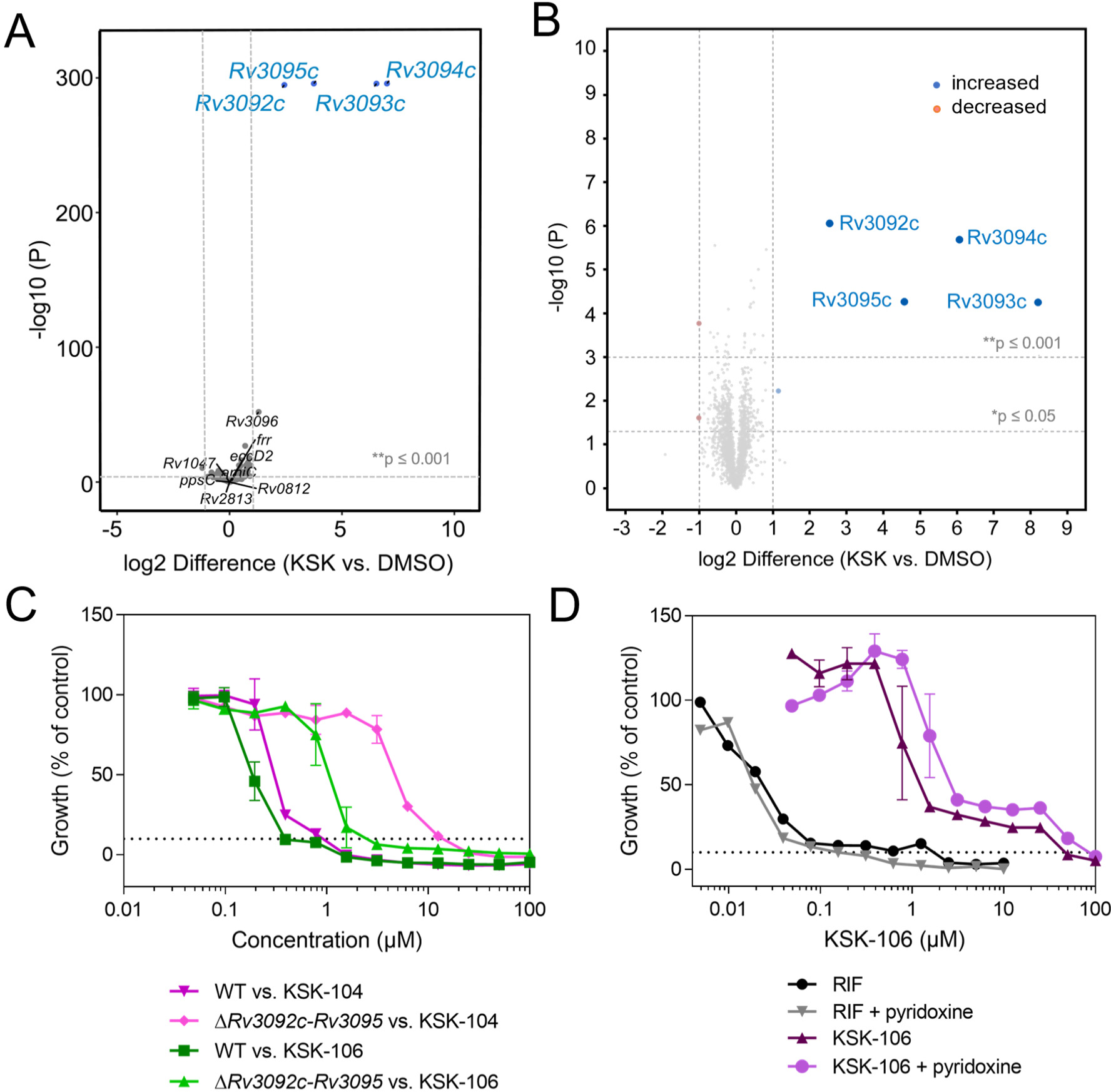
Upregulation of the gene cluster Rv3092c-Rv3095 as a misled stress response in KSK-106 treated *M. tuberculosis* cells. **A)** Full transcriptome analysis of cells of *M. tuberculosis* strain mc^2^6230 treated with a lethal concentration of KSK-106 compared to DMSO control. Exponential phase cells were exposed to 1.9 µM KSK-106 (5 × MIC_90_) for one generation time (24 h) and compared to DMSO treated controls. The plot shows the fold-change (log_2_) in gene expression abundance, plotted against p-value (-log_10_). **B)** LC-MS/MS-based whole protein analysis of silenced cells of *M. tuberculosis* H37Rv treated with a sublethal concentration of KSK-106 (0.05 µM, corresponding to 0.125 × MIC_90_) compared to DMSO control. The volcano plot illustrates the log_2_-fold change in abundance in KSK-106 treated vs. non-treated cells (X-axis) and corresponding −log_10_ p values (Y-axis). Proteins complying with the chosen threshold of significance and showing a log_2_-fold change ≥ 1 or ≤ −1 are marked in blue or red, respectively. Quantification was done via label free quantification (LFQ) of four to five replicates per sample group. To identify statistically significant hits from the analysis, P ≤ 0.05 (Student’s T-test; permutation-based FDR with 250 randomizations and FDR = 0.01) was applied. **C)** Dose-response curves of *M. tuberculosis* H37Rv wild type and the Δ*Rv3092c*-*Rv3095* gene deletion mutant against KSK-104 and KSK-106, demonstrating that the gene deletion leads to resistance against the compounds. **D)** Dose-response curves of the *M. tuberculosis* H37Rv Δ*Rv3092c*-*Rv3095* gene deletion mutant against KSK-106 during cultivation in minimal medium with or without 100 µg/mL pyridoxine, leading to increased resistance in presence of pyridoxine. Cells treated with rifampicin (RIF) served as negative control to demonstrate specificity. Data in **C**+**D** are shown as means of triplicates ± SD. Growth was quantified employing the resazurin reduction assay. The dashed lines indicate 10% residual growth.

To corroborate the transcriptomic results, we investigated the protein stress response of *M. tuberculosis* H37Rv cells following KSK-106 treatment. For this, *M. tuberculosis* H37Rv cells were treated with a sublethal concentration of KSK-106 (0.05 µM, corresponding to 0.125 × MIC_90_) for 10 days. Proteomic analysis revealed again a very distinct and narrow response profile with a high abundance of the four proteins Rv3092c-Rv3095 in treated cells compared to the DMSO control (Figure 6B), confirming the results of the transcriptome analysis. A similar response was observed for cells of *M. tuberculosis* H37Rv subjected to 0.2 µM KSK-106 treatment (corresponding to 0.5 × MIC_90_) (Figure S11). Such a distinctive and narrow transcriptomic and proteomic response profile is very unusual and has yet not been reported to occur in response to treatment with other antitubercular antibiotics. To further elucidate the role of the respective gene cluster in resistance and susceptibility towards KSK compounds, we generated a site-specific *M. tuberculosis* Δ*Rv3092c*-*Rv3095* gene deletion mutant and tested its susceptibility against KSK-104 and KSK-106. Surprisingly, cells lacking the described genes demonstrated marked resistance against KSK-106 with an 8-fold increase in MIC_90_, and high-level resistance to KSK-104 with a 16-fold increase in MIC_90_ values (Figure 6C). This implies that loss of this gene cluster provides a fitness advantage during treatment with α-aminooxyacetic acid derivatives and suggests that the strong and specific upregulation might represent a misguided stress response that enhances the antitubercular effect of the compounds. We finally assessed whether identified pathways that influence susceptibility to KSK-104 and KSK-106 cooperate in the antitubercular mechanism of α-aminooxyacetic acid derivatives. For this, we tested the susceptibility of the *M. tuberculosis* Δ*Rv3092c*-*Rv3095* gene deletion mutant in the presence of pyridoxine and observed enhanced resistance, demonstrating additive effects (Figure 6D).

## DISCUSSION

In this study, we have elucidated the antitubercular properties of KSK-104 and KSK-106, which belong to the family of α-aminooxyacetic acid derivatives that might pave the way for development of a new class of chemotherapeutics against *M. tuberculosis* infections. These compounds show antibacterial activity against *M. tuberculosis* in vitro and in infected human macrophages in a sub-micromolar range and are active against extensively drug-resistant (XDR-TB) clinical isolates. Furthermore, the KSKs show a large therapeutic window with excellent activity and no cytotoxicity in a broad panel of human cell lines. With a specific effect on tuberculous mycobacteria, the KSKs could facilitate the development of narrow-spectrum bactericidals that offer a reduced risk of resistance development and side effects, while also providing a greater opportunity to maintain a normal commensal microbiota in treated patients.^43,44^ Finally, the novel compounds constitute a group of chemical entities that are readily available with a straightforward synthesis route, allowing further derivatization for medicinal chemical optimization.

Similar to other anti-TB medications like isoniazid or ethionamide, the KSKs are pro-drugs. Activation is mediated through hydrolysis by the amidohydrolases AmiC and Rv0552. Genetic studies involving gene deletion mutants and overexpressing recombinant strains revealed that both amidases can simultaneously catalyze pro-drug activation, albeit with various efficacies depending on the substitution pattern of the studied molecules. This partially redundant activation mechanism might partly explain the observed low resistance frequencies. Development of α-aminooxyacetic acid derivatives that represent efficient substrates for both AmiC and Rv0552 might further decrease the likelihood of resistance emergence. AmiC has recently been discovered to mediate the activation of two other classes of amide-containing pro-drug compounds, indole-4-carboxamides and pyridine carboxamides.^17,18^ However, for these compounds, monoactivation only by AmiC occurs, while the studied α-aminooxyacetic acid derivatives differ by involving alternative activation by AmiC and Rv0552. To our knowledge, Rv0552 has not previously been reported to be involved in activation of, or resistance to, antimycobacterial compounds. We acknowledge that redundant amidohydrolase activation of α-aminooxyacetic acid derivatives in the bacilli is attractive, as it will decrease resistance frequency, but substrate promiscuity might also impair *in vivo* efficacy when the compounds are hydrolyzed by host amidohydrolases before they reach the tubercle bacilli. High plasma stability as well intracellular activity of the studied compounds in infected macrophages indicate that the current candidates are not substantially targeted by host amidohydrolases under the tested conditions, but further in-depth pharmacokinetic studies are required to explore metabolic stability in humans.

Pro-drug activation of α-aminooxyacetic acid derivatives by AmiC and Rv0552 can lead to the formation of various hydrolysis products. Identification of predicted hydrolysis products by LC/MS indicates both alkoxyamide and benzyloxyamide hydrolysis. Since even treatment with the combination of different possible hydrolysis products did not mimic the antibacterial effect of the parental compounds, we conclude that AmiC- and Rv0552-mediated hydrolysis needs to occur intracellularly following the uptake of the parental molecules. At least some of the hydrolysis products are likely subject to further metabolism, as we were unable to detect every corresponding hydrolysis product by LC/MS. Which of the resulting hydrolysis products is responsible for the antitubercular activity of the parental compounds remains to be elucidated, but the observed pleiotropic effects suggest that it is the combination of two or more hydrolysis products or their resulting metabolites that provokes bacterial cell death.

Thus, we propose that the studied α-aminooxyacetic acid derivatives likely act as “dirty drugs” simultaneously attacking different intracellular targets or pathways following intracellular hydrolysis. “Dirty drugs” were previously defined as small (molecular weight ranging from 100-300 g/mol), “fragment-like” antimycobacterials that can hit multiple targets and pathways inside the tubercle bacillus, and were shown to be less prone to the rapid emergence of resistance.^45^ Combining complementary untargeted genome-wide genetic, transcriptomic and proteomic analyses, we identified three distinct mechanisms that may be involved in determining susceptibility of *M. tuberculosis* to the studied α-aminooxyacetic acid derivatives.

Genetic interaction mapping connected the activity of the compounds to the oxidative stress network in *M. tuberculosis* cells by demonstrating the specific sensitizing effects of the oxidative stressor tBHP and the membrane protein DoxX. DoxX is an integral membrane protein that facilitates the coordination between cytosolic SseA and secreted SodA.^37^ In the cytosol, the free radical scavenging activity of mycothiol generates mycothiol thiyl radicals. DoxX and SseA enable the conversion of these thiyl radicals into oxidized mycothiol, which is subsequently recycled through the activity of mycothione reductase. As a result of this process, superoxide anions might be produced, that are detoxified by the associated superoxide dismutase SodA.^37^ When DoxX is inactivated, fewer superoxide anions may be generated, which might have less severe effects than the accumulation of free radicals. tBHP is known to interact with cytosolic mycothiol,^46,47^ which promotes generation of superoxide anions. Our findings are in agreement with mechanisms where the KSK compounds are either inhibiting the activity of SodA to detoxify the generated superoxide anions, directly or indirectly promote the generation of superoxide anions (e.g., by interfering with the respiratory electron transport chain), or lead to the production of free radicals that need to be detoxified in a mycothiol-dependent manner. The exact underlying mechanism needs further investigation. The elevated levels of oxidative stress in human phagocytic cells during infection^48^ could enhance the activity of the KSK compounds *in vivo*.

In addition, we found that the PLP synthesis and salvage pathways influence sensitivity towards α-aminooxyacetic acid derivatives. PLP-dependent enzymes are a target for drug therapy in TB, as evidenced by the second-line drug D-cycloserine, underscoring the potential of PLP biosynthesis as a promising drug target.^49,50^ The fitness of the cells under KSK-106 treatment is impacted by mutations in the PLP synthesis gene *snzP* and the PLP salvage gene *pdxH*. PdxH enzymatically oxidizes PNP to produce PLP with the aid of flavin mononucleotide (FMN) as a prosthetic group, whereby the C4’ alcohol PNP is oxidized to an aldehyde. During catalysis, FMN serves as the immediate electron acceptor, with molecular oxygen acting as the final electron acceptor and generating hydrogen peroxide under aerobic conditions.^51^ It has been reported that PLP can function as a potent quencher of reactive oxygen intermediates.^52,53^ Our study has demonstrated that exogenous supplementation of pyridoxine in a minimal medium confers resistance of *M. tuberculosis* cells towards KSK-104 and KSK-106. These results suggest that the studied α-aminooxyacetic acid derivatives may inhibit PLP-biosynthesis, otherwise generate an increased demand for PLP, or promote the formation of reactive oxygen species that cannot be quenched in the absence of sufficient PLP supply. Furthermore, complete alkoxyamide and benzyloxyamide hydrolysis of the studied KSKs will generate aminooxyacetic acid (compound **9**), which is a well-known inhibitor of PLP-dependent enzymes.^54^ It attacks the Schiff base linkage between PLP and the enzyme, generating PLP oxime *O*-acetic acid.^55^ While we found that exogenous aminooxyacetic acid has no direct antibacterial effects on whole cells likely due to poor uptake, the intracellular hydrolytic release of this metabolite might result in inhibition of PLP-dependent enzymes. While bioinformatic prediction of PLP-dependent enzymes is difficult due to low sequence similarities, *M. tuberculosis* is known to harbor at least 30 different PLP-dependent enzymes,^56^ several of which are known to be essential for viability. Pleiotropic attack of these PLP-dependent enzymes by aminooxyacetic acid could contribute to the potent antitubercular effect of the KSKs. Recently, it was shown that *M. tuberculosis* produces hydrogen sulfide (H_2_S) in an PLP-dependent manner and that this process can be inhibited by aminooxyacetic acid.^57^ In *Escherichia coli*, endogenously produced H_2_S protects the cells against oxidative stress.^58^ Thus, blocking PLP-dependent H_2_S formation by intracellular relase of aminooxyacetic acid might contribute to the increased sensitivity of *M. tuberculosis* cells towards oxidative stress during KSK treatment.

Furthermore, transcriptomic and proteomic analyses of KSK-106 treated cells revealed the role of the gene cluster *Rv3092c*-*Rv3095* in determining sensitivity towards α-aminooxyacetic acid derivatives. Inactivation of the genes led to notable cross-resistance towards the studied compounds. Little is known about the biological function and specificity of the corresponding proteins. *Rv3092c*-*Rv3094c* is potentially organized in an operon, whose expression is controlled by Rv3095, which is a transcriptional regulator belonging to the MarR family.^59^ While Rv3093c putatively represents a flavin adenine dinucleotide/flavin mononucleotide (FAD/FMN) reductase, Rv3094c was recently shown to be a putative FMN-containing monooxygenase that catalyzes the sulfoxygenation of ethionamide to its S-oxide (ETH-SO) as the first activation step,^40^ which then forms a covalent adduct with NAD (ETH-NAD) that targets InhA.^60,61^ It is unlikely that the α-aminooxyacetic acid derivatives are directly bioactivated by Rv3094c in a similar manner as they do not contain sulfur. However, it is conceivable that the monooxygenase activity of Rv3094c might generate reactive oxygen species that exacerbate the effect of the KSK compounds. In this regard, we conclude that the strong upregulation of the gene cluster represents a misguided stress response. It was reported that Rv3095 negatively regulates the expression of *Rv3093c* and *Rv3094c*.^40^ Counterintuitively, KSKs trigger a stress response that results in the complete deregulation of the gene cluster, where Rv3095 fails to repress the expression of *Rv3093c* and *Rv3094c*. It remains unknown how the KSK compounds elicit this highly specific upregulation.

In conclusion, we here present the synthesis and structural modification of α-aminooxyacetic acid derivatives as a class of bactericidal antitubercular pro-drugs with promising cytotoxicity profile that are active against drug-resistant clinical isolates and exhibit potent intracellular activity in a human macrophage infection model. Partially redundant bioactivation is mediated by the amidohydrolases AmiC and Rv0552 that intracellularly hydrolytically unleash the “dirty drugs” and trigger pleiotropic effects. Future research will aim to elucidate the precise intracellular antibacterial mechanisms and to identify the relevant metabolites. Further structural optimization of the KSK compounds, alongside the performance of *in-vivo* studies, may pave the way for the development of urgently needed *M. tuberculosis*-specific anti-TB drugs.

## MATERIAL & METHODS

### Generation and cultivation of bacteria

Strains, oligonucleotides and plasmids used in this study are listed in Tables S4-S6. Mycobacterial cultures were grown aerobically at 37 °C shaking at 80 rpm in Middlebrook 7H9 liquid media supplemented with 10% ADS (0.8% NaCl, 5% BSA, 2% dextrose), 0.5% glycerol, 0,025% tyloxapol, and appropriate antibiotics (50 µg/mL hygromycin, 20 µg/mL kanamycin). For growth of *M. tuberculosis* on solidified medium, Middlebrook 7H10 agar supplemented with 10% ADS (5% (w/v) bovine serum albumin; 2% (w/v) glucose; 0.085% (w/v) sodium chloride) and 0.5% (v/v) glycerol was used. For testing growth on media without pyridoxine, *M. tuberculosis* strains were grown in liquid minimal medium [per liter: 0.15 g l-Asparagine × H_2_O, 0.5 g (NaH_4_)_2_SO_4_, 1 g KH_2_PO_4_, 2.5 g Na_2_HPO_4_, 50 mg ferric ammonium citrate, 0.5 g MgSO_4_ × 7 H_2_O, 0.5 mg CaCl_2_, 0.1 mg ZnSO_4_, 0.05% (v/v) tyloxapol, pH 7.0 + 10% ADS and 0.5% glycerol]. *E. coli* cells were grown in lysogeny broth (LB)-medium or LB agar containing the respective antibiotics (150 µg/mL hygromycin, 40 µg/mL kanamycin, 100 µg/mL ampicillin).

### Minimum inhibitory concentration assay

The minimum inhibitory concentrations of the tested compounds were quantified by dose-response curves using the resazurin microplate assay. In short, a two-fold serial dilution of tested compounds was prepared in a polystyrene U-bottom 96-well plate (Greiner) to result in dose-response curves ranging from 100 μM to 0.048 μM final concentrations. 50 µl of exponentially growing cells (OD_600 nm_ ≤ 1, diluted to 1 x 10^6^ CFU/mL) were then added into each well to yield a total volume of 100 μl and cultivated for five days at 37°C (5% CO_2_, 80% humidity). Subsequently, 10 μl resazurin solution (100 μg/ml, Sigma Aldrich) was added into each well and incubated overnight. Cells were fixed for 30 min at room temperature after the addition of 10% (v/v) formalin. Growth was quantified based on fluorescence using a microplate reader (TECAN) (excitation: 540 nm, emission: 590 nm). Relative growth was calculated to the DMSO solvent control (= 100% growth) and uninoculated wells (subtraction of background fluorescence = 0% growth). Experiments were performed in triplicates. MIC_90_ values are given as discrete, stepwise values representing the actually tested lowest compound concentration that resulted in 10% residual growth or less.

### Cytotoxicity assay

To determine the cytotoxicity of the compounds *in vitro*, human cell lines derived from different tissues were used. THP-1 cells (leukemia monocytic cell line), CLS-54 (adenocarcinoma-derived lung epithelial cell line), and HUH7 (hepatocyte-derived carcinoma cell line) were grown in RPMI supplemented with 10% fetal bovine serum (FBS). H4 (neuroglioma cell line) and SH-SY5Y (neuroblastoma cell line) cell lines were cultivated in DMEM supplemented with 10% FBS. MRC-5 (normal lung fibroblasts), HEK293 (epithelial-like embryonic kidney cell line), and HEPG2 (hepatocellular carcinoma cell line) cells were grown in EMEM supplemented with 1% non-essential amino acids, 1 mM sodium pyruvate, and 10% FBS. Sterile 96-well flat-bottom polystyrene plates (Greiner) were prepared with a two-fold serial dilution of compounds with 100 μM as the highest concentration. Approx. 5×10^4^ cells were seeded in each well in a total volume of 100 μL per well. The cells were incubated for 48 h at 37°C and 5% CO_2_ before viability was quantified using the resazurin reduction assay as described above. All cell lines were obtained from CLS Cell Lines Dienstleistung GmbH.

### Killing curve assay

*M. tuberculosis* H37Rv cells were growing in Middlebrook 7H9 supplemented with 10% ADS, 0.5% glycerol, and 0.05% tyloxapol to the exponential phase. This pre-culture was used to prepare cultures containing 10^6^ CFU/mL, which were incubated either with KSK-104 or KSK-106 (0.25 μM) individually or in combination with the anti-tubular drugs isoniazid (10 μM), rifampicin (1 μM), bedaquiline, (0.5 μM), delamanid (0.5 μM) or ethambutol (10 μM). The cultures were incubated shaking at 80 rpm at 37 °C for 35 days. At indicated time points, aliquots were taken, serially diluted, and plated on Middlebrook 7H10 agar plates supplemented with 10% ADS and 0.5% glycerol to quantify colony forming units to determine the effects on the growth of *M. tuberculosis*. After 3 weeks colonies were counted. All experiments were performed as triplicates.

### Macrophage Infection Assay

THP-1 cells were grown in RPMI medium supplemented with 10% FBS. Cells were counted using a haemocytometer and 10^5^ cells were seeded into each well of a sterile 96-well flat-bottom polystyrene microtiter plate (Greiner) in a total volume of 100 μL. To differentiate the cells to adherent macrophage-like cells, the medium was supplemented with 50 nM phorbol-12-myristate-13-acetate.^62^ After differentiation to adherent cells overnight, the macrophage-like cells were washed twice with PBS. An mCherry expressing recombinant *M. tuberculosis* H37Rv reporter strain was used for infection. Cells were grown in Middlebrook 7H9 broth containing 150 μg/mL hygromycin, harvested, washed and resuspended in RPMI supplemented with 10% FBS to a density of 3×10^6^ CFU/mL. 100 μL of this cell suspension was added to each well, resulting in a multiplicity of infection = 3. After 3 h, cells were washed twice with PBS to remove non-phagocytosed bacteria. PBS was replaced with 100 μL RPMI + 10% FBS containing 0.5 μM KSK-104 or KSK-106 or different antibiotics (3 μM rifampicin, 20 μM streptomycin). After 5 days at 37°C, 5% CO_2_ and 85% humidity, the macrophage-like cells were fixed with formalin (5% final concentration), and fluorescence was detected using a Nikon Eclipse TS100 fluorescence microscope. Additionally, viable cell counts were determined by lysing macrophages with ddH_2_O for 30 minutes. Dilutions of each well were plated on Middlebrook 7H10 plates and colonies were counted after 3 weeks of incubation at 37°C.

### Starvation-induced non-replicating persistence model

To test activity of compounds against non-replicating cells of *M. tuberculosis* H37Rv, cells were grown to stationary phase, harvested, washed thrice with PBS+0.025% tyloxapol, resuspended in PBS + 0.025% tyloxapol in the original culture volume and starved by incubation at 37 C for three weeks. Next, cells were diluted to 1×10^8^ CFU/mL with PBS + 0.025% tyloxapol and transferred into 96-well round bottom microtiter plates to a final volume of 100 µl per well, and compounds were added at the indicated final concentrations. After five days of incubation at 37 °C as standing cultures, resazurin solution (10 μl/well from 100 μg/mL stock) was added, and cells were incubated for 48 h at 37 C.

### Isolation of spontaneous resistant mutants and whole genome sequencing

Spontaneous resistant mutants were isolated by plating each approximately 6×10^7^ – 1x 10^8^ cells of *M. tuberculosis* H37Rv or the respective merodiploid or gene deletion strain, on solid media containing either 4, 6, or 10-fold MIC concentrations of the KSK derivatives, respectively. After four to six weeks, colonies were isolated. Genomic DNA from *M. tuberculosis* was isolated using the cetyltrimethylammonium bromide (CTAB)-lysozyme method as described by Larsen et *al.*, ^63^ quantified using the AccuClear® Ultra High Sensitivity dsDNA Quantitation Kit, and quality was measured by capillary electrophoresis using the Fragment Analyzer and the ‘High Sensitivity genomic DNA Assay’ (Agilent Technologies, Inc.). Genomes of resistant mutants were sequenced with an Illumina HiSeq 2500 next generation sequencer (at Texas A&M University, College Station, TX, USA) after preparing sequencing libraries using standard paired-end genomic DNA sample prep kit from Illumina. Paired-end sequence data was collected with a read length of 150 bp. Base-calling was performed with Casava software, v1.8. The reads were assembled using a comparative genome assembly method, with *M. tuberculosis* H37RvMA as a reference sequence (GenBank accession GCA_000751615.1).^64^ The mean depth of coverage ranged from 214-282x for KSK-104 mutants, and 12-28x for KSK-106 mutants.

### Heterologous expression in mycobacteria

The respective gene regions were amplified using the designed oligonucleotides (Table S5) and cloned into the expression vector pMV361, which contains a kanamycin resistance gene and a strong constitutive *hsp60* promoter, using *Hin*dIII and *Pac*I (New England Bioloabs) restriction sites by chemically transformation of *E. coli* NEB-5α cells. Sequenced plasmids containing the respective gene of interest or the pMV361::empty vector (EV) were electroporated into *M. smegmatis* mc^2^155, *M. bovis* BCG Pasteur or *M. tuberculosis* H37Rv and plated on 7H10 selective plates as described previously.^63^ Single colonies were picked and grown in selective 7H9 media after three weeks of incubation.

### Construction of targeted gene deletion mutants

Specialized transduction was employed to achieve gene disruptions in *M. tuberculosis* H37Rv.^65^ Briefly, an allelic exchange substrate was designed to replace the gene/s of interest in *M. tuberculosis* with a γδ*res*-*sacB*-*hyg*-γδ*res* cassette comprising a *sacB* as well as a hygromycin resistance gene flanked by *res*-sites of the γδ-resolvase. Upstream and downstream flanking regions of the gene/s of interest were amplified by PCR using primers listed in Table S5. Subsequently, the flanking regions were digested with the indicated restriction enzymes and ligated with the *Van91*I-digested p0004S vector. The resulting allelic exchange plasmid was then linearized with *PacI*, cloned, and packaged in the temperature-sensitive phage ФphAE159, yielding knock-out phages that were propagated in *M. smegmatis* at 30 °C. Allelic exchange in *M. tuberculosis* was performed through specialized transduction at the non-permissive temperature of 37°C, using hygromycin for selection. This led to the resulting gene deletion/s and replacement by the γδ*res*-*sacB*-*hyg*-γδ*res* cassette. After isolation of genomic DNA, the obtained hygromycin-resistant transductants were then screened for correct gene disruption by diagnostic PCR analysis (Figure S6).

### Site-directed mutagenesis

Suitable primer pairs (Table S5) were designed utilizing the Q5 site-directed mutagenesis kit (New England Biolabs) to generate mutated amidases. The pMV361::*Rv0552* or pMV361::*amiC* constructs were employed as templates and the procedure was conducted as instructed by the manufacturer. The resultant products were then transformed into competent *E. coli* NEB-5α cells. Plasmids containing the predicted mutations were confirmed through Sanger-sequencing.

### Protein homology modelling

The Phyre2 web portal was used for protein modelling, prediction and analysis.^19^ Regarding Rv0552, 474 residues (equivalent to 89% of the sequence) were modelled with a 100.0% accuracy rate by the single highest-scoring template. The template for Rv0552 was presented by an uncharacterized, metal-dependent hydrolase from *Pyrococcus horikoshii* ot3 (PDB ID: 3igh). AmiC has also been modelled with a 100.0% confidence rate using the highest-scoring template, resulting in 466 residues being modelled effectively. The structure of *Arabidopsis* fatty acid amide hydrolase was utilized (PDB ID: 6DII) as a template. For further structural analysis, the software tool PyMol was used.^66^

### Transposon mutagenesis and sequencing

*M. tuberculosis* H37Ra cells were mutagenized with the mariner *himar1* transposon via the temperature-sensitive mycobacteriophage phAE180.^67,68^ Cultures containing the mutagenized cells, with a starting inoculum of OD_600 nm_ = 0.01, were grown on Middlebrook 7H9 medium supplemented with 10% OADC, 0.5% glycerol, 0,025% tyloxapol, and 50 μg/ml kanamycin. Cells were incubated at 37°C without or with 0.18 μM KSK-106 for five generation times until the final OD_600 nm_ = 0.3 was attained. For transposon insertion sequencing, cells were collected, and genomic DNA was extracted as described above. DNA was fragmented and Ilumina P7 adapter with the sequence CAAGCAGAAGACGGCATACGAGAT were ligated using the NeoPrep library prep system (Ilumina). Next, transposon junctions were amplified by using a transposon-specific primer (TCGTCGGCAGCGTCAGATGTGTATAAGAGACAGCCGGGGACTTATCAGCCAACC) and a primer P7 (CAAGCAGAAGACGGCATACGAGAT) using the HotStarTaq master mix kit (Qiagen). The himar1-enriched samples were diluted in a ratio of 1:50. Afterward, amplification was carried out using a p5 indexing primer comprising the sequence AATGATACGGCGACCACCGAGATCTACAC[i5]TCGTCGGCAGCGTC (where [i5] denotes the barcode sequence) and a P7 primer in combination with the HotStarTaq master mix kit from Qiagen. This process added unique barcodes as well as the necessary P5 and P7 flow cell adapter sites required for Illumina sequencing. The PCR protocol employed comprised of an initial denaturation step at 94°C for 3 minutes, followed by a cycle of denaturation at 94°C for 30 seconds, annealing at 55°C for 30 seconds, and extension at 72°C for 30°C. The sequencing was carried out on an Illumina MiSeq system at the University of Minnesota Genomics Center.

After sequencing, transposon and adapter sequences were removed from the 5’ end of the sequencing reads using Cutadapt. ^69^ Furthermore, reads lacking adapter sequences in the 5’ trimming process were discarded. After trimming, all sequence reads started with “TA”. For analysis, all sequences shorter than 18 base pairs were excluded, and a default error rate of 0.1 was applied during the trimming processes. Next, the trimmed sequence reads were aligned to the *M. tuberculosis* H37Ra reference genome (GenBank no. NC_009525.1) using bowtie2.^70^ The alignment permitted a maximum of one base pair mismatch. The genome-mapped sequence reads were printed as a SAM file format, and the count of sequence reads per TA site was determined using the SAMreader_TA script.^71^ These SAM files were subsequently converted into WIG files for further analysis in the TRANSIT software using the resampling method for differentially essentiality analysis.^27^ In short, this analysis calculates the read counts at each gene for each replicate of each condition. The mean read-out in condition A is subtracted from condition B to calculate the observed difference in means. Following this, the TA sites are permuted for a given number of “samples”. or each permutation, we generate a null distribution for the discrepancy in mean read-counts. A p-value was then calculated for the observed discrepancy in mean read-counts P-values were adjusted for multiple testing using the Benjamini-Hochberg procedure.^72^

### RNA-isolation and sequencing for transcriptome analysis

At least 20 mL of *M. tuberculosis* mc^2^6030 cultures were cultivated to the mid-logarithmic phase (OD_600 nm_= 0.5) in 7H9 broth. Triplicate cultures were treated with a lethal concentration (1.9 µM) of KSK-106, whereas control cells were treated with a n equivalent amount of DMSO. After incubation for one generation time, total RNA was isolated. Cells were pelleted by centrifugation and resuspended in 500 µL of Tri Reagent (Invitrogen) plus 1% polyacryl carrier (Molecular Research Center). The samples were transferred to tubes containing 250 µL of 0.1 mm zirconia beads (BioSpec). The samples were bead-beated twice, each time for one minute with a two-minute break on ice between runs. Samples were centrifuged to pellet the beads and the supernatant solution was transferred to fresh tubes. Next, 50 µL of 5-bromo-3-chloro-propane was added to each sample, which were then vortexed and incubated at room temperature for 10 minutes. Following this, samples were centrifuged for 10 minutes at 10,000 rpm at 4°C before the upper aqueous phase was transferred into a fresh tube. Subsequently, 250 µL of isopropanol was added to each sample, they were inverted and incubated at room temperature for 10 min again. The RNA pellets were then formed by centrifuging the samples at 10,000 rpm for an additional 15 minutes at 4°C. The supernatant was removed, and 300 µL of 75% ethanol was added. DNase I Turbo (Invitrogen by Thermo Fisher Scientific) was utilized to remove DNA contaminations following the manufacturer’s protocol. The supernatant was removed after the re-pelleting of RNA, and the RNA was air-dried for 5 to 10 minutes. Thereafter, the RNA was eluted in 50 µL of RNase-free dH_2_O and stored at −80°C for long-term use.

For RNA-seqencing, the Illumina® Stranded Total RNA Prep and Ribo-Zero Plus Ligation kits were employed to convert total RNA into dual-indexed libraries. Briefly, abundant transcripts from total RNA were deleted and bound by specific depletion reagents before the remaining RNA was converted into cDNA throught reverse transcription. Subsequent ligation and amplification steps added adapters for clustering and sequencing on an Illumina system. The libraries were pooled and sequenced on a NovaSeq SP 2×50-bp run. The run generated more than 375 milion pass filter reads with all anticipated barcodes detected and well represented. The mean quality score was ≥Q30f for all libraries. Gel-sizing was done for the libraries, selecting inserts of approximately 200 bps. Raw sequencing reads were quality trimmed (3’ adapter CTGTCTCTTATACACATCT) by using the tool CutAdapt and short reads were eliminated. The trimmed reads were aligned with bowtie2 to the *M. tuberculosis* reference genome NC_000962.3. Normalized read counts were counted using FeatureCounts and log2 transformed with DEseq2 for further analysis, wherein differential gene expression was statistically determined. Subsequently, further analysis was carried out using Rstudio. RNA-sequencing was performed at the University of Minnesota Genomics Center.

### Proteomic profiling employing LC-MS/MS

Cells of *M. tuberculosis* H37Rv were sublethally treated with either 0.05 or 0.2 µM KSK-106, respectively, in 20 mL Middlebrook 7H9 medium supplemented with 0.5% glycerol, 0.2% glucose, 0.085% NaCl and 0.05% tyloxapol and incubated for 10 days. Cells cultivated in Middlebrook 7H9 medium containing an equivalent amount of DMSO were used a solvent control. Cells were centrifuged at 4 °C and washed thrice with PBS (137 mM NaCl; 2.7 mM KCl; 10 mM Na_2_HPO_4_; 1.8 mM KH_2_PO_4_; pH 7.4). Cells were finally resuspended in 2 mL PBS and lysed by bead beating using 100 µm silica-zirconium beads at 50 Hz for 3×3 minutes. Afterwards, 200 µL of a 10% SDS solution was added to each sample, vortexed carefully and incubated for 30 minutes at 4 °C. After centrifugation, the clear supernatant was collected and filter-sterilized thrice through a bacteria-tight 0.2 µM cellulose acetate filter. Protein concentration was measured with BCA assay (Merck Millipore).

For each sample, a volume equivalent to 30 µg of protein was transferred to fresh centrifuge tubes and PBS was added to give a final volume of 115 µL. To eliminate metabolites and non-protein impurities, a methanol-chloroform precipitation was performed. For this purpose, a fourfold excess of methanol was added, followed by 100 µL of chloroform and 300 µL of H_2_O, with thorough mixing in between. To facilitate phase separation, the samples were centrifuged at 9,000 g for 5 min, and the upper aqueous layer was then discarded. The precipitated proteins were washed three times with 300 µL of methanol (sedimentation for 2 min at 9,000 g), and the supernatant was discarded. Subsequently, the dried protein pellet was dissolved in 50 µL of 8 M urea (Cytiva, 17131901) in 100 mM NH_4_HCO_3_ (ABC; Sigma-Aldrich, 11213) and 5 mM dithiothreitol (DTT; Sigma-Aldrich, 11213) was added. After incubating for 40 min with shaking at 1,000 rpm, iodoacetamide (IAA; Sigma-Aldrich, A3221) was added to a final concentration of 20 mM. The mixture was then incubated for an additional 30 min at 37 °C with shaking at 1,000 rpm in the dark and unreacted IAA was afterwards inactivated by addition of DTT to a final concentration of 25 mM. To perform proteolytic digestion, 1 µg of Lys-C (FUJIFILM, 125-05061) was added to each sample, followed by incubation for 3 h at 37 °C with shaking at 1,000 rpm. Subsequently, the urea concentration was reduced to 2 M by the addition of 100 mM ABC. Trypsin (1 µg per sample; Thermo Fisher Scientific, 90057) was added and the digest was continued overnight at 37 °C with shaking at 1,000 rpm. The digest was stopped by addition of formic acid (FA; Fisher Chemical, A11705AMP) to a final concentration of 5% and samples were desalted on self-made C18 StageTips (two discs per tip; 3M, 66883-U) as described before.^73^ For LC-MS/MS analysis, the dry peptides were dissolved in 20 µL 0.1% FA (15 min, 1,500 rpm) and a volume of 5 µL was loaded on a self-packed fused silica capillary tube with integrated pico frit emitter (75 µm ID x 37 cm, 15 µm orifice; New Objectives, PF360-75-15-N-5) filled with ReproSil-Pur 120 C18-AQ (particle size 1.9 µm, Pore Size 120 Å; Dr. Maisch, r119.aq.) material. Peptides were separated using a 140 min gradient generated by an EASY-nLC 1000 liquid chromatography (Thermo Fisher Scientific) with the column heated to 50 °C by a PRSO-V1 column oven (Sonation). For gradient mixture, a rising proportion of acetonitrile (ACN; Honeywell, 14261) with 0.1% FA (solvent B) in H_2_O (Honeywell, 14263) with 0.1% FA (solvent A) was used (7-35% B in A within the first 120 min, 35-80% in the next 10 min, hold for 10 min) at a flow rate of 300 nL/min. Peptides were ionized using a Nanospray Flex ion source (Thermo Fisher Scientific) with 1,800 V spray voltage and MS acquisition was performed in an Orbitrap Elite spectrometer (Thermo Fisher Scientific). MS1 data acquisition was done in a *m/z* range of 300 to 1,800 at a 60,000 orbitrap resolution with a maximal injection time of 50 ns. For data-dependent MS2 acquisition, the 15 most intense MS1 scans were selected with a dynamic exclusion duration of 120 seconds. For precursor isolation, a 2.0 *m/z* quadrupole isolation width was used with subsequent CID fragmentation with a normalized collision energy of 35% and data acquisition at rapid ion trap scan rate with 300% of normalized AGC target. For data processing, MaxQuant 2.3.1.0^74^ was used with the Uniprot proteome UP000001584 as the protein database. ^75^ MaxQuant standard settings were applied with LFQ algorithm and retention time alignment turned on.^76^ For subsequent statistical analysis Perseus 2.0.7.0. was used.^77^ The LFQ data was transformed to the log_2_ scale and missing data points were imputed with values from the lower range of the normal distribution. A Student’s T-test with permutation-based false discovery rate (FDR) with 250 randomizations and an FDR threshold of 0.01 was performed to identify significantly changed protein groups.

### Analysis of KSK cleavage products using LC-MS

Cells of an exponentially growing *M. tuberculosis* H37Rv culture were harvested, washed twice with Middlebrook 7H9 medium containing 0.5% (v/v) glycerol, 0.2% (w/v) glucose and 0.085% (w/v) sodium chloride and resuspended in the same medium to result in an OD_600 nm_ of 1. A final concentration of 100 μM of KSK-104 or KSK-106 or the corresponding volume of DMSO was added to the cells. After 48 h of incubation, a 2.5 mL aliquot was removed from each culture and lysed by bead beating at 50 Hz for 5×3 minutes using 100 μm silica-zirconium beads. The samples were mixed with equal amounts of methanol and incubated for at least one hour at room temperature, before they were centrifugated for 10 minutes at 14,000 rpm. The supernatants were evaporated by freeze-drying in a Savant SpeedVac (Thermo Scientific), and the dried concentrate was solved in 250 μL methanol. The concentrated methanol extracts were measured using an UHR-QTOF maXis 4G (Bruker Daltonics) coupled to an Ultimate 3000 RS UHPLC (Dionex) at the following parameters: Ascentis Express C18 column, 5 cm x 2.1 mm, 2 µm; injection volume 2 µl; solvents: CH_3_CN + 0.1 % FA, H_2_O + 0.1 % FA; flow: 300 µl/min; gradient: 10 % CH_3_CN to 100 % CH_3_CN in 8 min, keep constant for 3 min. ESI-MS was done in positive ion mode with a scan from 50 m/z to 1500 m/z. Measurement was performed at the Center of Molecular and Structural Analytics@Heinrich Heine University (CeMSA@HHU).

### Pharmacokinetic investigations using LC-MS/MS

A liquid chromatography coupled to mass spectrometry (LC-MS/MS) method was developed to determine KSK 104, KSK 106 and their main metabolites. Chromatographic separation was conducted using a Waters Acquity UPLC (Waters, Milford, USA) consisting of a binary pump, the column oven and an autosampler. An Aqua 3u C18 125A (100×2.0mm 3 µm, Phenomenex, Torrance, USA) column was utilized, applying 0.1% formic acid in water and methanol as mobile phase A and B. The flow rate was set to 0.5 mL/min and the following gradient was used: 0-1.5 min: 20% B, 1.4-4.0 min 20-50%, 4-4.5 min 50% B, 4.5-5 min: 50-70% B, 5-5.5 min 70-100% B, 5.5-7 min: 100% B, 7-7.5 min: 100-20% B with a re-equilibration time of 2 min. The column oven was set to 50 °C. Mass spectrometric detection was performed with a Waters Quattro Premier XE in electrospray ionization positive multiple reaction monitoring mode. The capillary voltage was set to 3.5 kV, the source temperature to 135 °C, desolvation temperature to 500 °C, the cone gas flow to 50 L/h, and the desolvation gas flow to 900 L/h. The analyte-specific settings were as follows: KSK-104 376.8 à 180.9 m/z (cone voltage (CV): 18 V, collision energy (CE): 20 V) and KSK-106 386.7 à 190.9 m/z (CV 20 V, CE 20 V). Carvedilol was used as the internal standard with the transition measured 406.8 à 100.0 m/z (CV: 36, CE: 29 V).

*Plasma and whole blood stability:* In vitro plasma/whole blood stability was studied in fresh human EDTA plasma/whole blood at 37 °C. Fresh whole blood/plasma was prewarmed to 37 °C and reactions were started by spiking KSK 104 and KSK 106 to the plasma to a final concentration of 50 ng/mL. Sample aliquots of 100 µL were taken at 0, 120, 240, 360 minutes and after 24 hours. Each aliquot was mixed with 300 µL ice-cold acetonitrile containing the internal standard, and directly vortexed followed by 30 min shaking at 800 rpm at room temperature. Then, samples were centrifuged for 10 min at 13.200 xg. 300 µL of the supernatant was evaporated to dryness under a gentle nitrogen stream and shaking at 450 rpm at 60 °C. Samples were reconstituted in 100 µL 50/50 methanol/water (v/v). The assay was conducted in triplicate. In vitro plasma half-life (t_1/2_) was calculated by t_1/2_=ln2/k_e_, where k_e_ is the slope in the linear fit of the natural logarithm of the fraction remaining of the parent compound vs. incubation time.

### Synthesis of lead structures 7, 8 and analogs 15 – 22

### General procedure 1 (GP 1): EDC-mediated coupling reactions

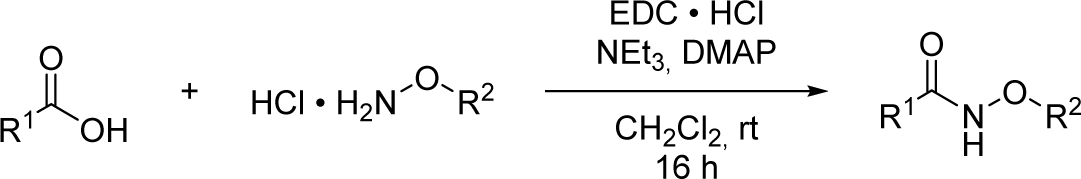

**Method A:** Under nitrogen atmosphere the appropriate *O*-substituted hydroxylamine hydrochloride (1.00 eq.), triethylamine (1.30 eq.) and *N,N*-dimethylpyridin-4-amine (0.10 eq.) were dissolved in dry dichloromethane (20.0 mL/mmol). After stirring for 10 min at room temperature, the corresponding carboxylic acid (1.00 eq.) and 1-ethyl-3-(3-dimethylaminopropyl)carbodiimide hydrochloride (1.25 eq.) were added. The reaction mixture was stirred for 16 h at room temperature.

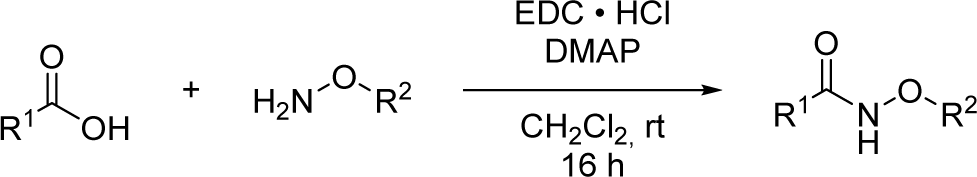

**Method B:** Under nitrogen atmosphere the appropriate *O*-substituted hydroxylamine (1.00 eq.) and *N,N*-dimethylpyridin-4-amine (0.10 eq.) were dissolved in dry dichloromethane (20.0 mL/mmol). After stirring for 10 min at room temperature, the corresponding carboxylic acid (1.00 eq.) and 1-ethyl-3-(3-dimethylaminopropyl)carbodiimide hydrochloride (1.25 eq.) were added. The reaction mixture was stirred for 16 h at room temperature.

**Work-up A:** The reaction mixture was washed three times with saturated sodium bicarbonate solution (20.0 mL/mmol), once with citric acid solution (20.0 mL/mmol) and once with saturated sodium chloride solution (20.0 mL/mmol). The organic layer was dried over anhydrous sodium sulfate, filtered and the solvent was removed under reduced pressure.

**Work-up B:** The reaction mixture was washed three times with (20.0 mL/mmol) saturated sodium bicarbonate solution and once with (20.0 mL/mmol) saturated sodium chloride solution. The organic layer was dried over anhydrous sodium sulfate, filtered and the solvent was removed under reduced pressure.

### N-(2-((benzyloxy)amino)-2-oxoethoxy)-[1,1’-biphenyl]-4-carboxamide: KSK-104 (7)

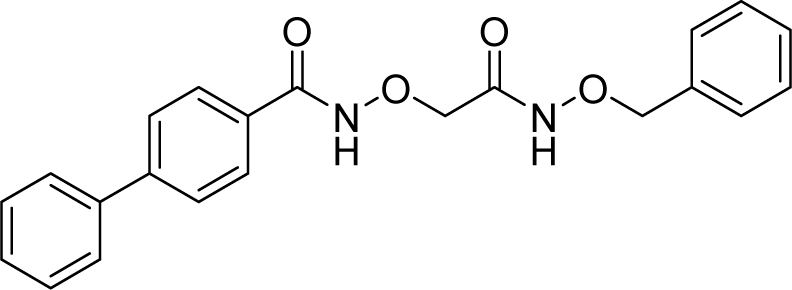

According to **GP 1** (method A, work-up A), KSK-104 (**7**) was synthesized from **6a** and **5** and obtained as a colorless solid (72 % yield). **^1^H-NMR (600 MHz, DMSO-*d*_6_)** δ 12.16 (s, 1H), 11.50 (s, 1H), 7.90 – 7.83 (m, 2H), 7.79 (d, *J* = 8.1 Hz, 2H), 7.76 – 7.70 (m, 2H), 7.50 (t, *J* = 7.7 Hz, 2H), 7.44 – 7.38 (m, 3H), 7.39 – 7.27 (m, 3H), 4.86 (s, 2H), 4.39 (s, 2H).

**^13^C-NMR (151 MHz, DMSO-*d*_6_)** δ 164.91, 164.66, 143.48, 138.95, 135.74, 130.25, 129.06, 128.84, 128.31, 128.21, 127.93, 126.89, 126.69, 77.04, 73.19 **HPLC** *t_R_* = 13.06 min, purity = 96.2 % **Mp.** T_M_ = 151.2 °C

### N-(2-((benzyloxy)amino)-2-oxoethoxy)-4-(pentyloxy)benzamide: KSK-106 (8)

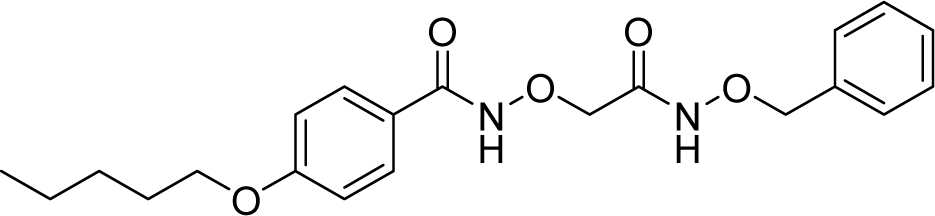

According to **GP 1** (method A, work-up A), KSK-106 (**8**) was synthesized from **6b** and **5** and obtained as a colorless solid (74 % yield).

**^1^H-NMR (600 MHz, DMSO-*d*_6_)** δ 11.93 (s, 1H), 11.52 (s, 1H), 7.76 – 7.70 (m, 2H), 7.36 (m, 5H), 6.98 – 7.02 (m, 2H), 4.85 (s, 2H), 4.36 (s, 2H), 4.01 (t, *J* = 6.54 Hz, 2H), 1.71 (m, 2H), 1.32 – 1.40 (m, 4H), 0.89 (t, *J* = 7.10 Hz, 3H) **^13^C-NMR (151 MHz, DMSO-*d*_6_)** δ 165.14, 164.87, 161.62, 135.74, 129.13, 128.84, 128.31, 123.23, 114.2, 77.03, 73.34, 67.72, 28.23, 27.64, 21.87, 13.9 **HPLC** *t_R_* = 14.60 min, purity ≥ 99.9 % **Mp.** T_M_ = 134.0 °C

### *N*-(2-((benzyloxy)amino)-2-oxoethoxy)-4-(heptyloxy)benzamide (15)

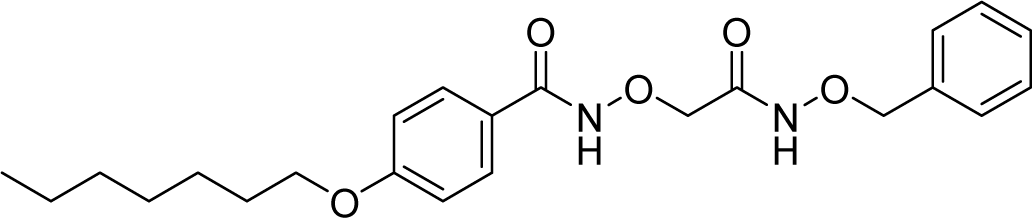

According to **GP 1** (method A, work-up A), KSK-Analog **15** was synthesized from **6c** and **5** and obtained as a colorless solid (49 % yield).

**^1^H-NMR (300 MHz, chloroform*-d*)** δ 11.42 (s, 1H), 9.43 (s, 1H), 7.70 – 7.62 (m, 2H), 7.35 (ddd, *J* = 2.0, 5.9, 33.2 Hz, 5H), 6.93 – 6.86 (m, 2H), 4.95 (s, 2H), 4.48 (s, 2H), 3.98 (t, *J* = 6.6 Hz, 2H), 1.78 (dt, *J* = 6.5, 8.0 Hz, 2H), 1.51 – 1.24 (m, 8H), 0.95 – 0.84 (m, 3H) **^13^C-NMR (75 MHz, chloroform*-d*)** δ 168.23, 166.34, 163.06, 135.12, 129.43, 129.21, 128.75, 128.57, 121.90, 114.64, 78.38, 75.62, 68.43, 31.88, 29.20, 29.15, 26.05, 22.72, 14.21 **HPLC** *t_R_* = 17.29 min, purity = 97.9 % **Mp.** T_M_ = 134.3 °C

### *N*-(2-(tert-butoxyamino)-2-oxoethoxy)-4-(pentyloxy)benzamide (16)

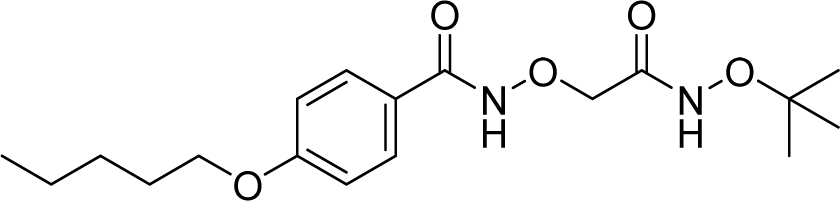

According to **GP 1** (method A, work-up A), KSK-Analog **16** was synthesized from **11** and **14a** and obtained as a yellow oil (55 % yield).

**^1^H-NMR (600 MHz, DMSO*-d_6_*)** δ 11.98 (s, 1H), 10.91 (s, 1H), 7.77 – 7.69 (m, 2H), 7.03 – 6.97 (m, 2H), 4.39 (s, 2H), 4.02 (t, *J* = 6.5 Hz, 2H), 1.76 – 1.68 (m, 2H), 1.43 – 1.30 (m, 4H), 1.17 (s, 9H), 0.89 (t, *J* = 7.2 Hz, 3H) **^13^C-NMR (126 MHz, DMSO*-d_6_*)** δ 165.60, 165.45, 161.57, 128.99, 123.11, 114.15, 80.76, 73.72, 67.66, 28.09, 27.50, 26.14, 21.69, 13.71 **HPLC** *t_R_* = 13.95 min, purity ≥ 99.9 %

### *N*-(2-((benzyloxy)amino)-2-oxoethoxy)-4-(thiazol-2-yl)benzamide (17)

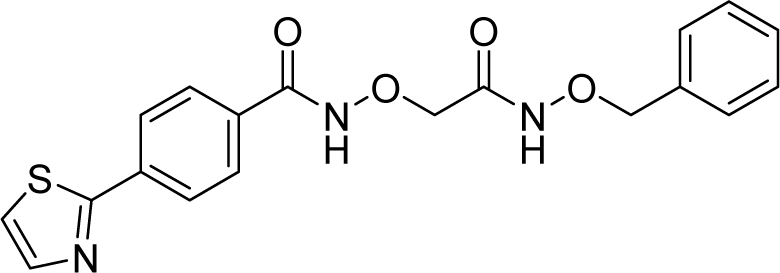

According to **GP 1** (method A, work-up B), KSK-Analog **17** was synthesized from **6d** and **5** and obtained as a colorless solid (18 % yield).

**^1^H-NMR (300 MHz, chloroform*-d*)** δ 11.26 (s, 1H), 10.16 (s, 1H), 8.05 – 7.96 (m, 2H), 7.92 (d, *J* = 3.3 Hz, 1H), 7.86 – 7.73 (m, 2H), 7.44 (d, *J* = 3.3 Hz, 1H), 7.43 – 7.27 (m, 5H), 4.96 (s, 2H), 4.52 (s, 2H) **^13^C-NMR (75 MHz, chloroform*-d*)** δ 167.10, 166.02, 157.60, 143.49, 136.52, 135.14, 131.72, 129.24, 128.83, 128.64, 128.29, 127.06, 120.54, 78.45, 77.36 **HPLC** *t_R_* = 10.50 min, purity ≥ 99.9 % **Mp.** T_M_ = 138.1 °C

### *N*-(2-oxo-2-((pyridin-2-ylmethoxy)amino)ethoxy)-[1,1’-biphenyl]-4-carboxamide (18)

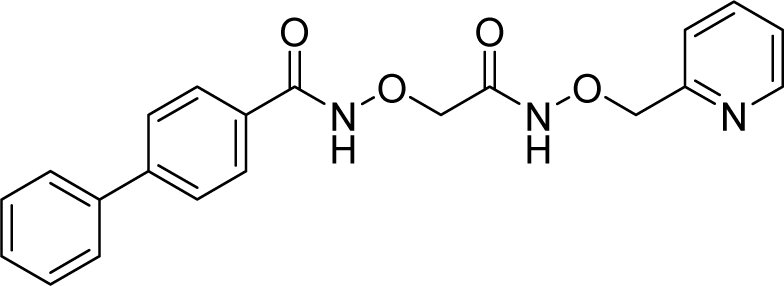

According to **GP 1** (method A, work-up B), KSK-Analog **18** was synthesized from **10** and **14e** and obtained as a colorless solid (63 % yield).

**^1^H-NMR (300 MHz, DMSO-d_6_)** δ 4.40 (s, 2H), 4.96 (s, 2H), 7.34 (ddd, J = 7.6, 4.8, 1.2 Hz, 1H), 7.38 – 7.46 (m, 1H), 7.46 – 7.59 (m, 3H), 7.70 – 7.76 (m, 2H), 7.76 – 7.88 (m, 5H), 8.54 (ddd, J = 5.0, 1.8, 0.9 Hz, 1H), 11.64 (s, 1H), 12.06 (s, 1H) **^13^C-NMR (151 MHz, DMSO)** δ 73.6, 78.3, 123.0, 123.7, 127.1, 127.3, 128.3, 128.7, 129.5, 130.7, 137.2, 139.4, 143.8, 149.5, 156.1, 165.3, 165.5 **HPLC** t*_R_* = 8.77 min, purity = 99 % **Mp.** T_M_ = 152.8 °C

### *N*-(2-oxo-2-(phenoxyamino)ethoxy)-[1,1’-biphenyl]-4-carboxamide (19)

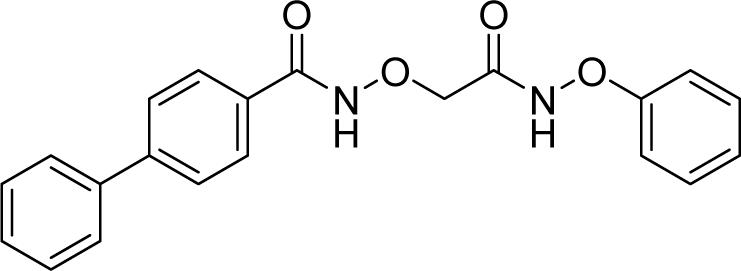

According to **GP 1** (method A, work-up A), KSK-Analog **19** was synthesized from **10** and **14c** and obtained as a colorless solid (20 % yield).

**^1^H-NMR (600 MHz, DMSO-d_6_)** δ 4.59 (s, 2H), 7.01 – 7.10 (m, 3H), 7.28 – 7.36 (m, 2H), 7.39 – 7.45 (m, 1H), 7.50 (t, *J* = 7.7 Hz, 2H), 7.74 (d, 2H), 7.81 (d, *J* = 8.0 Hz, 2H), 7.90 (d, *J* = 7.9 Hz, 2H), 12.25 (s, 2H) **^13^C-NMR (151 MHz, DMSO)** δ 73.4, 113.4, 123.0, 127.2, 127.4, 128.4, 128.7, 129.5, 129.9, 130.7, 139.4, 144.0, 159.7, 165.9, 168.4

**HPLC** t*_R_* = 13.33 min, purity = 96.0 % **Mp.** T_M_ = 135.4 °C

### *N*-(2-((hexyloxy)amino)-2-oxoethoxy)-[1,1’-biphenyl]-4-carboxamide (20)

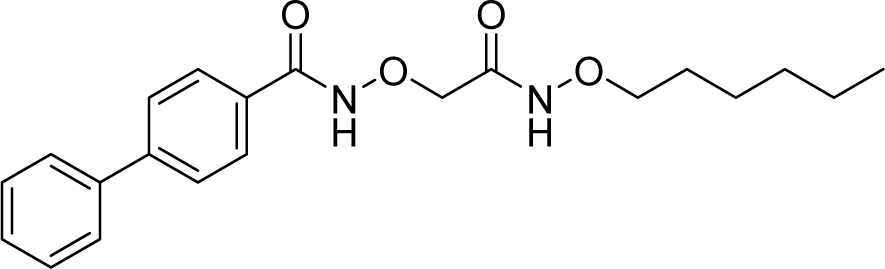

According to **GP 1** (method A, work-up A), KSK-Analog **20** was synthesized from **10** and **14b** and obtained as a colorless solid (74 % yield).

**^1^H-NMR (300 MHz, chloroform-d)** δ 0.80 – 0.91 (m, 3H), 1.20 – 1.46 (m, 6H), 1.61 – 1.74 (m, 2H), 3.95 (t, *J* = 6.8 Hz, 2H), 4.57 (s, 2H), 7.33 – 7.52 (m, 3H), 7.55 – 7.64 (m, 2H), 7.63 – 7.70 (m, 2H), 7.82 – 7.92 (m, 2H), 9.88 (br s, 1H), 11.43 (br s, 1H) **^13^C-NMR (126 MHz, chloroform-d)** δ 14.1, 22.7, 25.5, 28.1, 31.7, 75.8, 77.1, 127.4, 127.6, 128.0, 128.5, 129.0, 129.1, 139.8, 145.8, 166.2, 167.9 **HPLC** t*_R_* = 14.93 min, purity = 97.3 % **Mp.** T_M_ = 102.5 °C

### *N*-((1-((benzyloxy)amino)-1-oxopropan-2-yl)oxy)-[1,1’-biphenyl]-4-carboxamide (21)

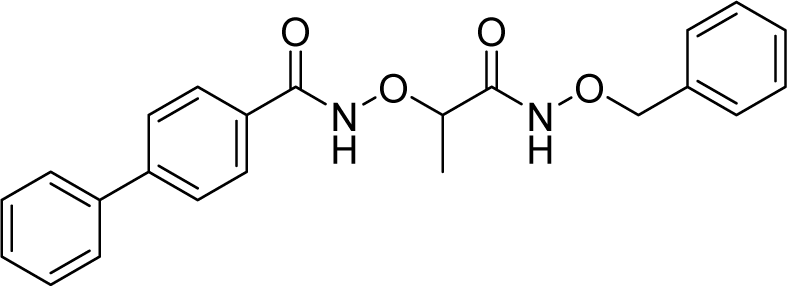

According to **GP 1** (method B, work-up A), KSK-Analog **21** was synthesized from **13** and **2** and obtained as a colorless solid (70 % yield).

**^1^H-NMR (600 MHz, DMSO-d_6_)** δ 1.35 (d, J = 6.7 Hz, 3H), 4.40 (q, J = 6.6 Hz, 1H), 4.84 (q, J = 11.0 Hz, 2H), 7.29 – 7.34 (m, 3H), 7.36 – 7.40 (m, 2H), 7.40 – 7.44 (m, 1H), 7.50 (dd, J = 8.4, 7.0 Hz, 2H), 7.73 (dt, J = 6.3, 1.3 Hz, 2H), 7.79 (d, J = 8.0 Hz, 2H), 7.86 (d, J = 8.2 Hz, 2H), 11.45 (s, 1H), 12.02 (s, 1H) **^13^C-NMR (151 MHz, DMSO)** δ 17.2, 77.4, 79.2, 127.1, 127.3, 128.4, 128.6, 128.7, 128.8, 129.4, 129.5, 130.9, 136.2, 139.5, 143.9, 165.4, 168.1 **HPLC** t*_R_* = 13.33 min, purity = 97.9 % **Mp.** T_M_ = 177.8 °C

### *N*-(2-((cyclohexylmethoxy)amino)-2-oxoethoxy)-[1,1’-biphenyl]-4-carboxamide (22)

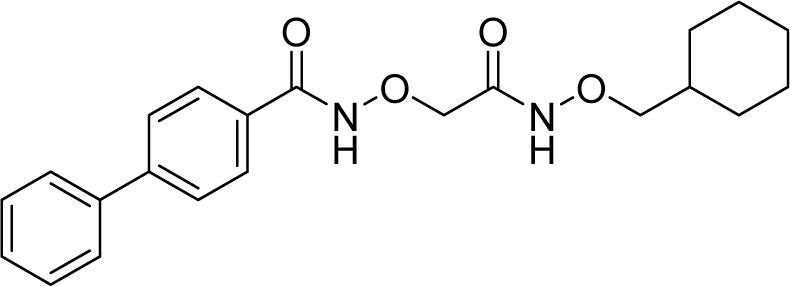

According to **GP 1** (method A, work-up A), KSK-Analog **22** was synthesized from **10** and **14d** and obtained as a colorless solid (94 % yield).

**^1^H-NMR (600 MHz, DMSO-d_6_)** δ 0.88 – 0.99 (m, 2H), 1.06 – 1.25 (m, 3H), 1.56 – 1.69 (m, 4H), 1.69 – 1.77 (m, 2H), 3.62 (d, *J* = 6.7 Hz, 2H), 4.39 (s, 2H), 7.39 – 7.46 (m, 1H), 7.50 (t, *J* = 7.7 Hz, 2H), 7.70 – 7.75 (m, 2H), 7.79 (d, *J* = 8.1 Hz, 2H), 7.87 (d, 2H), 11.37 (s, 1H), 12.14 (s, 1H) **^13^C-NMR (151 MHz, DMSO)** δ 25.6, 26.5, 29.6, 36.4, 73.7, 81.1, 127.2, 127.3, 128.3, 128.7, 129.5, 130.7, 139.4, 143.9, 164.8, br s 165.3 **HPLC** t*_R_* = 14.93 min, purity = 99 % **Mp.** T_M_ = 115.6 °C

## Supporting information

KSK Supplementary Information

## Acknowledgement

Financial support for this study was provided to R.K. by the German Research Foundation (Deutsche Forschungsgemeinschaft, DFG, project number KA 2259/4-1) and by the Jürgen Manchot Stiftung (graduate school MOI IV). T.K. acknowledges support from the DFG (project number KU 1577/3-1). Thanks to the CeMSA@HHU (Center for Molecular and Structural Analytics @ Heinrich Heine University) for recording some of the mass-spectrometric data.

## Author contributions

Conceptualization, funding acquisition, and supervision, T.K., R.K; microbiological investigation, K.V., L.v.G., A-L.K-D.; chemical synthesis, O.M., A.B., K.S., B.L.; proteome analysis, D.P., F.Ka.; Tn-seq analysis, T.A.C., M.D.H., L.O., Z.J.; pharmacokinetic investigations, B.B.; data analysis, F.Ko., T.R.I., M.K., A.D.B.; writing – original draft, K.V., L.v.G., O.M., T.K., R.K..

