## Supplementary material for "*α*-Aminooxyacetic acid derivatives acting as pro-drugs against *Mycobacterium tuberculosis*": KSK Supplementary Information

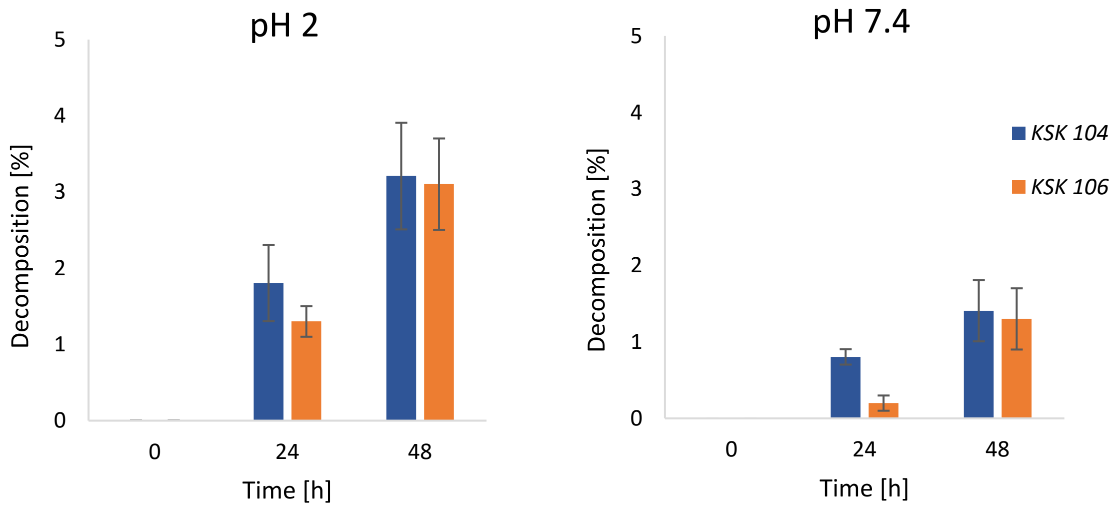


**Figure S1. Stability testing of KSK-104 and KSK-106 in aqueous media.** Both compounds were dissolved (0.5 mg/mL) in a vehicle of 90% PBS (pH 2 and pH 7.4), 7% Tween® 80 and 3% ethanol (v/v). These solutions were shaken at 37 °C for 48 h, and the compound decomposition was evaluated by HPLC analysis. Values represent means of duplicate measurements (n = 2) ± standard deviation.

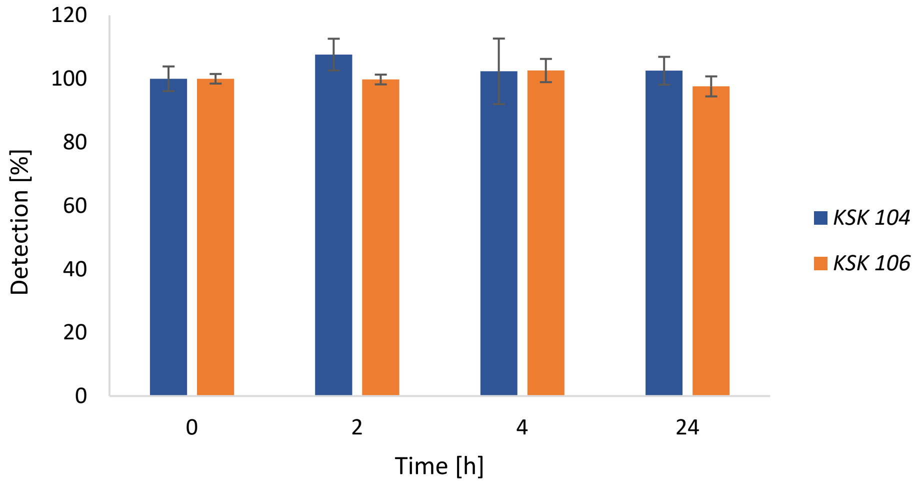


**Figure S2. Stability testing of KSK 104 and KSK 106 in human EDTA-plasma.** Both compounds were dissolved in human EDTA-plasma (50 ng/mL) and constantly incubated at 37 °C for 24 h. Detection was performed via LC-MS/MS analysis. Values represent means of duplicate measurements (n = 2) ± standard deviation.

A

| Strain | KSK-104 | KSK-106 |
|---|---|---|
| *M. bovis* BCG Pasteur | 0.78 | 0.39 |
| *M. smegmatis* $mc^2155$ | >100 | >100 |
| *M. abscessus* (CF001s) | >100 | >100 |
| *M. marinum* DSM 44344 | >100 | >100 |
| *Staphylococcus aureus* Mu50 | >100 | >100 |
| *Acinetobacter baumannii* ATCC BAA-1605 | >100 | >100 |
| *Pseudomonas aeruginosa* ATCC 27853 | >100 | >100 |

B

| Cell line | Tissue | $IC_{50}$ [µM] |
|---|---|---|
| MRC-5 | Lung fibroblasts | > 100 |
| THP-1 | Monocytes | > 100 |
| HEPG2 | Liver | > 100 |
| HUH7 | Liver | > 100 |
| CLS-54 | Lung | > 100 |
| HEK293 | Kidney | > 100 |
| H4 | Brain | > 100 |
| SH-SY5Y | Neuroblasts | > 100 |

**Figure S3. Antibacterial activity and cytotoxicity profile of KSK molecules. A**) KSK-104 and KSK-106 have been tested in microbroth dilution assays against various mycobacteria and nosocomial bacteria. Concentrations of $MIC_{90}$ values are given in µM. **B)** Cytotoxicity of KSK-106 against various human cell lines *in vitro*. $IC_{50}$ values are given in µM. Growth in **A** and **B** was quantified employing the resazurin reduction assay. Measurements were performed in triplicates revealing no deviations in the reported $MIC_{90}$ and $IC_{50}$ values.

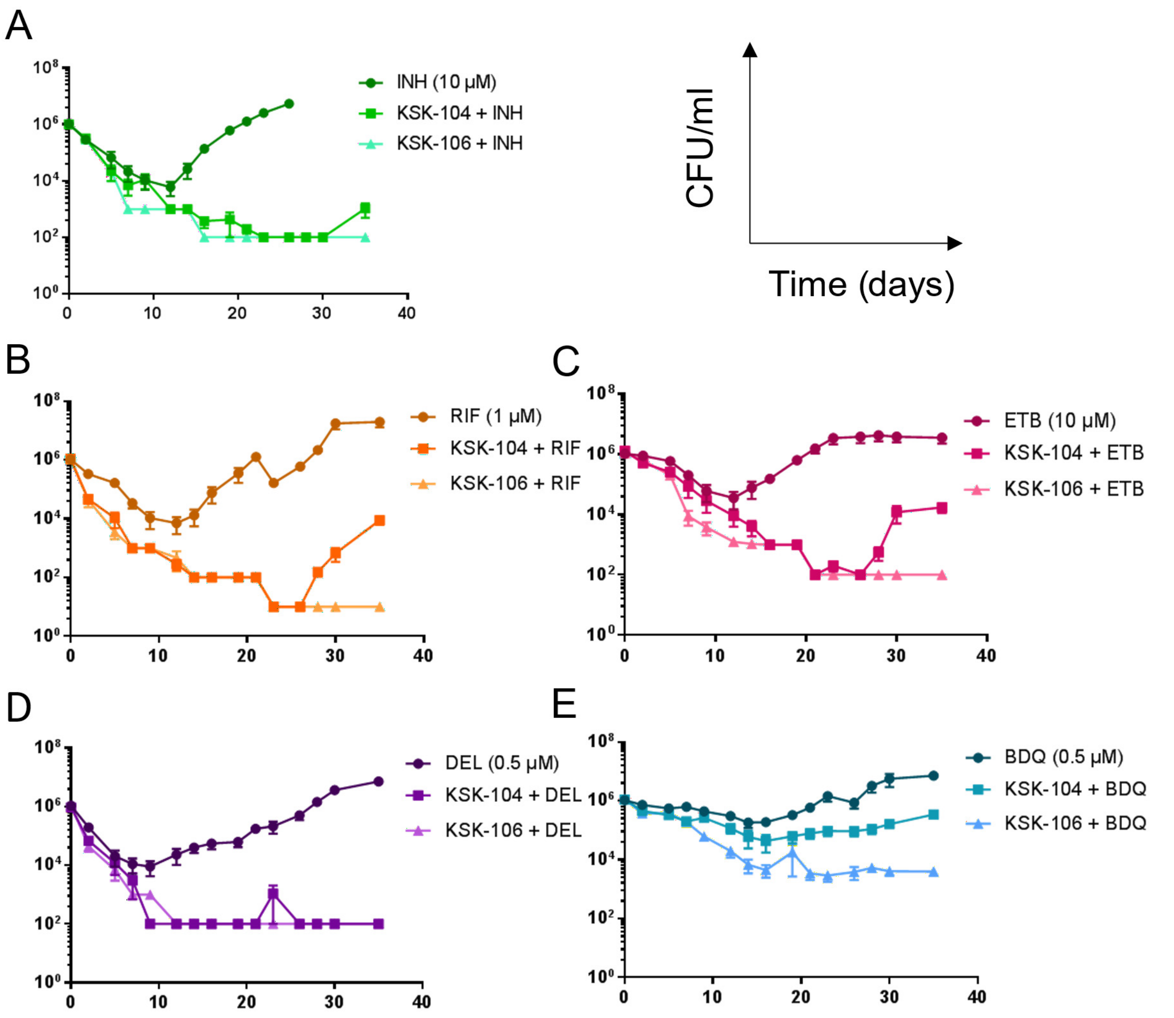


**Figure S4. *In vitro* activity of KSK-104 and KSK-106 in combination with different first- and second-line antibiotics.** *M. tuberculosis* H37Rv was treated with the indicated concentrations of clinically used drugs in monotherapy and in combination with 0.25 µM KSK-104 or KSK-106, respectively. Combination with isoniazid (INH, **A**), rifampicin (RIF, **B**), ethambutol (ETB, **C**) and delamanid (**D**) led to additive effects concerning the killing efficacy and delayed resurgence of bacteria surviving the initial treatments. In contrast, combination with bedaquiline (BDQ, **E**) resulted in antagonistic effects. Experiments have been performed in triplicates. The limit of detection was 10 CFU/mL in **B** and 100 CFU/mL in **A** and **C**-**E**. CFU, colony forming units.

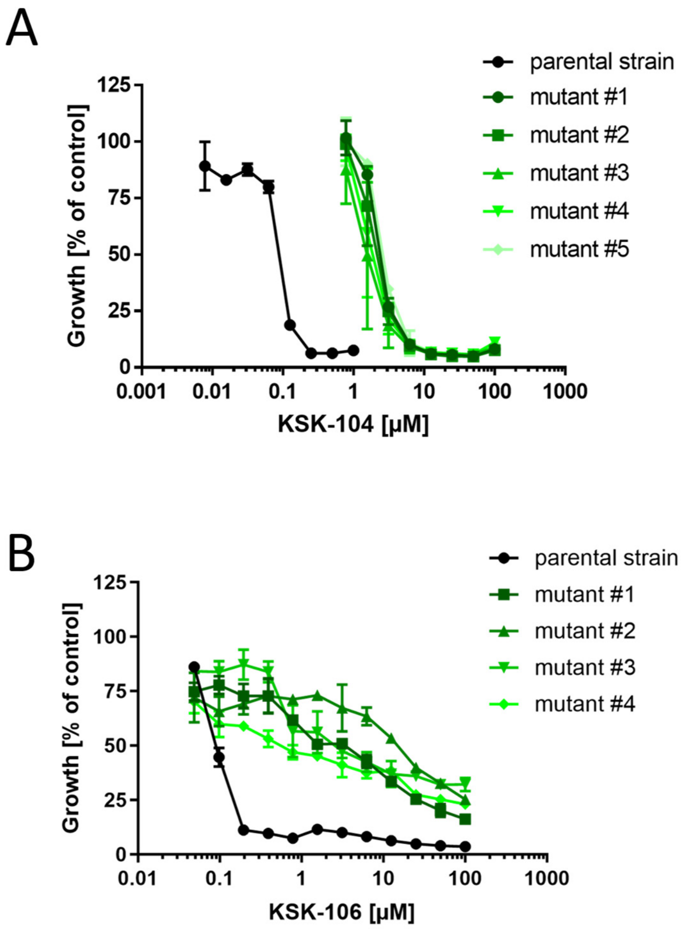


**Figure S5. Spontaneous single-step resistant mutants isolated against KSK-104 and KSK-106.** SRMs were generated against KSK-104 (**A**) and KSK-106 (**B**). MICs were tested in microbroth dilution assays. Growth in **A** and **B** was quantified employing the resazurin reduction assay. Data shown as means of triplicates with SD. The mutations that were found during the whole-genome-sequencing in the respective mutants are listed in the tables.

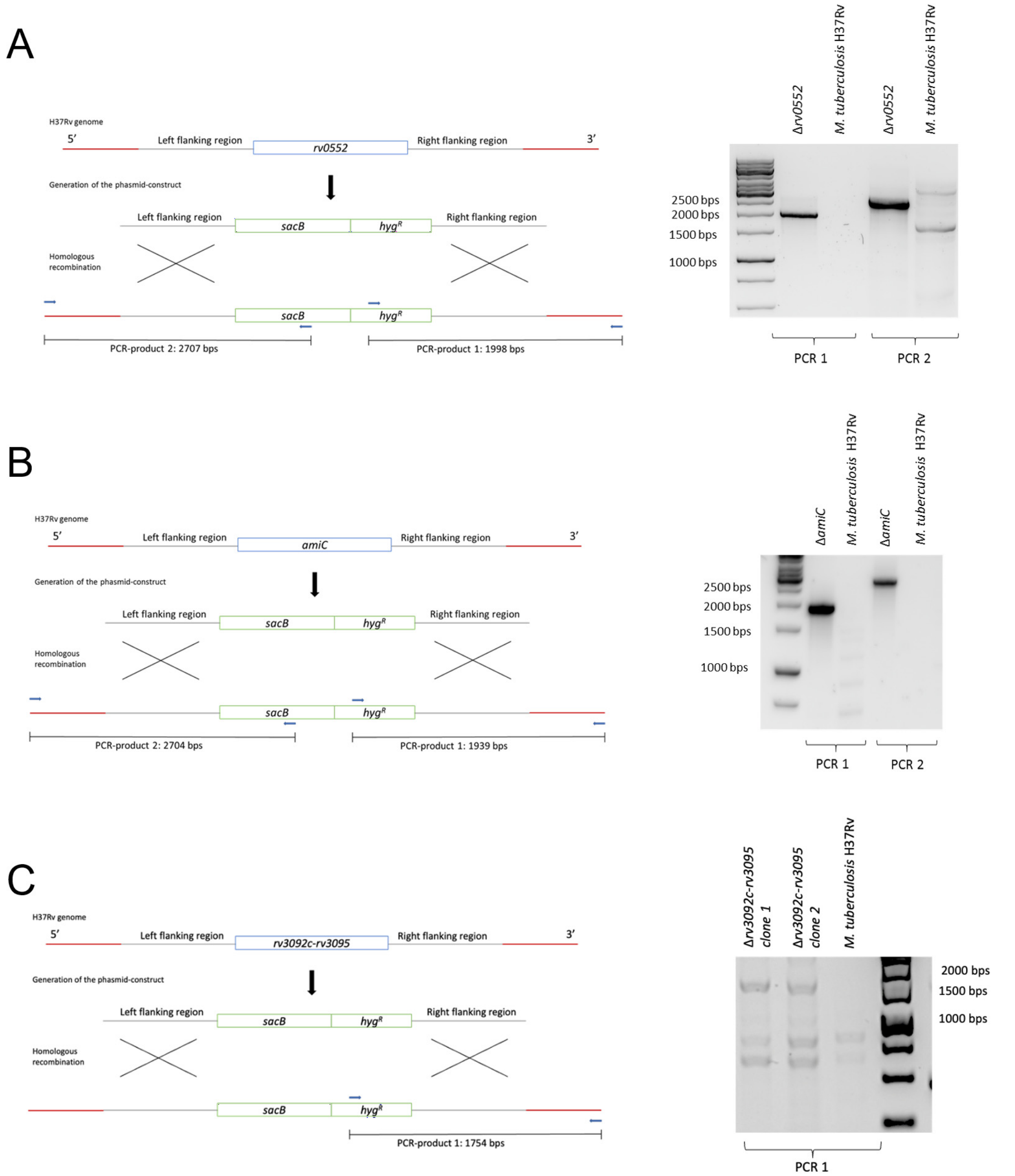


**Figure S6. Generation of site-specific gene deletion mutants using specialized transduction.** Organization of the respective loci in *M. tuberculosis* H37Rv wild type and **A)** the *Rv0552*, **B)** *amiC* and **C)** *Rv3092c-Rv3095* gene deletion mutants (left panels). The location of flanking regions used to construct the allelic exchange substrate as well as of primers used in diagnostic PCR are indicated. The sizes of the relevant diagnostic PCR products for verification of gene disruption are shown. Diagnostic PCR to verify gene deletion mutants was performed (right panels). **A)** PCR product 1 (2,707 bp) was produced using forward primer binding 1,000 bps upstream of *Rv0552* and reverse primer binding in the *sacB* gene as indicated. PCR product 2 (1,998 bp) was produced using forward primer binding in the hygromycin resistance (*hyg*$^{R}$) gene and reverse primer binding 1,000 bps downstream of *Rv0552*. B) PCR product 1 (2,704 bp) was produced using forward primer binding 1,000 bps upstream of *amiC* and reverse primer binding in the *sacB* gene as indicated. PCR product 2 (1,939 bp) was produced using forward primer binding in the hygromycin resistance (*hyg*$^{R}$) gene and reverse primer binding 1,000 bps downstream of *amiC*. **C)** PCR product (1,754 bp) was produced using forward primer binding in the hygromycin resistance (*hyg*$^{R}$) gene and reverse primer binding 1,000 bps downstream of *rv3095*.

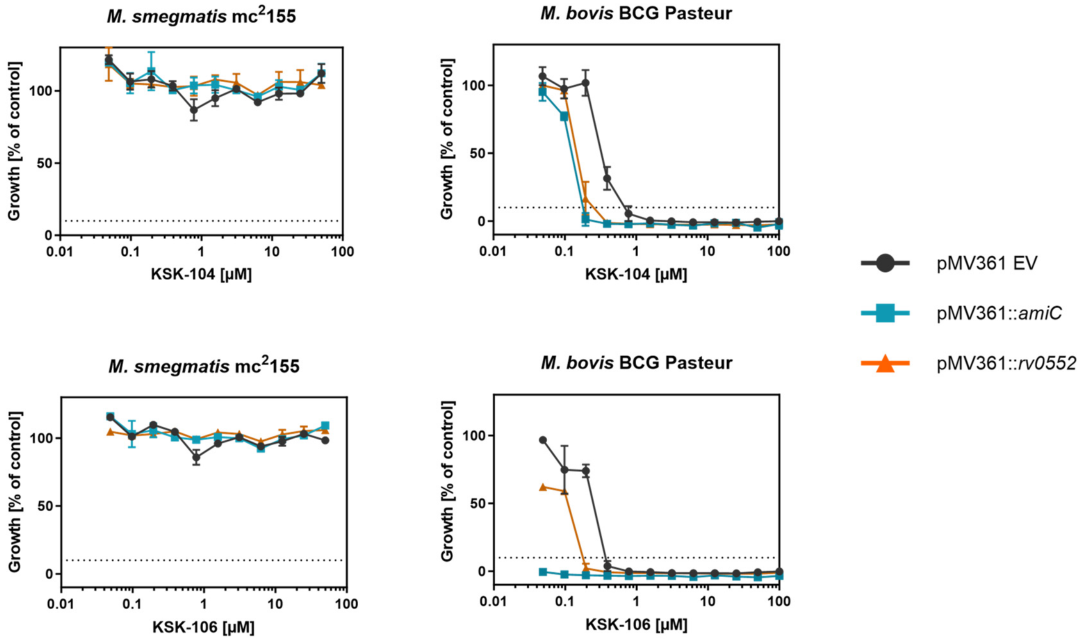


**Figure S7. Overexpression of *amiC* or *Rv0552* leads to increased sensitivity of *M. bovis*, but not *M. smegmatis*, towards KSK-104 and KSK-106.** Dose-response curves for KSK-104 (top) and KSK-106 (bottom) showing a concentration-dependent growth inhibition of recombinant strains of *M. bovis* BCG Pasteur harboring the empty vector control pMV361::EV or the overexpression constructs pMV361::*amiC* or pMV361::*Rv0552,* respectively*,* leading to an increased susceptibility of the cells towards the KSKs. In contrast, there was no sensitivity of *M. smegmatis* mc$^2$155 cells towards the KSKs even during overexpression of *Rv0552* or *amiC*. Data shown as means of duplicates ± SD. Growth was quantified employing the resazurin reduction assay. The dashed lines indicate 10% residual growth.

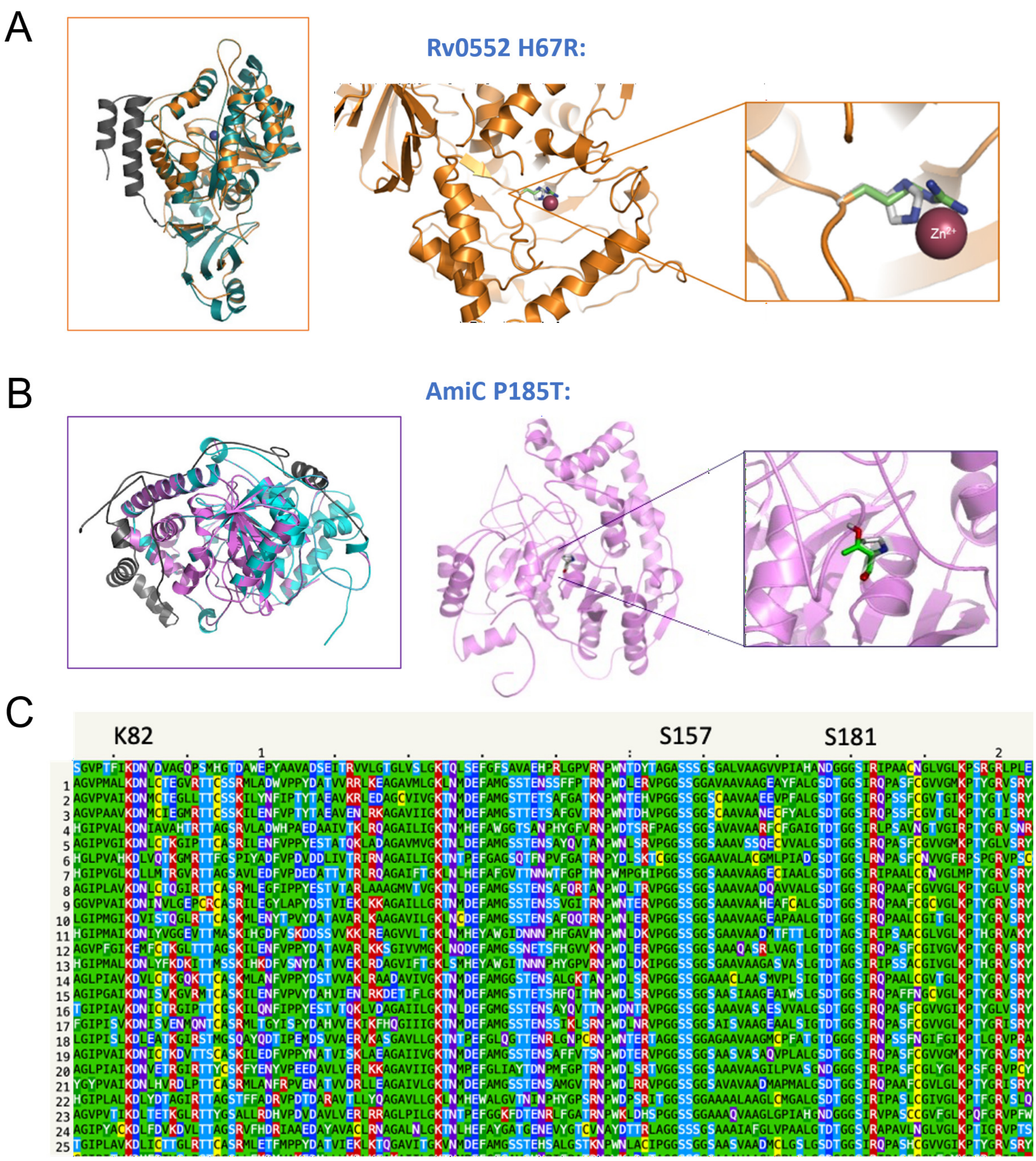


**Figure S8. Structural features of the putative amidohydrolases Rv0552 and AmiC. A)** Structural homology modeling using Phyre2 revealed that mutations in the gene Rv0552 observed in the spontaneously KSK-104-resistant mutants are likely causing an altered interaction with Zn$^{2+}$ atoms resulting in destabilized and inactive proteins. **B**) Structural homology modeling of AmiC indicating that the observed mutation P195T leads to a loss of rigidity inside the hydrophobic core resulting in a non-accessibility of the active center. **C)** *PSI-Blast Pseudo*-Multiple sequence alignment proposes that AmiC belongs to the amidase signature superfamily. Within this family, Ser-*cis*-Ser-Lys triads have been proposed to form the catalytic center with the specific residues for AmiC expected to be Ser$^{181}$-*cis*-Ser$^{157}$-Lys$^{82}$. Identities of the labeled sequences: 1) UniRef50_Q2RGY4, 2) UniRef50_C4ZHB9, 3) UniRef50_UPI0001C3563A, 4) UniRef50_B1G4X5, 5) UniRef50_B2IYD7, 6) UniRef50_A5USQ6, 7) UniRef50_D1CAY8, 8) UniRef50_Q2JL51, 9) UniRef50_B3DWT4, 10) UniRef50_A7NKM0, 11) UniRef50_D2KYA7,

12) UniRef50_Q6MRL7, 13) UniRef50_B1HYA7, 14) UniRef50_D2R415, 15) UniRef50_D2RM15, 16) UniRef50_Q113L8, 17) UniRef50_B9DWL8, 18) UniRef50_B3T355, 19) UniRef50_B2A5W7, 20) UniRef50_O28325, 21) UniRef50_A3EU19, 22) UniRef50_A5UXU3, 23) UniRef50_D0MDU0, 24) UniRef50_B9QYT0, 25) UniRef50_A8ZSP8. Residues are coloured by properties: bright-green = hydrophobic, bright-blue = negative charge, dark-green = large hydrophobic, yellow = cysteine, bright-red = positive charge, purple = polar, dull-blue = small alcohol.

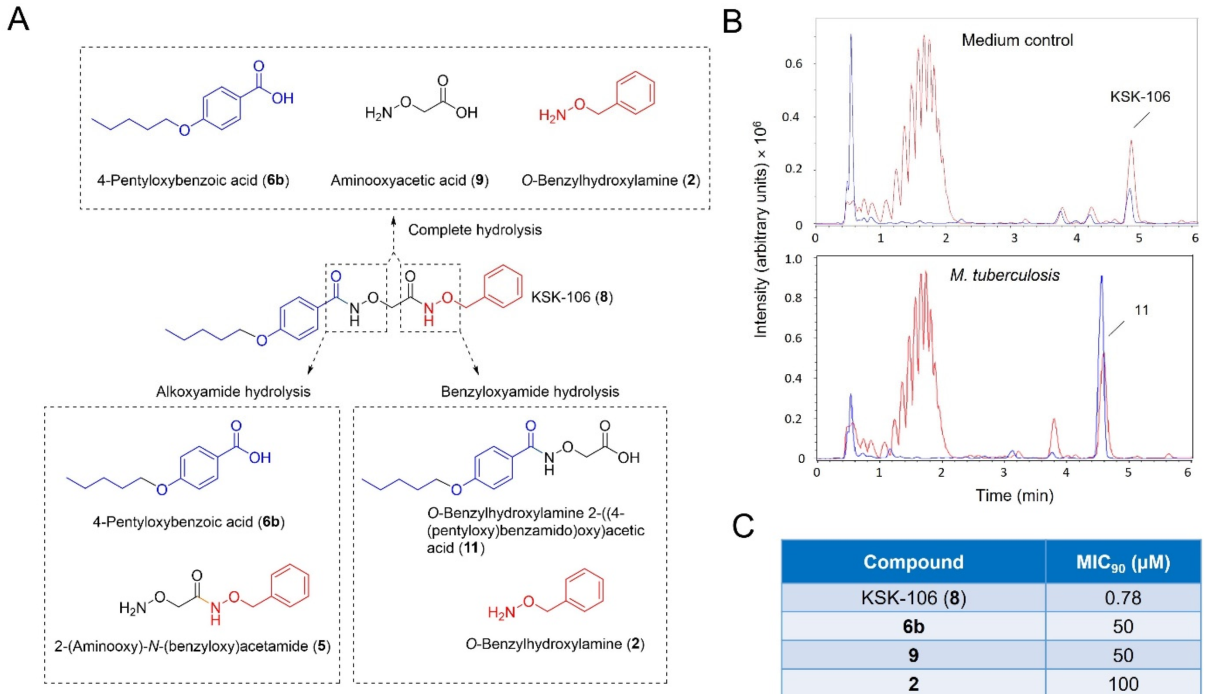


C

| Compound | $MIC_{90}$ (µM) |
|---|---|
| KSK-106 (**8**) | 0.78 |
| **6b** | 50 |
| **9** | 50 |
| **2** | 100 |
| **5** | 50 |
| **11** | 25 |
| **6b** + **9** + **2** | 50 |
| **6b** + **5** | 100 |
| **11** + **2** | 50 |

**Figure S9. Hydrolytic pro-drug activation of KSK-106 by Rv0552 and AmiC. A)** Structure of potential hydrolysis products released from KSK-106 by amidohydrolases AmiC and Rv0552. **B)** Qualitative ESI-LC-MS analysis of methanol extracts obtained after 48 h incubation of 100 µM KSK-106 in sterile 7H9 medium (top) or in 7H9 medium inoculated with *M. tuberculosis* H37Rv cells (bottom). Scan was from 50 to 1500 m/z in positive mode. Base peak chromatogram (+all MS) is shown in red, UV chromatogram at 254 nm is shown in blue. Identified peaks: KSK-106 $[m/z + H]^+$ = 387.19, **11** $[m/z + H]^+$ = 209.12. **C)** $MIC_{90}$ values of potential KSK-106 hydrolysis products against *M. tuberculosis* H37Rv. Compounds were tested individually and in various combinations. For combination treatments, equimolar mixtures were used containing each compound at the indicated concentration.

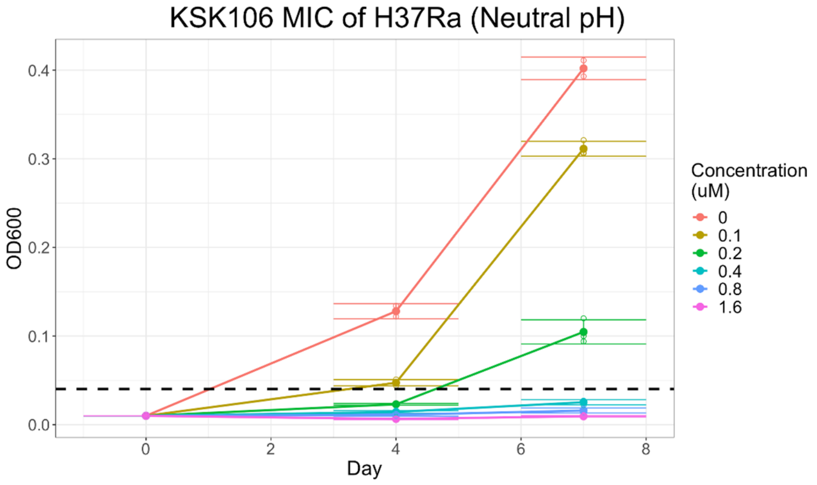


**Figure S10. Growth of *M. tuberculosis* H37Ra treated with different KSK-106 concentrations.** *M. tuberculosis* H37Ra cells were inoculated with a starting $OD_{600nm}$ of 0.01 from a growing culture in the exponential phase. Cells were treated in triplicates with different concentrations of KSK-106 as indicated in the legend. $OD_{600\ nm}$ was measured after 4 and 7 days. Data shown as means of triplicates with SD. From these data, a concentration of 0.18 µM was estimated to result in ca. 50% growth inhibition after 5 generations.

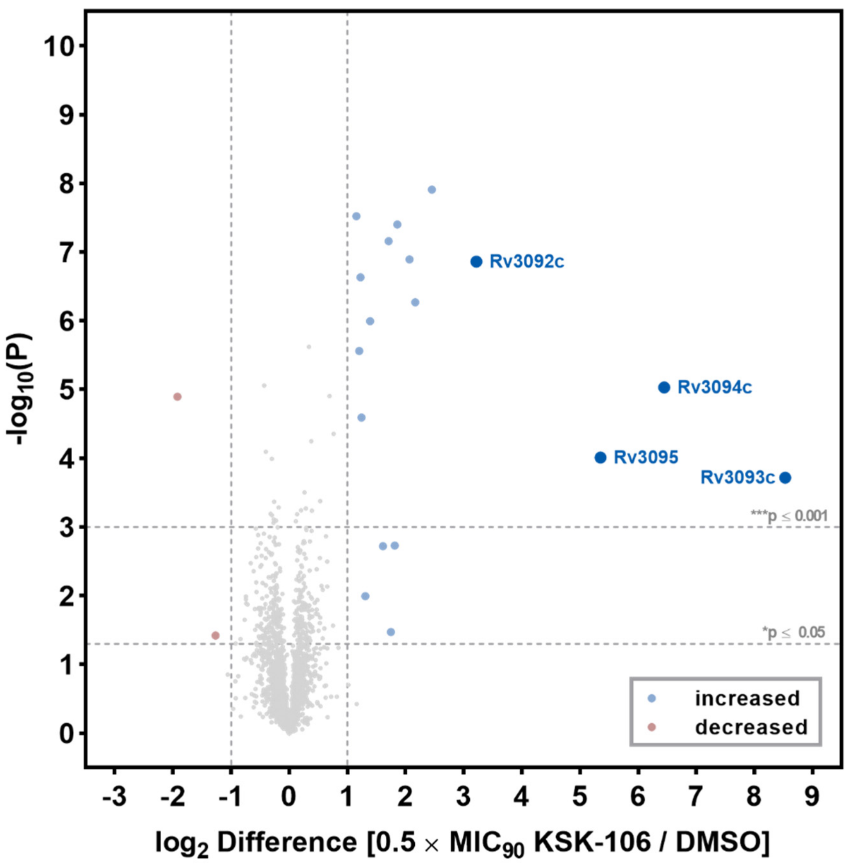


**Figure S11. LC-MS/MS-based whole protein analysis of silenced cells of *M. tuberculosis* H37Rv treated with a sublethal concentration of KSK-106 (0.2 µM, corresponding to 0.5 × $MIC_{90}$) compared to DMSO control.** The volcano plot illustrates the $log_2$-fold change in abundance in KSK-106 treated vs. non-treated cells (X-axis) and corresponding -$log_{10}$ p values (Y-axis). Proteins complying with the chosen threshold of significance and showing a $log_2$-fold change $\geq 1$ or $\leq -1$ are marked in blue or red, respectively. Quantification was done via label free quantification (LFQ) of four to five replicates per sample group. To identify statistically significant hits from the analysis, $P \leq 0.05$ (Student's T-test; permutation-based FDR with 250 randomizations and FDR = 0.01) was applied.

**Table S1 Transposon insertions in genes resulting in apparent fitness changes of transposon mutants of *M. tuberculosis* H37Ra cells during KSK-106 treatment.** Genes complying with the chosen threshold of significance $P_{adj}<0.05$ are listed. $P_{adj}$ values were rounded to 3 decimal points. Negative $log_2FC$ values indicate underrepresentation, positive values overrepresentation of the mutants in the pool, demonstrating aggravating or alleviating effects of insertions, respectively.

| #Orf | Name | Description | $log_2FC$ | Adj. p-value |
|---|---|---|---|---|
| **Rv2888c** | amiC | Probable amidase AmiC (aminohydrolase) | 1 | 0.000 |
| **Rv0552** | Rv0552 | Conserved protein/putative amidohydrolase | 0.94 | 0.000 |
| **Rv3696c** | glpK | Probable glycerol kinase GlpK (ATP:glycerol 3-phosphotransferase) (glycerokinase) (GK) | 0.91 | 0.000 |
| **Rv0544c** | Rv0544c | Possible conserved transmembrane protein | 0.88 | 0.000 |
| **Rv0545c** | pitA | Probable low-affinity inorganic phosphate transporter integral membrane protein PitA | 0.87 | 0.000 |
| **Rv0111** | Rv0111 | Possible transmembrane acyltransferase | 0.86 | 0.000 |
| **Rv2241** | aceE | Pyruvate dehydrogenase E1 component AceE (pyruvate decarboxylase) (pyruvate dehydrogenase) | 0.75 | 0.000 |
| **Rv0546c** | Rv0546c | Conserved protein/putative carbon-sulfur lyase | 0.62 | 0.000 |
| **Rv2932** | ppsB | Phenolpthiocerol synthesis type-I polyketide synthase PpsB | 0.59 | 0.000 |
| **Rv2935** | ppsE | Phenolpthiocerol synthesis type-I polyketide synthase PpsE | 0.57 | 0.000 |
| **Rv2930** | fadD26 | Fatty-acid-AMP ligase FadD26 (fatty-acid-AMP synthetase) (fatty-acid-AMP synthase) | 0.55 | 0.000 |
| **Rv3005c** | doxX | Probable membrane oxidoreductase component (MRC) DoxX | 0.55 | 0.000 |
| **Rv3484** | cpsA | Possible conserved protein CpsA | 0.54 | 0.000 |
| **Rv2941** | fadD28 | Fatty-acid-AMP ligase FadD28 (fatty-acid-AMP synthetase) (fatty-acid-AMP synthase) | 0.52 | 0.000 |
| **Rv2933** | ppsC | Phenolpthiocerol synthesis type-I polyketide synthase PpsC | 0.51 | 0.007 |
| **Rv2931** | ppsA | Phenolpthiocerol synthesis type-I polyketide synthase PpsA | 0.46 | 0.000 |
| **Rv0806c** | cpsY | Possible UDP-glucose-4-epimerase CpsY (galactowaldenase) (UDP-galactose-4-epimerase) (uridine diphosphate galactose-4-epimerase) | 0.45 | 0.000 |
| **Rv3066** | Rv3066 | Probable transcriptional regulatory protein (probably DeoR-family) | 0.43 | 0.007 |
| **Rv0678** | mmpR5 | MarR-like transcriptional regulator | 0.42 | 0.007 |
| **Rv1328** | glgP | Probable glycogen phosphorylase GlgP | 0.41 | 0.000 |
| **Rv2940c** | mas | Probable multifunctional mycocerosic acid synthase membrane-associated Mas | 0.36 | 0.007 |
| **Rv3717** | Rv3717 | N-acetylmuramyl-L-alanine amidase | 0.36 | 0.000 |
| **Rv1213** | glgC | Glucose-1-phosphate adenylyltransferase GlgC (ADP-glucose synthase) (ADP-glucose pyrophosphorylase) | 0.35 | 0.000 |
| **Rv2458** | mmuM | Probable homocysteine S-methyltransferase MmuM (S-methylmethionine:homocysteine methyltransferase) (cysteine methyltransferase) | 0.34 | 0.007 |
| **Rv0172** | mce1D | Mce-family protein Mce1D | 0.33 | 0.000 |
| **Rv0470c** | pcaA | Mycolic acid synthase PcaA (cyclopropane synthase) | 0.33 | 0.007 |
| **Rv2721c** | Rv2721c | Antigen | 0.33 | 0.000 |
| **Rv0554** | bpoC | Possible peroxidase BpoC (non-haem peroxidase) | 0.31 | 0.000 |
| **Rv2115c** | mpa | Mycobacterial proteasome ATPase Mpa | 0.31 | 0.000 |
| **Rv0503c** | cmaA2 | Cyclopropane-fatty-acyl-phospholipid synthase 2 CmaA2 (cyclopropane fatty acid synthase) (CFA synthase) (cyclopropane mycolic acid synthase 2) (mycolic acid trans-cyclopropane synthetase) | 0.3 | 0.000 |
| **Rv2914c** | pknI | Probable transmembrane serine/threonine-protein kinase I PknI (protein kinase I) (STPK I) (phosphorylase B kinase kinase) (hydroxyalkyl-protein kinase) | 0.3 | 0.000 |
| **Rv0169** | mce1A | Mce-family protein Mce1A | 0.29 | 0.000 |

**Table S1 continued**

| #Orf | Name | Description | $log_2FC$ | Adj. p-value |
|---|---|---|---|---|
| **Rv0465c** | lrpI | Transcriptional regulatory protein%2C local regulatory protein of icl1 | 0.29 | 0.007 |
| **Rv3045** | adhC | Probable NADP-dependent alcohol dehydrogenase AdhC | 0.25 | 0.035 |
| **Rv0019c** | fipA | FtsZ-interacting protein A%2C FipA | 0.24 | 0.014 |
| **Rv1195** | PE13 | PE family protein PE13 | 0.24 | 0.025 |
| **Rv1809** | PPE33 | PPE family protein PPE33 | 0.24 | 0.020 |
| **Rv2585c** | Rv2585c | Possible conserved lipoprotein/putative ATP-binding cassette (ABC) transport protein (oligopeptide transport) | 0.23 | 0.000 |
| **Rv1220c** | Rv1220c | Probable methyltransferase | 0.22 | 0.025 |
| **Rv1196** | PPE18 | Surfaced-exposed antigen | 0.21 | 0.014 |
| **Rv1808** | PPE32 | PPE family protein PPE32 | 0.21 | 0.000 |
| **Rv0483** | lprQ | Probable conserved lipoprotein LprQ | -0.17 | 0.000 |
| **Rv2048c** | pks12 | Polyketide synthase Pks12 | -0.17 | 0.007 |
| **Rv1364c** | Rv1364c | Possible sigma factor regulatory protein | -0.19 | 0.025 |
| **Rv3720** | Rv3720 | Possible fatty acid synthase | -0.21 | 0.031 |
| **Rv2203** | Rv2203 | Possible conserved membrane protein | -0.24 | 0.007 |
| **Rv2936** | drrA | Daunorubicin-dim-transport ATP-binding protein ABC transporter DrrA | -0.25 | 0.000 |
| **Rv3010c** | pfkA | Probable 6-phosphofructokinase PfkA (phosphohexokinase) (phosphofructokinase) | -0.28 | 0.014 |
| **Rv3719** | Rv3719 | Conserved protein/putative amide-bond oxidoreductase | -0.28 | 0.000 |
| **Rv2937** | drrB | Daunorubicin-dim-transport integral membrane protein ABC transporter DrrB | -0.31 | 0.007 |
| **Rv3057c** | Rv3057c | Probable short chain alcohol dehydrogenase/reductase | -0.32 | 0.000 |
| **Rv1813c** | Rv1813c | Conserved hypothetical protein | -0.33 | 0.035 |
| **Rv2607** | pdxH | Probable pyridoxamine 5'-phosphate oxidase PdxH (PNP/PMP oxidase) (pyridoxinephosphate oxidase) (PNPOX) (pyridoxine 5'-phosphate oxidase) | -0.36 | 0.040 |
| **Rv2942** | mmpL7 | Conserved transmembrane transport protein MmpL7 | -0.39 | 0.000 |
| **Rv3200c** | Rv3200c | Possible transmembrane cation transporter | -0.39 | 0.000 |
| **Rv1040c** | PE8 | PE family protein PE8 | -0.41 | 0.035 |
| **Rv1580c** | Rv1580c | Probable PhiRv1 phage protein | -0.42 | 0.007 |
| **Rv3134c** | Rv3134c | Universal stress protein family protein | -0.42 | 0.000 |
| **Rv3644c** | Rv3644c | Possible DNA polymerase | -0.42 | 0.000 |
| **Rv0989c** | grcC2 | Probable polyprenyl-diphosphate synthase GrcC2 (polyprenyl pyrophosphate synthetase) | -0.44 | 0.025 |
| **Rv3726** | Rv3726 | Possible dehydrogenase | -0.51 | 0.007 |
| **L_03517** | L_03517 | hypothetical protein | -0.52 | 0.035 |
| **Rv0146** | Rv0146 | Possible S-adenosylmethionine-dependent methyltransferase | -0.52 | 0.000 |
| **Rv1273c** | Rv1273c | Probable drugs-transport transmembrane ATP-binding protein ABC transporter | -0.55 | 0.000 |
| **Rv2851c** | Rv2851c | GCN5-related N-acetyltransferase | -0.63 | 0.007 |
| **Rv0400c** | fadE7 | Acyl-CoA dehydrogenase FadE7 | -0.67 | 0.020 |
| **Rv1421** | Rv1421 | conserved protein | -0.68 | 0.000 |
| **Rv1272c** | Rv1272c | Probable drugs-transport transmembrane ATP-binding protein ABC transporter | -0.76 | 0.000 |
| **Rv2606c** | snzP | Possible pyridoxine biosynthesis protein SnzP | -0.91 | 0.000 |
| **Rv0153c** | ptbB | Phosphotyrosine protein phosphatase PTPB (protein-tyrosine-phosphatase) (PTPase) | -1.1 | 0.000 |
| **Rv1287** | Rv1287 | Conserved hypothetical protein | -1.15 | 0.000 |
| **Rv0805** | Rv0805 | Class III cyclic nucleotide phosphodiesterase (cNMP PDE) | -1.16 | 0.000 |
| **Rv0410c** | pknG | Serine/threonine-protein kinase PknG (protein kinase G) (STPK G) | -1.29 | 0.000 |
| **Rv1248c** | Rv1248c | Multifunctional alpha-ketoglutarate metabolic enzyme | -1.73 | 0.000 |

**Table S2. Transposon insertions in genes possibly generally altering mycobacterial fitness of *M. tuberculosis* H37Ra cells during KSK-106 treatment.** Negative $log_2FC$ values indicate underrepresentation, positive values overrepresentation of the mutants in the pool, demonstrating aggravating or alleviating effects of insertions, respectively.

| #Orf | Name | Description | $log_2FC$ | Adj. p-value |
|---|---|---|---|---|
| **Genes involved in drug efflux** | | | | |
| Rv1272c | *Rv1272c* | Probable drugs-transport transmembrane ATP-binding protein ABC transporter | -0.76 | 0.000 |
| Rv1273c | *Rv1273c* | Probable drugs-transport transmembrane ATP-binding protein ABC transporter | -0.55 | 0.000 |
| Rv2936 | *drrA* | Daunorubicin-dim-transport ATP-binding protein ABC transporter DrrA | -0.25 | 0.000 |
| Rv2937 | *drrB* | Daunorubicin-dim-transport integral membrane protein ABC transporter DrrB | -0.31 | 0.007 |
| Rv2942 | *mmpL7* | Conserved transmembrane transport protein MmpL7 | -0.39 | 0.000 |
| **Genes involved in cell wall assembly and integrity** | | | | |
| Rv1195 | *PE13* | PE family protein PE13 | 0.24 | 0.025 |
| Rv1808 | *PPE32* | PPE family protein PPE32 | 0.21 | 0.000 |
| Rv1809 | *PPE33* | PPE family protein PPE33 | 0.24 | 0.020 |
| Rv1040c | *PE8* | PE family protein PE8 | -0.41 | 0.035 |
| Rv2931 | *ppsA* | Phenolpthiocerol synthesis type-I polyketide synthase PpsA | 0.46 | 0.000 |
| Rv2932 | *ppsB* | Phenolpthiocerol synthesis type-I polyketide synthase PpsB | 0.59 | 0.000 |
| Rv2933 | *ppsC* | Phenolpthiocerol synthesis type-I polyketide synthase PpsC | 0.51 | 0.007 |
| Rv2935 | *ppsE* | Phenolpthiocerol synthesis type-I polyketide synthase PpsE | 0.57 | 0.000 |
| **Genes associated with universal stress protein family** | | | | |
| Rv3134c | *Rv3134c* | Universal stress protein family protein | -0.42 | 0.000 |
| **Metabolic genes** | | | | |
| Rv3696c | glpK | Probable glycerol kinase GlpK (ATP:glycerol 3-phosphotransferase) (glycerokinase) (GK) | 0.91 | 0.000 |
| Rv1328 | glgP | Probable glycogen phosphorylase GlgP | 0.41 | 0.000 |
| Rv1213 | glgC | Glucose-1-phosphate adenylyltransferase GlgC (ADP-glucose synthase) (ADP-glucose pyrophosphorylase) | 0.35 | 0.000 |

**Table S3. Transposon insertions in genes altering mycobacterial fitness of *M. tuberculosis* H37Ra cells specific in response to KSK-106 treatment.** Negative $log_2FC$ values indicate underrepresentation, positive values overrepresentation of the mutants in the pool, demonstrating aggravating or alleviating effects of insertions, respectively.

| #Orf | Name | Description | $log_2FC$ | Adj. p-value |
|---|---|---|---|---|
| **Genes involved in KSK-resistance and activation** | | | | |
| **Rv2888c** | *amiC* | Probable amidase AmiC (aminohydrolase) | 1 | 0.000 |
| **Rv0552** | *Rv0552* | Conserved protein/putative endodeoxyribonuclease | 0.94 | 0.000 |
| ***Gene involved in the oxidative stress network*** | | | | |
| **Rv3005c** | *doxX* | Probable membrane oxidoreductase component (MRC) DoxX | 0.55 | 0.000 |
| ***Genes involved in pyridoxal-5'-phosphate pathway*** | | | | |
| **Rv2606c** | *snzP* | Possible pyridoxine biosynthesis protein SnzP | -0.91 | 0.000 |
| **Rv2607** | *pdxH* | Probable pyridoxamine 5'-phosphate oxidase PdxH (PNP/PMP oxidase) (pyridoxinephosphate oxidase) (PNPOX) (pyridoxine 5'-phosphate oxidase) | -0.36 | 0.040 |

**Table S4. Strains used in this study.**

| Strain | Relevant properties | Origin |
|---|---|---|
| *Mycobacterium tuberculosis* H37Rv wild type | Wild type | William R. Jacobs Jr., PhD, Albert Einstein College of Medicine, Bronx, USA |
| H37Rv pBEN::mCherry (Hsp60)/GFP (Atc) | $hyg^R$, constitutive expression of mCherry | Reference [1] |
| H37Rv pMV361::*amiC* | Merodiploid *amiC* strain for overexpression and complementation, $kan^R$ | This study |
| H37Rv pMV361::*Rv0552* | Merodiploid R*v0552* strain, for overexpression and complementation $kan^R$ | This study |
| H37Rv pMV361::EV | Empty vector control strain, $kan^R$ | This study |
| H37Rv Δ*amiC* | Gene deletion mutant of *amiC*, $hyg^R$ | This study |
| H37Rv Δ*amiC* pMV361::*amiC* | Gene deletion mutant of *amiC*, complemented with a wild type copy of *amiC*, constitutively expressed from a single-copy integrative plasmid, $hyg^R$, $kan^R$ | This study |
| H37Rv Δ*amiC* pMV361::EV | Gene deletion mutant of amiC, complemented with a empty vector control plasmid, $hyg^R$, $kan^R$ | This study |
| H37Rv Δ*amiC* pMV361::*amiC* P185T | Gene deletion mutant of *amiC*, complemented with a mutated copy of *amiC*, constitutively expressed from a single-copy integrative plasmid, $hyg^R$, $kan^R$ | This study |
| H37Rv Δ*amiC* pMV361::*amiC* Ins 100 +t | Gene deletion mutant of amiC, complemented with a mutated copy of amiC, constitutively expressed from a single-copy integrative plasmid, $hyg^R$, $kan^R$ | This study |
| H37Rv Δ*amiC* pMV361::R*v0552* | Gene deletion mutant of amiC, complemented with a wild type copy of *Rv0552*, constitutively expressed from a single-copy integrative plasmid, $hyg^R$, $kan^R$ | This study |
| H37Rv Δ*amiC* pMV361::*amiC* K82A_S157A_S181A | Gene deletion mutant of amiC, complemented with a mutated copy of amiC, constitutively expressed from a single-copy integrative plasmid, $hyg^R$, $kan^R$ | This study |
| H37Rv Δ*Rv0552* | Gene deletion mutant of Rv0552, $hyg^R$ | This study |
| H37Rv *ΔRv0552* pMV361::*Rv0552* | Gene deletion mutant of *Rv0552*, complemented with a wild type copy of *Rv0552*, constitutively expressed from a single-copy integrative plasmid, $hyg^R$, $kan^R$ | This study |
| H37Rv Δ*Rv0552* pMV361::EV | Gene deletion mutant of Rv0552, complemented with a empty vector control plasmid, $hyg^R$, $kan^R$ | This study |

| | | |
|---|---|---|
| H37Rv *ΔRv0552* pMV361::*Rv0552* H67R | Gene deletion mutant of Rv0552, complemented with a mutated copy of Rv0552, constitutively expressed from a single-copy integrative plasmid, *hyg*$^R$, *kan*$^R$ | This study |
| H37Rv *ΔRv0552* pMV361::*Rv0552* A229D | Gene deletion mutant of *Rv0552*, complemented with a mutated copy of *Rv0552*, constitutively expressed from a single-copy integrative plasmid, *hyg*$^R$, *kan*$^R$ | This study |
| H37Rv *ΔRv0552* pMV361::*amiC* | Gene deletion mutant of *Rv0552*, complemented with a wild type copy of *amiC*, constitutively expressed from a single-copy integrative plasmid, *hyg*$^R$, *kan*$^R$ | This study |
| H37Rv Δ*Rv3092c-Rv3095* | Deletion mutant of gene cluster *Rv3092c-Rv3095*, *hyg*$^R$ | This study |
| *M. tuberculosis strain* mc$^2$6030 | Auxotrophic mutant strain generated from H37Rv lacking RD1 region and *panCD* genes | obtained from William, R. Jacobs Jr., PhD, Albert Einstein College of Medicine, Bronx, USA; reference [2] |
| *M. tuberculosis* strain H37Ra | Avirulent strain | obtained from William, R. Jacobs Jr., PhD, Albert Einstein College of Medicine, Bronx, USA; |
| *M. tuberculosis* CDC1551 | Virulent lab strain | obtained from William, R. Jacobs Jr., PhD, Albert Einstein College of Medicine, Bronx, USA |
| *M. tuberculosis* Erdman | Virulent lab strain | obtained from William, R. Jacobs Jr., PhD, Albert Einstein College of Medicine, Bronx, USA |
| *M. tuberculosis* KZN06<br>*M. tuberculosis* KZN07<br>*M. tuberculosis* KZN13<br>*M. tuberculosis* KZN14<br>*M. tuberculosis* KZN15<br>*M. tuberculosis* KZN16 | clinical isolates from KZN, South Africa | obtained from William R. Jacobs Jr., PhD, Albert Einstein College of Medicine, Bronx, USA |
| *M. smegmatis* mc$^2$155 | Wild type | obtained from William R. Jacobs Jr., PhD, Albert Einstein College of Medicine, Bronx, USA |
| *M. smegmatis* pMV361::*amiC* | Merodiploid strain for overexpression of *amiC*, kan$^R$ | This study |
| *M. smegmatis* pMV361::*Rv0552* | Merodiploid strain, for overexpression of *Rv0552*, kan$^R$ | This study |
| *M. smegmatis* pMV361::EV | Empty vector control strain, *kan*$^R$ | This study |
| *M. bovis* BCG Pasteur | Wild type | obtained from William R. Jacobs Jr., PhD, Albert Einstein College of Medicine, Bronx, USA |

| | | |
|---|---|---|
| *M. bovis* pMV361::*amiC* | Merodiploid strain for overexpression of *amiC*, $kan^R$ | This study |
| *M. bovis* pMV361::*Rv0552* | Merodiploid strain, for overexpression of *Rv0552*, $kan^R$ | This study |
| *M. bovis* pMV361::EV | Empty vector control strain, *$kan^R$* | This study |
| *M. marinum* ATCC 927 | Wild type | DSMZ-German Collection of Microorganisms and Cell Cultures GmbH |
| *M. abscessus* (CF001s) | | Clinical isolate; reference [3] |
| *E. coli* NEB 5-alpha | Cloning strain for plasmids | New England Biolabs (Cat.-No. C2987I) |
| *E. coli* HB101 | Cloning strain for phasmids | Promega (Cat.-No. L2015) |

### Table S5. Oligonucleotides used in this study.

| Oligonucleotide | Sequence [5‘ – 3‘] |
|---|---|
| HindIII *amiC* | GTCGGAAGCTTCTACTCGGCGATATTTGGGG |
| PacI *amic* | GGAGGATTAATTAAATZTCGCGCGTACAGCTTC |
| HindIII *rv0552* | GGGAAGCTTTCACCGCCGATAAACCTGGC |
| PacI *rv0552* | GGAGGTTAATTAAATGGCCGATGCAGACCTCGTC |
| 5‘ *Rv0552* H67R | GCGGATGGCCGCGTGCCTCAACA |
| 3‘ *Rv0552* H67R | TACTGGAGGCGGTCGTGC |
| 5‘ *Rv0552* A229D | CTGGGTCAAAGTCCATCTCCGAGCATGTG |
| 3‘ *Rv0552* A229D | GGTATCGGCCGATGGTCG |
| 5‘ *amiC* Ins 100/473 | ATTCCCACGCGTCGGTGCC |
| 3‘ *amiC* 100/473 | CCATACGCGGCCGTCGCC |
| 5‘ *amiC* P185T | TGCAGGCGGCtgtAATACGGATCG |
| 3‘ *amiC* P185T | ACGGGTTGGTCGGGCTCA |
| 5' *amiC* K82A | GACCTTCATCGCAGACAACGTCGACG |
| 3' *amiC* K82A | GGCACTCCACTGAAGAAC |
| 5' *amiC* S157A | AGCGGGTGCCGCATCATCGGGAT |
| 3' *amiC* S157A | GTGTAGTCGGTATTCCACGGATTAC |
| 5' *amiC* S181A | CGGCGGCGGCGCAATCCGTATTC |
| 3' *amiC* S181A | TCGTTGGCGTGCGCGATC |
| *amic* LL | TTTCTTTTCACAAAGTGGGACTGGGGCGCGCCGACGG |
| *amiC* LR | TTTTGTCTCACTTCGTGCATACCCGGCTAAGCCTGGC |
| *amiC* RL | TTGTCTTTGCATAGATTGCTAGCCCGCGTCCGGATCTAG |
| *amiC* RR | TCCGTGGTGCATCTTTTGCCGACTTCAGTAACGACCTTG |
| *Rv0552* LL | TTTTCCATAAATTGGGCGCCGTAGTAGACGGTTTC |
| *Rv0552* LR | AAAAAAACCATTTCTTGGCATGGGCTCCAAGCGTAGTG |
| *Rv0552* RR | TTTTTTTTCCATCTTTTGGCCAGGTTGATCACCCGAAAG |
| *Rv0552* RL | GGGTTTTTCCATAGATTGGGTGATACCCGTGCTGCCCCC |
| 5′ *amiC* confirmation | ATCTAAGCGCCGTGACGGTCCAG |
| 3' *amiC* confirmation | TGAAAGAGACCGTCGCTCAC |
| 5' *rv0552* confirmation | ATTCTGAGCAACCGCCGGATG |
| 3' *rv0552* confirmation | AGCAATCTAGCTGACAGCGCAGTCCG |
| *Hyg* confirmation | GCACGGGACCAACATCTTCG |
| *sacB* confirmation | TTTGTAATGGCCAGCTGTCC |
| 3' *Rv3096* confirmation | TGTCTCGCATTCGGCGAGC |
| 5' *Rv3091* confirmation | TATGGGCGCGGGCTTCGTCTACA |
| Operon-*Rv3095* LL | TTGCGTTCTCAGAAACTGGACCAGTTCGCCCATACCCTG |
| Operon-*Rv3095* LR | TTGGCTTTCAGTTCCTGCCGTCGCTAGGTCAGGTGGC |
| Operon-*Rv3095* RL | TGCGCTCTCAGAGACTGCGATTCCCAACCTCAAATTG |
| Operon-*Rv3095* RR | TCTTGTTTCAGCTTCTGTTGGGACCGCGCCAGGTAC |

| | |
|---|---|
| 5' *amiC* (PCR) | GGCTCGTGTGGCGATTTTCGAC |
| 3' *amiC* (PCR) | TTAATGACGCCCCGCTGGGCTATC |
| 5' *Rv0552* (PCR) | ATAGCACCGTTGGCGTCCACCCGCACCAT |
| 3' *Rv0552* (PCR) | GGCGCATCAAAACTTCAGGACGGTTGAG |
| 5' *amiC* (seq) | AATTATGTCGAGGCCGCCATCGCCCG |
| 3' *amiC* (seq) | GAGAGCATCATGCCCACGG |
| 5' *Rv0552* (seq) | TTATCGTCAGGCGCTCCTCCGGTG |
| 3' *Rv0552* (seq) | TTATTGGAGGCGGTCGTGCTGTCGG |
| PacI *frr* | GCGTTAATTAAAGCGATGAGGAGGAGCGGCGCAG |
| HindIII_*frr* | GCGAAGCTTTGTCACGGATTTTGTTGCTGAGCG |
| PacI *Rv0812* | CGCGGTTAATTAAGGGACATGTTGAGGCAGACG |
| HindIII *Rv0812* | GAGAAGCTTGCGCTGACCACAACAGAGG |
| 5' NdeI *amiC*- 6xHis-Tag | GGGATGCATATGCACCACCATCACCATCACATGTCGCGCGTACACGC |
| 3' DraIII *amiC* | GGGCACTACGTGCTACTCGGCGATATTTGGG |

**Table S6. Plasmids and phasmids used in this study.**

| Plasmid (p) or Phasmid (ph) | Relevant properties |
|---|---|
| p0004S | *sacB*-hyg$^R$ cassette, oriE-*cos* site cassette; reference [4] |
| phAE159 | Temperature sensitive shuttle phasmid, derivative of mycobacteriophage TM4, amp$^R$; reference [4] |
| pMV361(kan)::EV | Integrative *E. coli*-mycobacteria shuttle palsmid, kan$^R$; reference [5] |
| pMV361(kan)::*Rv0552* | complementation and overexpression plasmid, kan$^R$ |
| pMV361(kan)::*amiC* | complementation and overexpression plasmid, kan$^R$ |
| p0004s-*amiC*-k.o. | Knock-out cassette, hyg$^R$, sacB, flanking regions of *amiC,* hyg$^R$ |
| p0004s-R*v0552*-k.o. | Knock-out cassette, hyg$^R$, sacB, flanking regions of *Rv0552,* hyg$^R$ |
| p0004s - operon rv3095- k.o. | Knock-out cassette, hyg$^R$, sacB, flanking regions of *Rv3092c-Rv3095,* hyg$^R$ |
| phAE159::*amiC*-k.o | Knock-out cassette, hyg$^R$, sacB, flanking regions of *amiC,* hyg$^R$ |
| phAE159::*Rv0552*-k.o. | Knock-out cassette, hyg$^R$, sacB, flanking regions of *Rv0552,* hyg$^R$ |
| phAE159::operon-*Rv3095*-k.o. | Knock-out cassette, hyg$^R$, sacB, flanking regions of *Rv3092c-Rv3095,* hyg$^R$ |
| pMV361::*amiC* P185T | complementation plasmid, kan$^R$ |
| pMV361::*amiC* Ins100/473+t | complementation plasmid, kan$^R$ |
| pMV361::*Rv0552* H67R | complementation plasmid, kan$^R$ |
| pMV361::*Rv0552* A229D | complementation plasmid, kan$^R$ |
| pMV361::*amiC*_K82A_S157A_S181A | complementation plasmid, kan$^R$ |

### Synthesis of starting materials and intermediates

#### 4-(Heptyloxy)benzoic acid (6c)

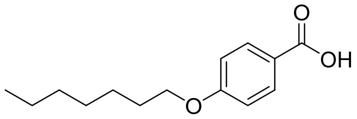

The preparation of **6c** was carried out according to a procedure of CHENG *et al.*[6] 4-(Heptyloxy)benzoic acid (**6c**) was obtained as a colorless solid (62 %, 2 steps). The spectroscopic data correspond to those reported in the literature.

**$^1$H-NMR (300 MHz, DMSO-*d*$_6$)** δ 7.93 – 7.80 (m, 2H), 7.05 – 6.93 (m, 2H), 4.02 (t, *J* = 6.5 Hz, 2H), 1.79 – 1.64 (m, 2H), 1.48 – 1.19 (m, 8H), 0.94 – 0.79 (m, 3H)

**HPLC** $t_R$ = 16.32 min, purity ≥ 99.9 %

#### 4-(Thiazol-2-yl)benzoic acid (6d)

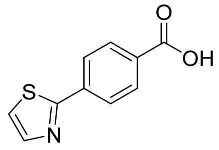

The preparation of **6d** was carried out according to a procedure of TANI *et al.*[7] 4-(Thiazol-2-yl)benzoic acid (**6d**) was obtained as a colorless solid (42 %, 3 steps). The spectroscopic data correspond to those reported in the literature.

**$^1$H-NMR (300 MHz, DMSO-*d*$_6$)** δ 13.15 (s, 1H), 8.12 – 8.02 (m, 4H), 8.01 (d, *J* = 3.2 Hz, 1H), 7.90 (d, *J* = 3.2 Hz, 1H)

**HPLC** $t_R$ = 9.05 min, purity ≥ 99.9 %

#### *N*-hydroxy-[1,1'-biphenyl]-4-carboxamide (12a)

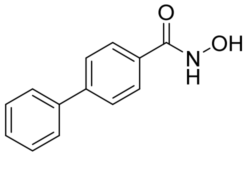

To a solution of biphenyl-4-carboxylic acid (**6a**) (50.0 mmol, 1.00 eq.) and *N,N*-dimethylformamide (50.0 mmol, 1.00 eq.) in 150 mL anhydrous dichloromethane oxalyl chloride (113 mmol, 2.25 eq.) was slowly added under ice cooling. The reaction mixture was stirred for 1 h at 0 °C and then slowly added to a solution of hydroxylamine hydrochloride (200 mmol, 4.00 eq.) and triethylamine (300 mmol, 6.00 eq.) in 90.0 mL tetrahydrofuran / water (5:1). The suspension was stirred for 16 h at room temperature, the pH was adjusted to 6 with 2 M hydrochloric acid solution and the mixture was extracted four times with 200 mL dichloromethane. The combined organic phases were dried over anhydrous sodium sulfate, filterd and the solvent was solvent was removed under reduced pressure. The crude product was recrystallized from water/ethanol and *N*-hydroxy-[1,1'-biphenyl]-4-carboxamide (**12a**) was obtained as white needles (87 % yield). The spectroscopic data agree with those reported in the literature.[8]

**$^{1}$H-NMR (300 MHz, DMSO-d$_6$)** δ 7.36 – 7.44 (m, 1H), 7.45 – 7.54 (m, 2H), 7.68 – 7.79 (m, 4H), 7.82 – 7.89 (m, 2H), 9.06 (s, 1H), 11.28 (s, 1H)
**$^{13}$C-NMR (75 MHz, DMSO)** δ 126.6, 126.8, 127.5, 128.0, 129.0, 131.5, 139.1, 142.7, 163.8
**HPLC** $t_R$ = 9.08 min, purity ≥ 99.9 %
**Mp.** $T_M$ = 197.6 °C

***N*-hydroxy-4-(pentyloxy)benzamide (12b)**

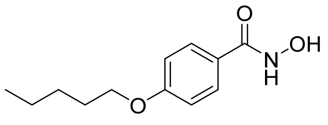

To a solution of 4-(pentyloxy)benzoic acid (**6b**) (50.0 mmol, 1.00 eq.) and *N,N*-dimethylformamide (50.0 mmol, 1.00 eq.) in 150 mL anhydrous dichloromethane oxalyl chloride (113 mmol, 2.25 eq.) was slowly added under ice cooling. The reaction mixture was stirred for 1 h at 0 °C and then slowly added to a solution of hydroxylamine hydrochloride (200 mmol, 4.00 eq.) and triethylamine (300 mmol, 6.00 eq.) in 100 mL tetrahydrofuran/ water (5:1). The suspension was stirred for 16 h at room temperature, the pH was adjusted to 6 with 2 M hydrochloric acid solution and the mixture was extracted four times with 200 mL dichloromethane. The combined organic phases were dried over anhydrous sodium sulfate, filtered and the solvent was removed under reduced pressure. The crude product was recrystallized from water / ethanol and *N*-hydroxy-4-(pentyloxy)benzamide and (**12b**) was obtained as white needles (73 % yield). The spectroscopic data agree with those reported in the literature.[8]
**$^{1}$H-NMR (300 MHz, DMSO-*d$_6$*)** δ 11.04 (s, 1H), 8.88 (s, 1H), 7.88 – 7.57 (m, 2H), 7.09 – 6.81 (m, 2H), 4.00 (t, *J* = 6.5 Hz, 2H), 1.83 – 1.62 (m, 2H), 1.49 – 1.23 (m, 4H), 0.99 – 0.79 (m, 3H)
**$^{13}$C-NMR (75 MHz, DMSO-*d$_6$*)** δ 164.02, 160.92, 128.61, 124.74, 114.02, 67.61, 28.26, 27.66, 21.87, 13.91
**HPLC** $t_R$ = 10.81 min, purity = 98.8 %
**Mp.** $T_M$ = 150 °C

**2-([1,1'-biphenyl]-4-carboxamidooxy)acetic acid** (**10**)[9]

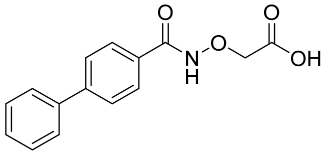

A solution of *N*-hydroxy-[1,1'-biphenyl]-4-carboxamide (**12a**, 34.6 mmol, 1.00 eq.), bromoacetic acid (**1a**, 34.6 mmol, 1.00 eq.) and sodium hydroxide (69.2 mmol, 2.00 eq.) in 200 mL ethanol was heated to reflux for 8 h. After cooling to room temperature, the solvent was removed under reduced pressure and the residue was dissolved in 200 mL distilled water and the pH was adjusted with 2 M hydrochloric acid solution to 4. The aqueous suspension was extracted four times with 200 mL dichloromethane, the combined organic phases were dried over anhydrous sodium sulfate, filtered and the solvent was removed under reduced pressure. The crude product was recrystallized from ethyl acetate and 2-([1,1'-biphenyl]-4-carboxamidooxy)acetic acid (**10**) was obtained as white needles (80 % yield).

**$^1$H-NMR (300 MHz, DMSO-d$_6$)** δ 4.53 (s, 2H), 7.37 – 7.45 (m, 1H), 7.45 – 7.55 (m, 2H), 7.68 – 7.74 (m, 2H), 7.74 – 7.81 (m, 2H), 7.82 – 7.92 (m, 2H), 12.07 (s, 1H), 12.99 (s, 1H)

**$^{13}$C-NMR (75 MHz, DMSO)** δ 71.8, 126.6, 126.8, 127.9, 128.1, 129.0, 130.5, 139.0, 143.2, 164.6, 170.0

**HPLC** $t_R$ = 10.23 min, purity = 98.0 %

**Mp.** $T_M$ = 191.8 °C

**2-((4-(pentyloxy)benzamido)oxy)acetic acid (11)**

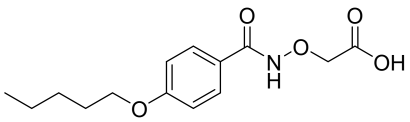


A solution of *N*-hydroxy-4-(pentyloxy)benzamide (**12b**, 10.0 mmol, 1.00 eq.), bromoacetic acid (**1a**, 10.0 mmol, 1.00 eq.) and sodium hydroxide (20.0 mmol, 2.00 eq.) in 40.0 mL ethanol was heated to reflux for 8 h. After cooling to room temperature, the solvent was removed under reduced pressure and the residue was dissolved in 50.0 mL distilled water and the pH was adjusted with 2 M hydrochloric acid solution to 4. The aqueous suspension was extracted four times with 40.0 mL dichloromethane, the combined organic phases were dried over anhydrous sodium sulfate, filtered and the solvent was removed under reduced pressure. The crude product was recrystallized from ethyl acetate and 2-((4-(pentyloxy)benzamido)oxy)acetic acid (**11**) was obtained as white needles (51 % yield).

**$^1$H-NMR (300 MHz, DMSO-*d$_6$*)** δ 13.00 (s, 1H), 11.84 (s, 1H), 7.85 – 7.62 (m, 2H), 7.06 – 6.86 (m, 2H), 4.48 (s, 2H), 4.01 (t, *J* = 6.5 Hz, 2H), 1.81 – 1.62 (m, 2H), 1.36 (tdt, *J* = 8.0, 3.6, 2.2 Hz, 4H), 0.97 – 0.81 (m, 3H)

**$^{13}$C-NMR (75 MHz, DMSO-*d$_6$*)** δ 170.12, 164.85, 161.49, 129.12, 123.53, 114.12, 71.97, 67.68, 28.23, 27.63, 21.86, 13.90

**HPLC** $t_R$ = 11.91 min, purity = 99.3 %

**Mp.** $T_M$ = 162.6 °C

**2-([1,1'-biphenyl]-4-carboxamidooxy)propanoic acid (13)**

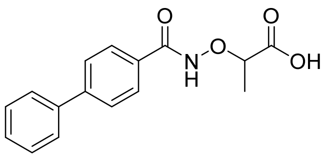


Under nitrogen atmosphere at -10 °C, to a solution of *N*-hydroxy-[1,1'-biphenyl]-4-carboxamide (**12a**, 4.69 mmol, 1.00 eq.) in 20.0 mL anhydrous tetrahydrofuran sodium hydride (60% dispersion in mineral oil, 5.16 mmol, 1.10 eq.) was slowly added and the reaction was stirred for 10 min. 2-bromopropanoic acid (**1b**, 4.69 mmol, 1.00 eq.) dissolved in 5.00 mL absolute tetrahydrofuran was added dropwise. The reaction mixture was heated to reflux for 24 h. After cooling down to room temperature, the solvent was removed under reduced pressure and the residue was dissolved in 50.0 mL distilled water and the pH was adjusted with a 2 M hydrochloric acid solution to 1. The aqueous phase was extracted three times with 50.0 mL ethyl acetate, the combined organic phases were dried over anhydrous sodium sulfate, filtered and the solvent was removed under reduced pressure. The crude product was purified by flash chromatography on silica gel using *n*-hexane/ethyl acetate as eluents. 2-([1,1'-Biphenyl]-4-carboxamidooxy)propanoic acid (**13**) was obtained as a light yellow solid (70 % yield).

**$^1$H-NMR (600 MHz, DMSO-d$_6$)** δ 1.41 (d, J = 7.0 Hz, 3H), 4.54 (q, J = 6.9 Hz, 1H), 7.38 – 7.45 (m, 1H), 7.49 (dd, J = 8.4, 7.0 Hz, 2H), 7.69 – 7.75 (m, 2H), 7.74 – 7.80 (m, 2H), 7.83 – 7.89 (m, 2H), 11.94 (br s, 1H), 12.95 (br s, 1H)

**$^{13}$C-NMR (151 MHz, DMSO)** δ 16.9, 79.1, 127.1, 127.3, 128.4, 128.6, 129.5, 131.0, 139.5, 143.7, 165.3, 173.3

**HPLC** $t_R$ = 12.22 min, purity = 96.8 %

**Mp.** $T_M$ = 173.8 °C

**General procedure 2 (GP 2): Synthesis of hydroxylamines[10]**

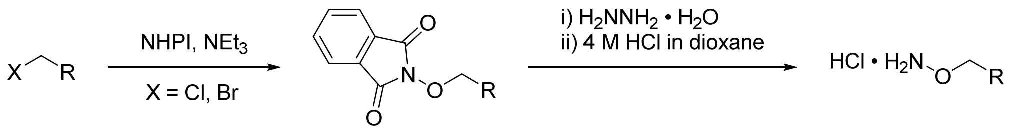


The respective alkyl halide (1.00 eq.), *N*-hydroxyphthalimide (NHPI, 1.20 eq.) and triethylamine (2.00 eq.) were heated in (10.0 mL/mmol) acetonitrile or *N,N*-dimethylformamide until complete conversion to reflux. After cooling down to room temperature, the solvent was removed under reduced pressure, the residue was dissolved in (10.0 mL/mmol) ethyl acetate and washed exhaustively with saturated sodium hydrogen carbonate solution. The organic phase was dried over anhydrous sodium sulfate, filtered and the solvent was removed under reduced pressure. The phthaloyl-protected hydroxylamines were used directly without further purification for deprotection using method A or B.

**Deprotection method A:**

The corresponding phthaloyl-protected hydroxylamine (1.00 eq.) was dissolved in (15.0 mL/mmol) ethanol and hydrazine monohydrate (2.00 eq.) was added. The reaction mixture was heated to reflux for 2 h and after cooling down to room temperature it was stored for 1 h at 0 °C. The precipitate was isolated by filtration and washed twice with ice-cold ethanol. The filtrate was cooled to 0 °C and the pH was adjusted to 2 - 3 with 4 M hydrochloric acid in 1,4-dioxane while stirring continuously. The precipitated hydroxylamine hydrochloride was isolated by filtration and washed with ice-cold 1,4-dioxane.

**Deprotection method B:**

The corresponding phthaloyl-protected hydroxylamine (1.00 eq.) was dissolved in dichloromethane or tetrahydrofuran (15.0 mL/mmol) and hydrazine monohydrate (2.00 eq.) was added. The reaction mixture was stirred for 4 h at room temperature, then stored overnight at 7 °C and the precipitate was removed by filtration. The filtrate was diluted three times with saturated sodium hydrogen carbonate solution (5.00 mL/mmol) and once with saturated sodium chloride solution (5.00 mL/mmol). The organic phase was dried over anhydrous sodium sulfate, filtered and the solvent removed under reduced pressure. The residue was dissolved in absolute diethyl ether (5.00 mL/mmol) and the pH was adjusted to 1 with 4 M hydrochloric acid in 1,4-dioxane under ice cooling. The suspension was allowed to stand overnight at -20 °C, the precipitated hydroxylamine hydrochloride was isolated by filtration and washed with ice-cold dry diethyl ether.

***O*-benzylhydroxylamine hydrochloride (2)**

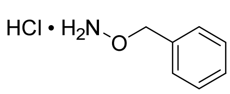


**2** is also commercially available and was used for **GP 1** without further purification.

***O*-(tert-butyl)hydroxylamine hydrochloride (14a)**

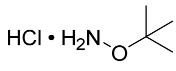


**14a** also is commercially available and was used for **GP 1** without further purification.

***O*-hexylhydroxylamine hydrochloride (14b)[11]**

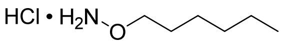


The synthesis was carried out according to **GP 2** (method A). *O*-hexylhydroxylamine hydrochloride (**14b**) was obtained as a colorless solid (55 %).

**$^1$H-NMR (300 MHz, DMSO-d$_6$)** δ 0.79 – 0.94 (m, 3H), 1.13 – 1.38 (m, 6H), 1.57 (tt, *J* = 7.4, 6.1 Hz, 2H), 3.99 (t, *J* = 6.5 Hz, 2H), 10.99 (br s, 3H).

**Mp**.: 151.3 °C

***O*-phenylhydroxylamine hydrochloride (14c)[12]**

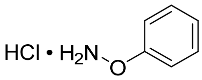


The synthesis was carried out according to **GP 2** (method B). *O*-phenylhydroxylamine hydrochloride (**14c**) was obtained as a colorless solid (80 %).

**$^1$H-NMR (600 MHz, DMSO-d$_6$)** δ = 7.06 – 7.11 (m, 1H), 7.15 – 7.21 (m, 2H), 7.35 – 7.40 (m, 2H).

**Mp**.: 129.5 °C

***O*-(cyclohexylmethyl)hydroxylamine hydrochloride (14d)[13]**

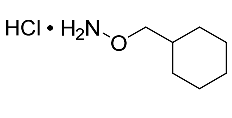


The synthesis was carried out according to **GP 2** (method B). *O*-(cyclohexylmethyl)hydroxylamine hydrochloride (**14d**) was obtained as a colorless solid (48 %).

**$^1$H-NMR (300 MHz, DMSO-d$_6$)** δ 0.83 – 1.04 (m, 2H), 1.19 (h, *J* = 12.1 Hz, 3H), 1.55 – 1.73 (m, 6H), 3.81 (d, *J* = 6.1 Hz, 2H), 10.92 (br s, 3H).

**Mp**.: 178.6 °C

#### *O*-(pyridin-2-ylmethyl)hydroxylamine dihydrochloride (14e)[14]

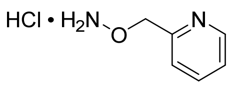

The synthesis was carried out according to **GP 2** (method B). *O*-(pyridin-2-ylmethyl)hydroxylamine dihydrochloride (**14e**) was obtained as a colorless solid (52 %).

**$^1$H-NMR (300 MHz, DMSO-d$_6$)** δ = 5.29 (s, 2H), 7.97 (ddd, *J* = 7.9, 5.5, 0.8 Hz, 1H), 8.48 (dt, *J* = 8.0, 1.8 Hz, 1H), 8.88 (dd, *J* = 5.6, 1.5 Hz, 1H), 8.94 (d, *J* = 2.0 Hz, 1H).

**Mp**.: 188.3 °C

#### References Supporting Information


1. Martin, C.J., Booty, M.G., Rosebrock, T.R., Nunes-Alves, C., Desjardins, D.M., Keren, I., Fortune, S.M., Remold, H.G., and Behar, S.M. (2012). Efferocytosis Is an Innate Antibacterial Mechanism. Cell Host & Microbe *12*, 289-300. 10.1016/j.chom.2012.06.010.
2. Sambandamurthy, V.K., Derrick, S.C., Hsu, T., Chen, B., Larsen, M.H., Jalapathy, K.V., Chen, M., Kim, J., Porcelli, S.A., Chan, J., et al. (2006). Mycobacterium tuberculosis DeltaRD1 DeltapanCD: a safe and limited replicating mutant strain that protects immunocompetent and immunocompromised mice against experimental tuberculosis. Vaccine *24*, 6309-6320. 10.1016/j.vaccine.2006.05.097.
3. Steindor, M., Nkwouano, V., Stefanski, A., Stuehler, K., Ioerger, T.R., Bogumil, D., Jacobsen, M., Mackenzie, C.R., and Kalscheuer, R. (2019). A proteomics approach for the identification of species-specific immunogenic proteins in the Mycobacterium abscessus complex. Microbes Infect *21*, 154-162. 10.1016/j.micinf.2018.10.006.
4. Jain, P., Hsu, T., Arai, M., Biermann, K., Thaler, D.S., Nguyen, A., Gonzalez, P.A., Tufariello, J.M., Kriakov, J., Chen, B., et al. (2014). Specialized transduction designed for precise high-throughput unmarked deletions in Mycobacterium tuberculosis. mBio *5*, e01245-01214. 10.1128/mBio.01245-14.
5. Jacobs, W.R., Tuckman, M., and Bloom, B.R. (1987). Introduction of Foreign DNA into Mycobacteria Using a Shuttle Phasmid. Nature *327*, 532-535. DOI 10.1038/327532a0.
6. Cheng, X.H., Bai, X.Q., Jing, S., Ebert, H., Prehm, M., and Tschierske, C. (2010). Self-Assembly of Imidazolium-Based Rodlike Ionic Liquid Crystals: Transition from Lamellar to Micellar Organization. Chem-Eur J *16*, 4588-4601. 10.1002/chem.200903210.
7. Tani, S., Uehara, T.N., Yamaguchi, J., and Itami, K. (2014). Programmed synthesis of arylthiazoles through sequential C-H couplings. Chem Sci *5*, 123-135. 10.1039/c3sc52199k.
8. Feng, C., and Loh, T.P. (2014). Rhodium-Catalyzed C-H Alkynylation of Arenes at Room Temperature. Angew Chem Int Edit *53*, 2722-2726. 10.1002/anie.201309198.
9. Mchale, D., Green, J., and Mamalis, P. (1960). Amino-Oxy-Derivatives .1. Some Alpha-Amino-Oxy-Acids and Alpha-Amino-Oxy-Hydrazides. J Chem Soc, 225-229. DOI 10.1039/jr9600000225.
10. Pflieger, M., Hamacher, A., Öz, T., Horstick-Muche, N., Boesen, B., Schrenk, C., Kassack, M.U., and Kurz, T. (2019). Novel α,β-unsaturated hydroxamic acid

derivatives overcome cisplatin resistance. Bioorgan Med Chem *27*, 115036. 10.1016/j.bmc.2019.07.052.
11. High, A., Prior, T., Bell, R.A., and Rangachari, P.K. (1999). Probing the "active site" of diamine oxidase:: Structure-activity relations for histamine potentiation by O-alkylhydroxylamines on colonic epithelium. J Pharmacol Exp Ther *288*, 490-501.
12. Wang, M.Z., Xu, H., Liu, T.W., Feng, Q., Yu, S.J., Wang, S.H., and Li, Z.M. (2011). Design, synthesis and antifungal activities of novel pyrrole alkaloid analogs. Eur J Med Chem *46*, 1463-1472. 10.1016/j.ejmech.2011.01.031.
13. Laruelle, C., Lepant, M., and Raynier, B. (1989) Symmetrical O-substituted dioximes of benzo-fused β-diketocyclo-alkylenes, the processes for their preparation and their application as drugs. United States patent US4804684A.
14. Faolli, A.A., Bleyman, O.I., Kao, W., and Abou-Gharbia, M.A. (1996) Carbamates of rapamycin. United States patent US5550133
